# Ultra-high density electrodes improve detection, yield, and cell type identification in neuronal recordings

**DOI:** 10.1101/2023.08.23.554527

**Authors:** Zhiwen Ye, Andrew M. Shelton, Jordan R. Shaker, Julien Boussard, Jennifer Colonell, Daniel Birman, Sahar Manavi, Susu Chen, Charlie Windolf, Cole Hurwitz, Tomoyuki Namima, Federico Pedraja, Shahaf Weiss, Bogdan Raducanu, Torbjørn V. Ness, Xiaoxuan Jia, Giulia Mastroberardino, L. Federico Rossi, Matteo Carandini, Michael Häusser, Gaute T. Einevoll, Gilles Laurent, Nathaniel B. Sawtell, Wyeth Bair, Anitha Pasupathy, Carolina Mora Lopez, Barun Dutta, Liam Paninski, Joshua H. Siegle, Christof Koch, Shawn R. Olsen, Timothy D. Harris, Nicholas A. Steinmetz

**Author notes:** equal contribution (Zhiwen Ye, Andrew M. Shelton).

## Abstract

To understand the neural basis of behavior, it is essential to sensitively and accurately measure neural activity at single neuron and single spike resolution. Extracellular electrophysiology delivers this, but it has biases in the neurons it detects and it imperfectly resolves their action potentials. To minimize these limitations, we developed a silicon probe with much smaller and denser recording sites than previous designs, called Neuropixels Ultra (*NP Ultra*). This device samples neuronal activity at ultra-high spatial density (∼10 times higher than previous probes) with low noise levels, while trading off recording span. NP Ultra is effectively an implantable voltage-sensing camera that captures a planar image of a neuron’s electrical field. We use a spike sorting algorithm optimized for these probes to demonstrate that the yield of visually-responsive neurons in recordings from mouse visual cortex improves up to ∼3-fold. We show that NP Ultra can record from small neuronal structures including axons and dendrites. Recordings across multiple brain regions and four species revealed a subset of extracellular action potentials with unexpectedly small spatial spread and axon-like features. We share a large-scale dataset of these brain-wide recordings in mice as a resource for studies of neuronal biophysics. Finally, using ground-truth identification of three major inhibitory cortical cell types, we found that these cell types were discriminable with approximately 75% success, a significant improvement over lower-resolution recordings. NP Ultra improves spike sorting performance, detection of subcellular compartments, and cell type classification to enable more powerful dissection of neural circuit activity during behavior.

## Introduction

High-density silicon electrode arrays, such as Neuropixels probes, have enabled prolonged recordings of hundreds to thousands of neurons across many brain regions in a single experiment (Jun et al., 2017; Steinmetz et al., 2018, 2021). Their versatility and robustness has enabled collection of these datasets in diverse species including fish, rodents, monkeys, reptiles, and humans (Steinmetz et al., 2019; Norimoto et al., 2020; Metzen and Chacron, 2021; Siegle et al., 2021; Chung et al., 2022; Paulk et al., 2022; Trautmann et al., 2023). The use of Neuropixels has driven discoveries and insights into the nature of decision making, perception, and the brain-wide dynamics of neuronal processing (Allen et al., 2019; Musall et al., 2019; Steinmetz et al., 2019; Stringer et al., 2019; Vesuna et al., 2020; Siegle et al., 2021; Gardner et al., 2022; Jia et al., 2022; Chen et al., 2023; International Brain Laboratory et al., 2023b).

Despite these successes, key technical challenges impede progress on a number of pressing issues in neuroscience. First, the spatial resolution of previous generations of Neuropixels probes is relatively low: the nearest contact-to-contact spacing is ∼25 μm in Neuropixels 1.0 and ∼15 µm in 2.0. Accordingly, previous probe generations do not have sufficient site density to optimally sample fine-scale brain structures such as small nuclei or thin cell layers. In addition, our ability to sample electrical fields of individual neurons at a high spatial resolution has been limited. While we have advanced knowledge of electrical fields at the columnar level and beyond, such as the LFP and EEG (Buzsaki, 2006; Nunez and Srinivasan, 2006; Katzner et al., 2009; Buzsáki et al., 2012; Halnes et al., 2024), we have been unable to access the microstructure of electrical fields at the micrometer scale. Moreover, the low density of previous probes may undersample extracellular action potentials with electrical “footprints” (i.e. the detectable spatial extent of the extracellular action potential) smaller than the span between electrode contacts, and therefore create a sampling bias against such small-footprint signals. If such small-footprint signals are present in neural tissue, this bias against them would limit the yield of recordings and contribute to the ’dark matter problem’ in neuroscience (Shoham et al., 2006).

A second challenge for large-scale electrophysiology is motion of the brain relative to the probe, which can cause spikes to drift between recording sites, posing problems for accurate spike sorting (Steinmetz et al., 2021; Windolf et al., 2023). Probes with insufficient recording site density may even “lose” neurons entirely as their spikes become undetectable if the neuron’s extracellular action potential falls into the gaps between sites. Conversely, a higher density of recording sites and thus higher spatial sampling would allow for a more accurate estimate of brain motion, improved ability to record all spikes from neurons, and improved *post-hoc* ability to remove associated sorting errors from electrophysiological recordings.

Finally, the brain contains a large diversity of cell types, and extracellular electrophysiology has been limited in its ability to discriminate between these types (Gouwens et al., 2020; Yao et al., 2021). Classically, extracellularly recorded units in some brain regions such as cortex and striatum have been separated into coarse cell types on the basis of features such as waveform shape and firing pattern (McCormick et al., 1985; Barthó et al., 2004; Mitchell et al., 2007; Niell and Stryker, 2008; Yamin et al., 2013; Roux et al., 2014; Senzai and Buzsáki, 2017; Yu et al., 2019; Lee et al., 2021, 2024; Beau et al., 2024). NP 1.0 probes can provide additional coarse morphological information about the electrical field surrounding a neuron useful for neuron classification (Buccino et al., 2018; Jia et al., 2019), but this information may not be detailed enough to resolve the large diversity of cell types in the brain. The possibility that higher resolution electrode arrays could improve cell type classification remains largely unexplored.

We hypothesized that an electrophysiological probe with increased sampling density would address each of the above technical challenges by giving access to electrical field microstructures. Here we describe ‘Neuropixels Ultra’, a device with substantially smaller and denser recording sites than previous devices. We found that this technology increases spike sorting signal-to-noise ratio (SNR) and yield. While the ability of NP Ultra to track neurons stably during motion did not further improve over NP 2.0, the higher density did reduce sampling bias, revealing a population of small-footprint units in many brain regions and in multiple species. Pharmacological experiments demonstrated that small footprint units correspond to recordings from axons. Moreover, these probes improved cell type discrimination between the three major interneuron types expressing parvalbumin (PV), somatostatin (SST), and vasoactive intestinal peptide (VIP), which could be discriminated with approximately 75% accuracy, significantly better than could be achieved with NP 1.0 recordings. The advantages in yield, detection, and cell-class specificity make NP Ultra a superior probe to NP 2.0 for experiments requiring those properties or for experiments focusing on small brain regions, while NP2.0 remains superior for studies requiring large recording span. Finally, we share a dataset of thousands of neuronal waveforms recorded with NP Ultra from over a dozen brain regions and four species as a resource for understanding the biophysical basis of diverse extracellular action potentials across the brain (https://npultra.steinmetzlab.net/). NP Ultra probes open a window into the fine-grained spatio-temporal structure of electrically excitable cells in behaving animals.

## Results

### Probe design and characterization

We designed a new version of the Neuropixels probe with substantially smaller and denser recording sites, called Neuropixels Ultra (NP Ultra). The probe has 5 × 5 µm titanium nitride (TiN) recording sites on an 48 × 8 grid with 1 µm gaps (for 6 µm center-to-center spacing), densely sampling a 288 × 48 µm span of brain tissue (**Fig. 1A,B**). The site size and spacing compare favorably with Neuropixels 1.0 (NP 1.0) probes with 12 × 12 µm electrodes and 20 × 16 µm staggered spacing. The probe form-factor (shank dimensions and probe base) are identical to NP 1.0, as are mechanical characteristics. A second version of NP Ultra features switchable recording sites for multiple configurations including 96 × 4, 192 × 2, and 384 × 1 site arrangements for longer span at high density (**Supp. Fig. S1**). Like existing NP 1.0 and 2.0 probes (Jun et al., 2017; Steinmetz et al., 2021), NP Ultra records from 384 channels simultaneously. Thus, NP Ultra has substantially higher site density (1.3 sites/µm vs 0.10 and 0.13 sites/µm for 1.0 and 2.0 probes, **Fig. 1B**), at the cost of reduced recording span. This increased recording site density provides much higher spatial resolution of extracellular voltages compared to lower-density probes (**Fig. 1C,D,E**). Indeed, recording individual neurons *in vivo* with this device revealed detailed portraits of their extracellular action potentials (**Fig. 1G**).

**Figure 1.**
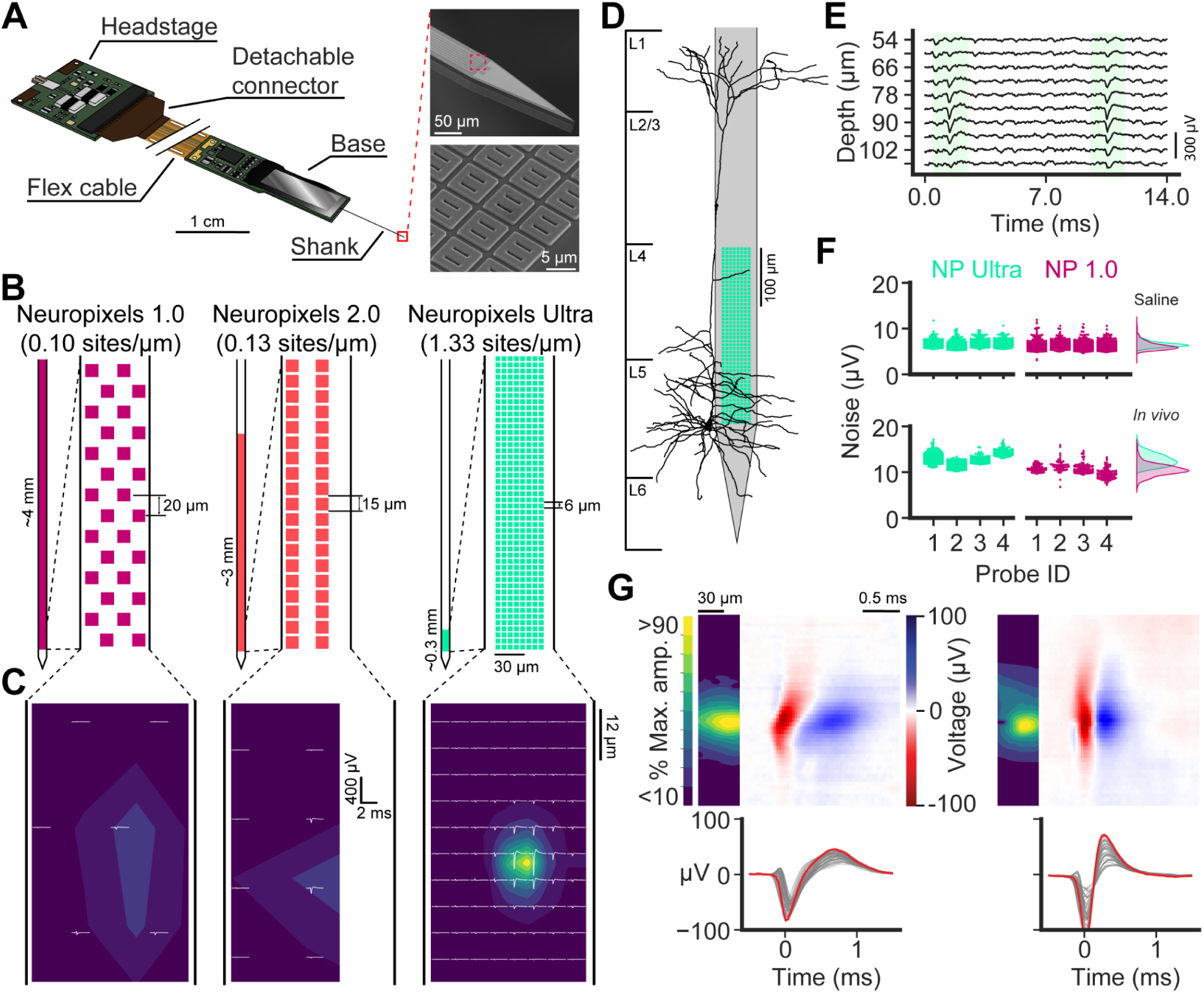
Neuropixels Ultra probes have a considerably denser site layout over a smaller spatial range compared to NP 1.0 and 2.0. **A** (left), Schematic of a complete NP Ultra probe, including headstage and flex cable. (right) Scanning electron microscope images of an NP Ultra probe tip (top) and individual contacts, each with two slits that increase electrode surface area (bottom, top inset). **B**, Layout of NP Ultra sites compared to previous probes. **C**, Comparison of an example waveform on all three probes. The NP Ultra panel (right) shows the mean waveform of a unit recorded in the mouse primary visual cortex; this waveform was spatially re-sampled to approximate the signal on the 12 × 12 µm grid for NP 1.0 (left) and NP 2.0 (middle) site configurations. Heatmaps represent the interpolated voltage as proportion of the trough-to-peak amplitude recorded on the peak amplitude channel in the NP Ultra configuration. **D**, Example reconstructed morphology of a mouse L5 pyramidal neuron from the Allen Institute Cell Types Database (ID: 488679042) compared at scale to a NP Ultra probe. **E**, Example raw traces from a column of vertically adjacent recording sites, plotted according to their depth along on an NP Ultra probe. Shading indicates the time and location of two detected spikes. **F**, Measures of noise in saline (root-mean-square) and during recording in vivo (median absolute deviation) for four NP Ultra and four NP 1.0 probes. **G**, Example spatial (top left), spatiotemporal (top right) and temporal (bottom) waveforms recorded by NP Ultra from a regular-spiking (left) and a fast-spiking (right) unit in the visual cortex of an awake mouse. Spatial plots show the contour map of channel-wise maximum amplitudes normalized to the peak channel. The spatio-temporal plots represent voltage as a colormap across all channels in the column of sites containing the peak amplitude site. The temporal plots show the peak amplitude channel (red) with 40 nearby channels (gray).

We hypothesized that smaller recording sites would improve multiple aspects of data quality, as described in the following sections, but that these would also have larger electrical impedance and, accordingly, larger noise levels (López et al., 2012). In self-referenced noise measurements of NP 1.0 and NP Ultra probes in saline, we observe a small but significant difference in RMS noise (**Fig. 1F**; 0.26 µV; 95% confidence interval from t-test = 0.19-0.34 µV). Estimating noise within tissue is complicated by the presence of true biological signals. We obtain an approximation by selecting sites with low activity levels and estimating the noise with the median absolute deviation, which is less sensitive to outliers than RMS (Rey et al., 2015). We found that noise levels *in vivo* were 20 ± 2% higher for NP Ultra than for NP 1.0 (**Fig. 1F**; difference = 2.1 µV; 95% confidence interval from t-test = 2.0-2.3 µV). The difference in noise between NP Ultra and 1.0, and the difference between saline and *in vivo* measurements, is well-captured by a noise model incorporating the properties of TiN electrodes and probe electronics, with higher impedance *in vivo* and a contribution of background neural activity (**Supp. Fig. S2;** see Methods). Though per-site noise levels are higher for these probes, we found they nevertheless have highly uniform channel-to-channel gain (**Supp. Fig. S3**), negligible crosstalk (**Supp. Fig. S4**), and no larger light sensitivity than probes with larger sites (**Supp. Fig. S5**). Despite the small site size, recordings of local field potentials (LFPs) were similar to probes with larger sites, revealing characteristic LFP features of the hippocampus (**Supp Fig. S6**).

### Higher site density improves the quality and yield of spike sorting

We reasoned that a probe with ten-fold higher site density would provide improved data quality, as it would sample individual action potentials at more recording sites. As detailed below, this increased channel density could lead to several advantages: 1) Higher peak waveform amplitudes; 2) Higher signal-to-noise ratio (SNR) for each waveform; 3) Improved spatial localization of individual action potential sources; and 4) Improved ability to stably record and track all spikes generated by a single neuron during relative motion of the probe and the brain. Before examining these four aspects of data quality, we first describe a spike sorting algorithm, DARTsort (Boussard et al., 2023), that is well suited to this dense data. We test the algorithm’s performance in challenging validation experiments, in which we recorded from mouse visual cortex and intentionally moved the probe by adjusting the position of the micromanipulator in a triangle-wave pattern for ∼500 seconds during the session (**Fig. 2**), a strategy that provides a difficult test case for the algorithm with an approximately known pattern of motion.

**Figure 2.**
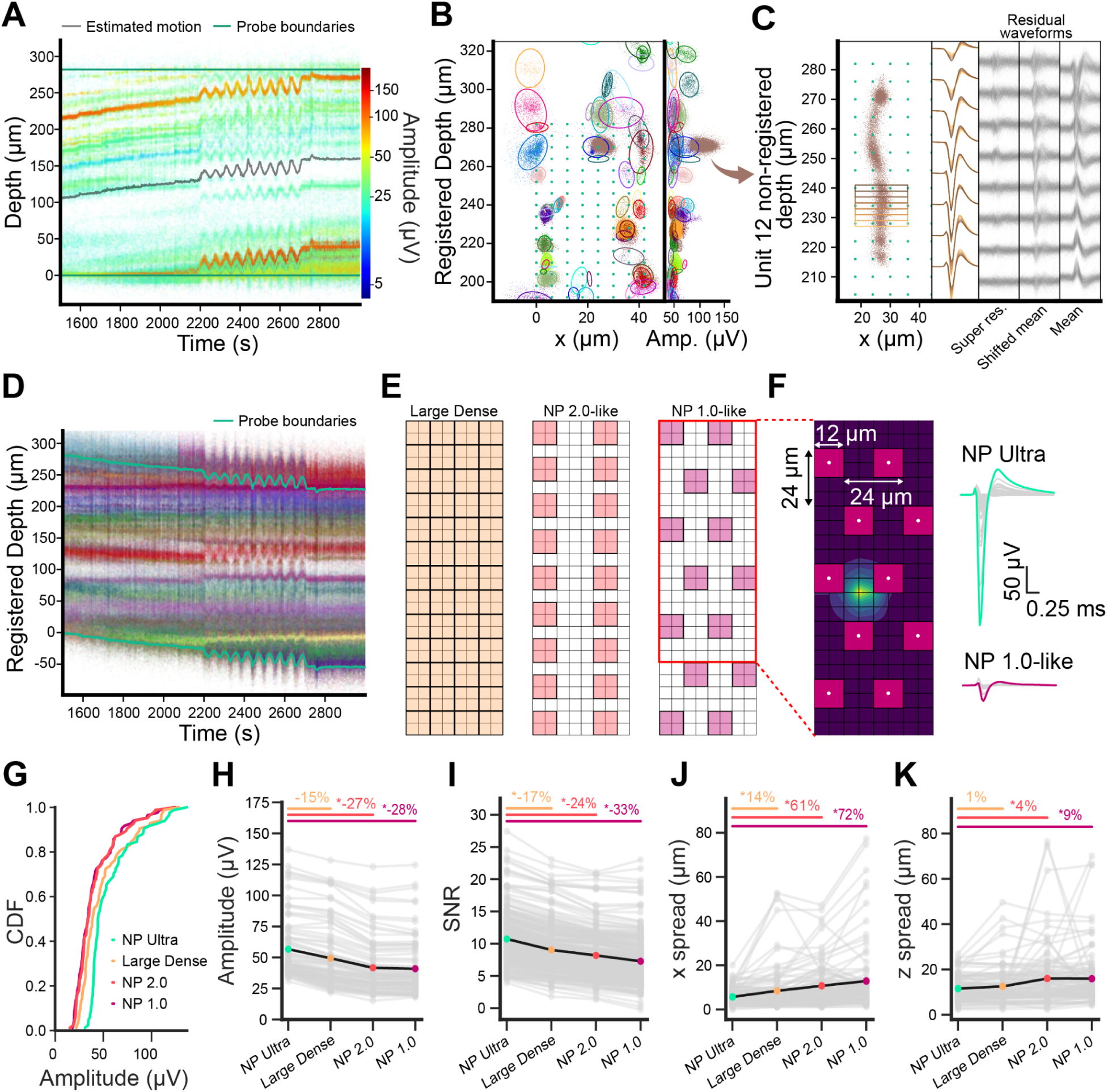
Localizing, clustering, and tracking spikes from Neuropixels Ultra probes. **A,** “Drift map” spatial spike raster of detected neurons in an example recording. Each point represents one spike; y-coordinate represents (unregistered) depth along the probe at which the spike was detected; x-coordinate detection time; color is log-scaled spike amplitude. Superimposed black line indicates the estimated motion of the probe relative to the brain. Green horizontal lines indicate top and bottom boundaries of the probe. **B,** After probe motion registration, detected spikes are localized in 2D across the face of the probe. Left panel, each point represents a spike in registered 2D space, colored by cluster identity, with ellipses representing the spatial spread of each cluster. Green markers indicate the recording site locations. Right, scatter plot of depth versus spike amplitude; colors as in left. **C,** The template waveform for cluster 12, with the super-resolution template estimates overlaid. Left, the non-registered positions of cluster 12 spikes. The colored rectangles overlaid on top of the non-registered spike positions demarcate the spikes used for estimating each super-resolution template. Middle-left, super-resolved templates for cluster 12 are overlaid, showing how these templates capture shape variability for this unit. They are shown on 8 channels corresponding to 8 rows of the middle-right column of the recording channels. Each template is colored according to the bin where it was estimated shown in the left panel. Middle, residual waveforms obtained after subtracting the super-resolution templates shifted according to the estimated motion. Middle-right, residual waveforms obtained after subtracting the mean template. Right, residual waveforms were obtained after subtracting the mean template, not shifted. **D,** Drift map as in **A**, post-motion registration. Spikes colored by their cluster identity**. E,** Three spatial resampling patterns (Large Dense, NP2.0-like, and NP 1.0-like, from left to right). Superimposed grids indicate the NP Ultra site locations (thin gray grid) and the channel size and positions for the resampling patterns (bold black grid and colored sites). **F,** Left, Grid indicating the NP Ultra recording channels (black) and the NP 1.0-like resampled sites (magenta squares). Colormap indicates amplitude as in Fig. 1C. The corresponding NP Ultra and resampled waveforms are shown on the right, with all channels overlaid on top of each other, and the main channel colored in green for the NP Ultra and magenta for NP 1.0-like resampled pattern. **G,** Cumulative distribution function of all spike amplitudes, comparing large-dense, 2.0-like, and 1.0-like site arrangements. **H,** Peak site amplitudes of neurons in the original dataset recorded with NP Ultra versus the spatially resampled versions. * indicates the statistical significance of the difference, obtained using a two-sample z-test on the unit amplitude distribution. **I,** Template SNR measured for each recording site pattern, from the original dataset and spatially resampled versions. **J,** Spatial spread, computed as the standard deviation of coordinates, in the horizontal dimension. **K,** Spatial spread in the depth dimension across the face of the probe.

The algorithm, described in detail in the Methods and demonstrated on simulated data in (Boussard et al., 2023), relies heavily on the spatial location of each spike relative to the probe, estimated using a simple point-source model (Boussard et al., 2021). We use these locations to infer the motion of the probe relative to the brain over time (Windolf et al., 2023), register the spike positions, and cluster the spikes (**Fig. 2A,B**). Algorithms that infer spatial locations of spikes and use these for clustering have proven successful on data recorded *in vitro* (Hilgen et al., 2017), but here we find that this strategy can now also be used *in vivo* given the improved ability of NP Ultra to resolve spike locations relative to NP 1.0 and 2.0. Using these cluster assignments, we estimate a template waveform for each unit at each position relative to the probe (**Fig. 2C**). This template is “super-resolved’’ in the sense that it is defined on a fine spatial grid (here chosen to be 2 µm spacing though the probe only samples at 6 µm resolution). With the motion of the unit relative to the probe, each unit’s waveform is sampled at different positions over time; we bin spikes localized on this fine spatial grid to compute the unit’s super-resolved shape. Using computational simulations of extracellular waveforms, we estimated that this spatial sampling (2 µm) should be sufficient to capture the relevant details of the extracellular action potentials along the length of the probe (**Supp. Fig. S7**). Computing the super-resolution template, and shifting it according to a precise motion estimate, provides an accurate approximation of the neuron’s spike shapes at any time in the recording, demonstrated by small residuals (**Fig. 2C**). Finally, these templates are used to deconvolve the original data, extracting spike times for each neuron and resolving spatiotemporally overlapping spikes. During template deconvolution, the estimated super-resolution templates are shifted according to the estimated motion (rather than attempting to shift and interpolate the raw data, as in Kilosort2.5 (Steinmetz et al., 2021)) to infer a waveform shape best matching each spike at that moment in time (**Fig. 2D**). Our strategy of estimating super-resolution waveforms and shifting them according to probe motion to optimize deconvolution is novel relative to previous approaches. Using this sorting algorithm, we automatically sorted about one hundred neurons from recordings with NP Ultra probes in mouse visual cortex (113 ± 62 total neurons, 44 ± 22 neurons with reliable visual responses per recording, mean ± S.D., n=6 recordings).

We next asked whether spikes recorded with NP Ultra probes have higher peak waveform amplitudes than could be measured with previous Neuropixels probes, due to the increased spatial sampling. To test this, we spatially resampled our NP Ultra recordings to simulate the signals that would have been recorded from the exact same position under different recording site patterns: an NP 1.0-like pattern with staggered recording sites and large spacing; an NP 2.0-like pattern with aligned recording sites and tighter spacing; and a ‘Large Dense’ pattern with large 12×12 µm recording sites as in NP 1.0 and 2.0, but with dense, gap-less spacing as in NP Ultra (**Fig. 2E,F**). This computational resampling strategy produces datasets exactly matched in terms of the spiking events present, differing only in the sampling density, thus permitting us to directly compare the properties of the signal as observed with NP Ultra to what would have been observed with other, less-dense probes.

Peak spike amplitudes were highest in the original NP Ultra recording (**Fig. 2G,H**; Ultra vs 1.0, 28% median relative difference, p = 4.4e-5; Ultra vs 2.0, 27% difference, p=6.9e-5; Ultra vs Large Dense, 15% difference, p=0.06; two-sample z-test on the distribution of amplitudes in each dataset). Reductions in peak site amplitude observed on the large-dense pattern reflect the effect of larger recording sites relative to NP Ultra, while reductions in amplitude under the 1.0-like and 2.0-like site patterns reflect the effect of larger sites in addition to a failure to sample the peak location due to gaps between sites. Large relative to small recording sites therefore account for about a 15% reduction in spike amplitude, and greater site spacing results in a further 14% reduction on average. The effect of site size and site density is a function of the spatial footprint of the signal; that is, a neuron whose extracellular action potential is visible over a large region will be relatively unaffected by sampling with a low site density. Therefore, the impact of sampling density on amplitude depends on the local population of neurons and varies across brain regions. We have characterized this for several regions (**Supp. Fig. S8, S9** and **S10**). Together, these results demonstrate that NP Ultra records larger amplitude signals from individual neurons.

We next asked whether the higher amplitude of spikes measured with NP Ultra (**Fig. 2G,H**) overcomes the higher noise of its small sites (**Fig. 1F**), resulting in higher SNR. We computed a “Template SNR” for each neuron by convolving the identified template waveform with the spikes identified as belonging to the neuron or with samples from random other time points, and computing the SNR between these two distributions (see Methods; **Fig. 2I**). Neurons recorded with NP Ultra had the greatest template SNR, followed by the Large Dense, 2.0-like, and 1.0-like site patterns in order (Ultra vs 1.0, 33% difference, p = 5.5e-24; Ultra vs 2.0, 24% difference, p = 1.4e-14; Ultra vs Large Dense, 17% difference, p = 6.4e-7.). As with amplitude, the magnitude of the effect varies across brain regions, depending on the distribution of unit footprint (**Supp. Fig. S9**). Using a spatial resampling strategy with a range of site size and spacing, our data suggest that an intermediate site size, and sampling without gaps between sites, may comprise an optimal site layout for most neurons in most mouse brain regions (**Supp. Fig. S10**). Thus, despite higher noise levels at individual channels, NP Ultra ultimately yields higher SNR for identifying spikes from recorded neurons.

The increased recording site density of NP Ultra also produced substantially greater precision in the estimated spatial position of each spike relative to previous probes (**Fig. 2J-K**; **Supp. Fig. S8, S9**). The success of the spike sorting strategy described above depends on the ability to spatially segregate spikes from individual neurons relative to nearby neurons. To understand how much this strategy will gain from the high sampling density of NP Ultra, we compared the spatial scatter in estimated spike positions from sorted neurons in the original data to the scatter obtained when estimating the spatial positions of the same spikes in the spatially resampled data (**Fig. 2J-K**; **Supp. Fig. S11**). Spikes were estimated with substantially higher precision in the NP Ultra dataset along the horizontal dimension of the probe (and to some extent also along the vertical dimension; **Fig. 2J**; Ultra vs 1.0, 72% difference, p = 5e-6; Ultra vs 2.0, 61% difference, p=1.6e-6; Ultra vs large-dense, 14% difference, p=0.013. **Fig. 2K**; Along vertical dimension: Ultra vs 1.0, 9% difference, p = 0.001; Ultra vs 2.0, 4% difference, p=0.002; Ultra vs large-dense, 1% difference, p=0.34.), confirming that these probes are well-suited to employ this spike localization-based sorting strategy.

We next asked whether improved SNR translated to improved yield of sortable neurons. Here, we evaluated yield by focusing on neurons that were present throughout the recording and had a biologically meaningful response; in this case, reliable and selective responses to visual stimuli. We collected new datasets from the primary visual cortex in which awake, head-fixed mice were presented with a battery of 118 natural images. Neurons were included in the final yield calculation if they had a significant visual “fingerprint”, defined here as visual responses that are reliable (i.e., consistent across repetitions of the same image) and selective (i.e. differ in a stereotyped manner across different images) (see Methods; **Fig. 3A-C, Supp. Fig. S12**). We ran our new spike sorting algorithm on these datasets and on the spatially resampled versions of the datasets and found that the yield of stably firing neurons with reliable visual responses was highest for NP Ultra, which yielded over 20 extra neurons on average than the 1.0-like pattern, and more than the large dense and 2.0-like patterns (**Fig. 3D-E**, mean ± standard error of the mean (SEM) difference in neurons: Ultra vs large-dense, 4.5±2.2, p=0.047; Ultra vs 2.0, 20.2±3.5, p=0.016; Ultra vs 1.0, 22.2±4.3, p=0.016). The large difference in yield between Ultra and 1.0 amounted to an approximately 3-fold improvement (3.05x greater yield of NP Ultra relative to 1.0-like on average). This yield difference did not arise from over-splitting of neurons in NP Ultra relative to other patterns (**Supp. Fig. S13**). NP Ultra recorded from more (visually reliable) neurons partly because it captured more neurons with low amplitude than the 2.0 and especially the 1.0 probes (**Fig. 3F**, median amplitude, bootstrapped 95% CI in μV: NP Ultra = 48.5, 44.4-54.2; large-dense = 39.7, 35.1-45.0; 2.0-like = 54.6, 49.0-63.3; 1.0-like = 67.5, 50.0-80.78). When further filtering stably firing and visually reliable neurons for those passing a sliding refractory period metric (International Brain Laboratory et al., 2023), the yield differences between NP Ultra and resampled patterns are relatively smaller, though NP Ultra still improved the yield of these well-isolated single neurons relative to NP 1.0 (1.59x greater yield of NP Ultra relative to 1.0-like on average; mean ± SEM difference in neurons = 2.8±0.7; p = 0.016 **Fig. 3E**). In sum, when recording from spatially restricted regions with a high density of neurons such as the deep layers of primary visual cortex, the higher site density of NP Ultra can greatly increase the yield of neurons with biologically meaningful responses compared to lower density probes.

**Figure 3.**
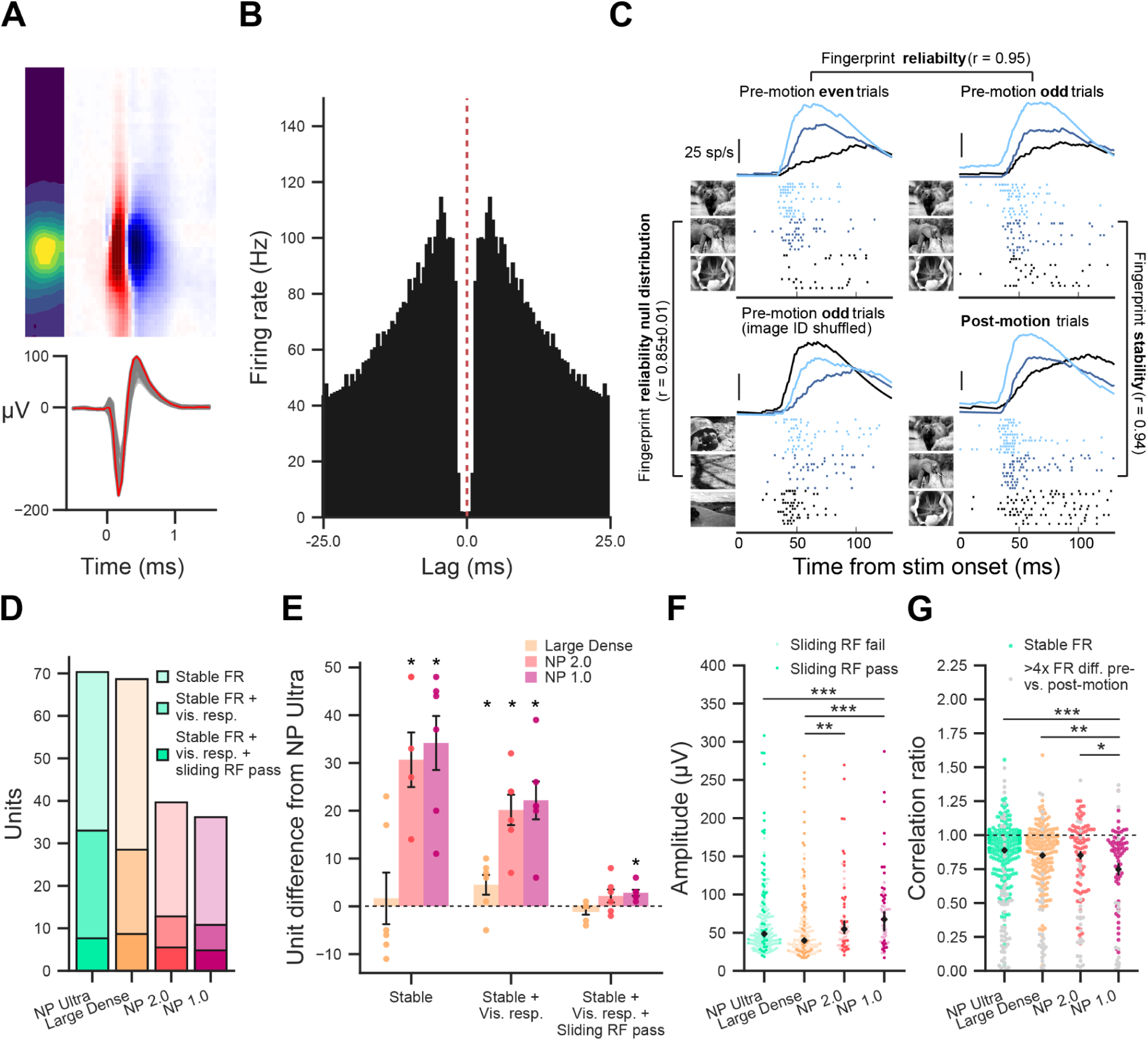
Tracking neurons with NP Ultra using visually-evoked fingerprint responses. **A**, Spatial (top left), spatio-temporal (top right) and temporal (bottom) waveforms recorded by NP Ultra of an exemplar neuron shown in B and C. **B**, Autocorrelogram of exemplar neuron shown in A and C. **C**, Peri-stimulus spike rasters and average rate (smoothed with causal half-gaussian filter with standard deviation = 45 ms) for an example neuron with reliable and stable visual response, aligned to the onsets of 3 of the 118 visual stimuli (indicated in 3 different bluish hues). Top two plots show odd and even trials from the pre-motion period, revealing a close match. Bottom left plot shows shuffled image ID control, with a poor match. Bottom right shows the post-motion responses, which again match the pre-stimulus un-shuffled responses. **D**, Mean yield of neurons (across n=6 sessions) from NP Ultra and spatially resampled datasets progressively filtered by the following three metrics: ≤4x difference in pre vs. post-motion firing rate (FR) in spikes/second (“Stable FR”), reliable visual response (“vis. resp.”), and 90% confidence of ≤20% contamination in the sliding refractory period metric (“sliding RF pass”). **E,** Differences in neuron yield between NP Ultra and each spatially resampled pattern calculated per session, using the same quality metrics as in D. Bars and error bars are mean±SEM. Stars indicate significant p-values from the Wilcoxon signed-rank test. **F**, Amplitudes of stable neurons with reliable visual responses across all sessions. Black diamonds and error bars are median and n=10000 bootstrapped 95% confidence interval of the median. Stars indicate significant p-values from Mann-Whitney U test. **G**, Stability ratios (ratio of correlations between mean pre- vs. post-motion and mean pre- vs. pre-motion visual fingerprints; see **Supp. Fig. S12B,C**) for neurons with reliable visual responses across all sessions. Colored points indicate neurons with a stable firing rate from the pre-to post-motion period; gray dots indicate neurons with unstable firing rates. Black diamonds and error bars are mean±SEM only from neurons with stable firing rates. Stars indicate significant p-values from unpaired two-sample t-test; only stable firing rate neurons included in the test. For all panels, * indicates p<0.05; ** indicates p<0.01, *** indicates p<0.001.

We additionally reasoned that the higher site density of NP Ultra would afford a superior ability to stably track drifting neuron locations across time. To test this, we imposed probe motion as a ground truth pattern of unstable probe location relative to the brain during the visual fingerprint sessions (Steinmetz et al., 2021). We presented each image an average of 30 times before moving the probe, and 15 times afterwards. To assess tracking stability, we computed a stability ratio for each neuron exhibiting reliable visual responses by dividing the mean pre- vs. post-motion fingerprint correlation by the mean pre- vs. pre-motion correlation. These mean correlations are calculated by randomly splitting the 30 pre-motion repetitions of all images into two non-overlapping 15-repetition subsets 1000 times, and correlating the fingerprints obtained from one pre-motion trial subset and from the other pre-motion subset (pre- vs. pre-motion) or from the post-motion repetitions (pre- vs. post-motion) (**Supp. Fig. S12B-C**). A stability ratio of 1 indicates a stably tracked neuron: the visual responses before and after motion are just as similar as those from two subsets of the trials before motion. Pooling the stability ratios of neurons within patterns across sessions, we found that there was no significant difference in the mean ratios between NP Ultra and large-dense or NP 2.0-like patterns (Ultra vs. large-dense, 4.1% difference, p=0.12; Ultra vs. 2.0, 2.3% difference, p=0.50); but that the mean NP 1.0-like stability ratio is significantly lower than all other patterns (Ultra vs. 1.0, 19.7% difference, p = 4.8e-5; large-dense vs. 1.0, 15.0% difference, p = 4.4e-3; 2.0 vs 1.0, 17.0% difference, p = 0.015) (**Fig. 3G**). These findings are corroborated by computational simulations of extracellular action potentials (EAP) sampled at different spatial densities, which reveal that significant information for reconstructing the EAP begins to be lost around the NP 1.0 spatial frequency, but is relatively preserved at higher spatial frequencies (e.g. smaller site spacing; **Supp. Fig. S7**). Thus, the extra spatial resolution available from NP Ultra does not provide additional improvements in motion correction compared to Neuropixels 2.0 probes, indicating that the 2.0 probe site density is likely sufficient for effective motion correction.

### Subcellular recordings from axons and dendrites in mouse isocortex

Given the improved ability to detect and sort the spikes from individual neurons, we sought to determine whether NP Ultra probes can record extracellular action potentials from neuronal structures typically under-sampled by other probes. For instance, units or subcellular structures with waveforms that decay over distances smaller than 20 μm would likely not be detected by NP 1.0 probes if an action potential was initiated between electrode contacts. Indeed, in recordings from the visual cortex of awake mice, we noted a substantial number of extracellular action potentials that had large amplitude but were present only on a few recording channels of NP Ultra (**Fig 4A**). To quantify this observation, for each waveform we calculated its *spatial footprint* as the radius from the peak channel at which the average spike amplitude fell below 30 μV, a value sufficiently above the noise level. We observed a substantial fraction of units with a footprint less than 20 µm which we define as having a *small* footprint (**Fig 4B**).

**Figure 4.**
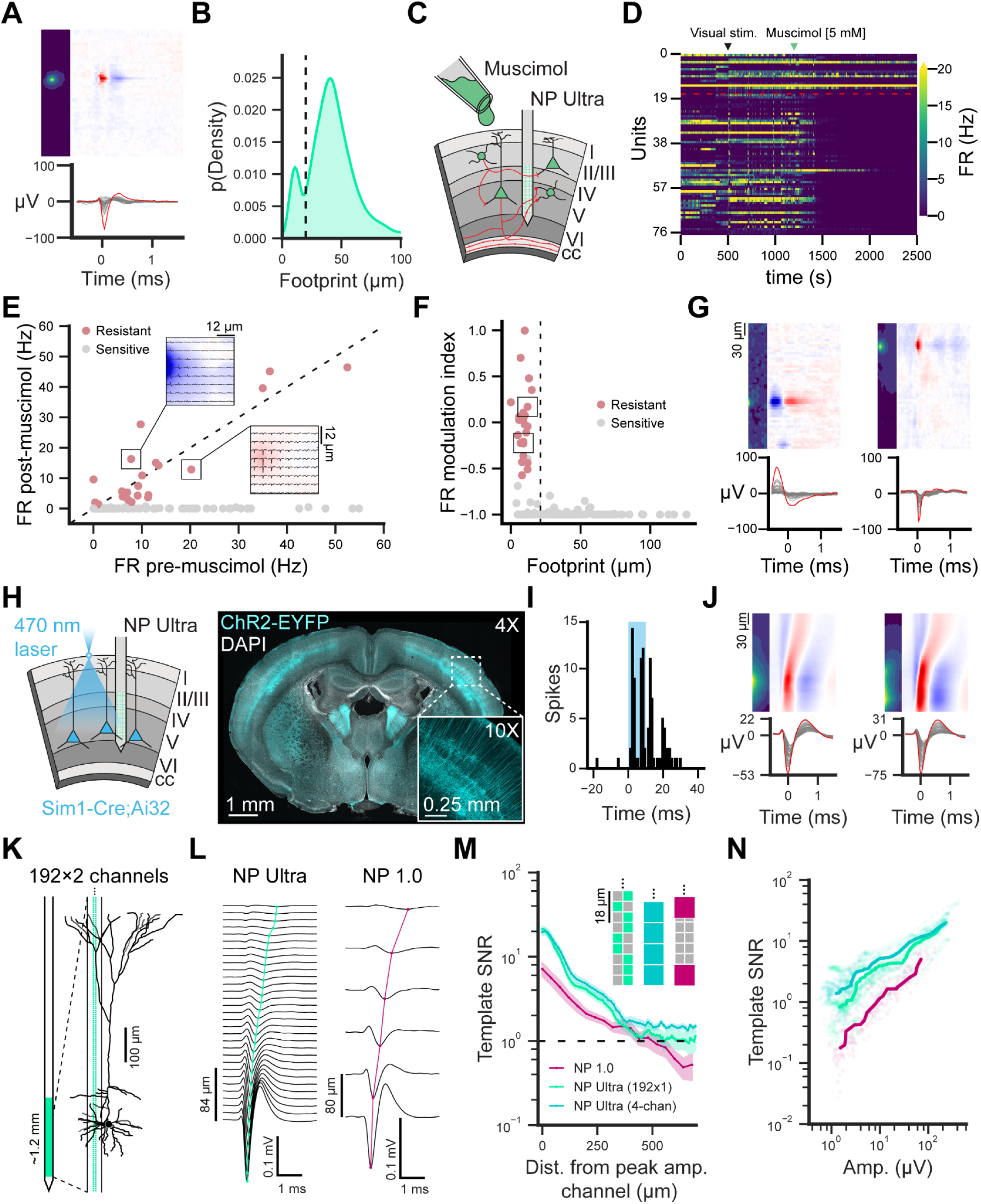
Recordings from subcellular compartments. **A**, Example small-footprint unit recorded from primary visual cortex. **B**, Kernel density histogram of measured spatial footprint across a total of 9,496 units recorded in visual cortex. Dashed line denotes the boundary (20 μm) between what we define as large and small footprint units. **C**, Schematic of NP Ultra recordings in cortex with surface application of muscimol [5 μM]. Red lines indicate axonal segments of intra- and extra-cortical neurons. **D**, Raster plot showing the firing rate of 77 units during an exemplar NP Ultra recording. Black arrow = time of visual stimulation, green arrow = time of muscimol application. Units are sorted by ascending footprint size, the boundary between large and small units is denoted by the red dashed line. **E**, Scatter plot of firing rate across all units measured within a 100 ms window pre- and post-muscimol application, colored by where a given unit was resistant or sensitive to muscimol. Dashed line indicates identity between pre- and post-FR. Insets display the waveforms and heat maps of two example muscimol-resistant units. **F**, Relationship between the spatial footprint and firing rate modulation index for all recorded units. Units from **E** marked by black boxes. **G**, Full waveform plots of the two units in **E**, respectively. **H** (left), Schematic of optotagging experiment in Sim1-Cre;Ai32+ L5b pyramidal neurons. (Right) histology from a Sim1-Cre;Ai32 animal showing labeled L5 pyramidal neurons in S1. **I**, PSTH of an example Sim1+ unit in response to 10 ms 470 nm photostimulation. **J**, Two example Sim1+ units with visible potentials propagating along the NP Ultra probe. **K**, Schematic of a L5 cell recorded with a switchable NP Ultra probe in the linearized (192 × 2 channels) configuration. **L**, Example bAPs recorded with NP Ultra (left) and NP 1.0 (right). Traces show spike-triggered average waveforms (n = 2,000 spikes) recorded from the column of channels that includes the peak amplitude channel. Colored traces track the minimum voltage recorded on each channel with a recorded amplitude above 5% of the maximum. **M**, Dendritic channel template SNR measured as a function of vertical distance from the peak amplitude channel for NP Ultra (192×2 configuration, best column selected, green), and NP Ultra (192×2 configuration, 4-channel average, cerulean) or NP 1.0 probes (magenta). Channel configuration diagrams are shown on the top right. **N**, Template SNR plotted against channel waveform amplitude for each channel in every probe configuration from **M**.

Though these waveforms had diverse temporal characteristics, we hypothesized that they may originate within axons on the basis of their small spatial extent. To test this possibility, we applied the GABA agonist muscimol to the cortex to suppress somatic spiking while leaving the activity of non-local axons unaffected. Consistent with our hypothesis, all recorded waveforms that survived muscimol application (firing rate > 1 spike/s in muscimol) had a small footprint (23/175, n=6 sessions, n=3 mice; **Fig. 4C-G**). Importantly, not all small footprint units survived muscimol application, which is consistent either with axons of local origin or with a subset of small footprint waveforms belonging to an as yet unidentified small cell type (as discussed later). Aside from their small spatial footprint, the muscimol-resistant axonal waveforms were inconsistent in appearance: some had narrow, negative peaks with little or no early positive component while others had prominent early positive peaks (**Supp. Fig. S14**). Similarly, spike waveforms histologically localized to the corpus callosum could have either or both of these waveform characteristics (**Supp. Fig. S15A-C**). Biophysical simulations confirmed that small-footprint, narrow, negative spikes could be observed at nodes of Ranvier (**Supp. Fig. S15D;** (Halnes et al., 2024)), that positive peaks may arise in unmyelinated segments of otherwise myelinated axons (**Supp. Fig. S15E**), and that unmyelinated axons are unlikely to be detected given their small amplitude except when they have substantial branching and termination (**Supp. Fig. S15F;** (McColgan et al., 2017)). Therefore, the characteristics of axonal spikes vary dramatically depending on the exact properties of the axon recorded, but are unified by their small spatial footprint rather than any particular waveform-peak direction, number, or width.

We considered that NP Ultra may also permit recording from dendritic compartments of neurons with higher resolution and SNR than previous technologies. Extracellular recordings of back-propagating action potentials in apical dendrites of cortical pyramidal neurons have been reported with NP 1.0 (Jia et al., 2019) and with other probes (Buzsáki and Kandel, 1998). We made recordings with NP Ultra and NP 1.0 using an insertion strategy to obtain recordings in mouse V1 approximately aligned with apical dendrites. First, we opto-tagged layer 5 pyramidal neurons (**Fig. 4H-I**, Sim1-Cre;Ai32), and observed propagation in waveforms consistent with known back-propagation waveform characteristics (i.e. initial positive peak at the time of the somatic spike; (Gold et al., 2006)) and speed (Stuart et al., 1997; **Fig. 4J**). Then, to record dendritic signals across a larger extent of the apical tree (**Fig. 4L)**, we switched to the linear 192 × 2 configuration (**Fig. 4K)**. With this configuration, we recorded back-propagation dynamics spanning more than 400 μm from the putative somatic channel, at ∼7x higher spatial resolution than with NP 1.0 (6 µm linear spacing versus 40 µm; **Fig. 4L**). This enhanced resolution led to significantly greater SNR in resolving back-propagating signals across the apical dendrite (**Fig. 4M**). These improvements over NP 1.0 recordings did not depend on probe alignment or recording quality, as they persisted when comparing dendritic signals of similar amplitudes (**Fig. 4N**). In addition to back-propagation, our dataset revealed a number of distinctive waveforms with spatiotemporal characteristics not previously reported, which appear similar to biophysical simulations of close dendritic apposition (**Supp. Fig. S16**). Taken together, NP Ultra enables subcellular recordings from dendrites with improved resolution and SNR relative to prior devices.

### Units with axon-like small footprints are prevalent across the mouse brain

To assess the prevalence of small footprint, axon-like extracellular action potentials across the brain, we used NP Ultra probes to make recordings from 4,666 single units in 18 brain regions (>50 single units/region; n=4 mice; n=12 sessions; **Fig. 5A**). In each experiment, we inserted probes and recorded for a short time (∼5 minutes) at a given recording site before advancing the probe 250 µm, then repeating this process several times over many recording locations. Targeted areas included several regions of the isocortex, hippocampus, basal ganglia, thalamus, midbrain, and corpus callosum. We sorted spikes from each recording position separately (see Methods) and compiled the waveforms of all single units, which exhibited a range of diverse spatial features (**Supp. Fig. S17**).

**Figure 5.**
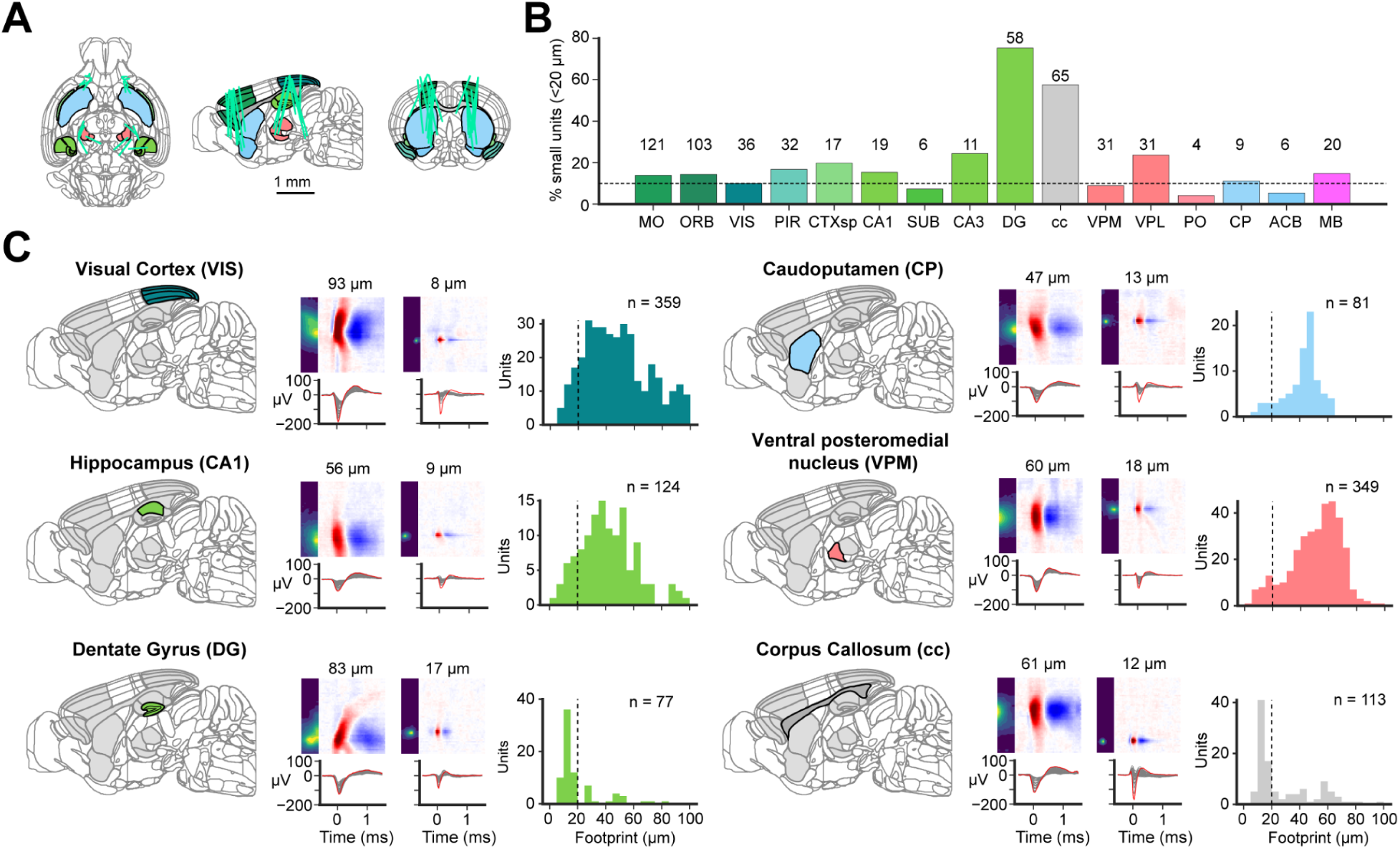
NP Ultra recordings reveal small electrical footprint units in many brain regions of the mouse. **A**, NP Ultra insertion trajectories from different mice and sessions, registered to the CCF atlas coordinates. **B**, Percentage of units with small footprints less than 20 μm in regions with the most recorded single units. Number of small units per region indicated. Dashed line corresponds to 10% **C**, Two example waveforms with one large and one small spatial footprint from each of six different brain regions, and spatial footprint histogram of all units from these regions. MO: Somatomotor areas; ORB: orbital area; VIS: visual areas; PIR: Piriform area; CTXsp: cortical subplate; CA1: hippocampal CA1; SUB: Subiculum; CA3: hippocampal CA3; DG: Dentate gyrus; cc: corpus callosum; VPM: ventral posteromedial nucleus of the thalamus; VPL: Ventral posterolateral nucleus of the thalamus; PO: Posterior complex of the thalamus; CP: Caudoputamen; ACB: Nucleus accumbens; MB: midbrain.

To determine whether we could sample otherwise unobservable or under-sampled units, we examined the extracellular field features of the units we recorded. As in NP 1.0 recordings (Jia et al., 2019), we observed a wide distribution of single channel spike waveform features, such as amplitude, peak-to-trough ratio (PTR), and spike duration, across brain regions (**Supp. Fig. S18, S19**). With the high density NP Ultra probes, we are able to extract an additional feature, “spatial footprint”, as described above (**Fig. 4**). In each brain region we recorded, a significant fraction of extracellular action potentials (>10% in most regions) had a spatial footprint of less than 20 µm (**Fig. 5B,C**). This fraction of small units was significantly greater in recordings localized to the corpus callosum (57.5%, 31.4 ± 26.1 μm, mean ± SD, 113 units) and the dentate gyrus (75.3%, 20.0 ± 16.2 μm, 77 units), compared to primary visual cortex (VISp), for example (10.0%, 50.0 ± 25.6 μm, 359 units; **Supp. Fig. S20**). These small-footprint units may reflect recordings from axons or from neurons with exceptionally small somata, such as granule cells, or both.

### Small extracellular action potentials across species

To gain further insight into the nature of such small extracellular action potentials, we compared the spatial footprints recorded across four vertebrate species. Species with larger brains tend to have larger neurons (Herculano-Houzel et al., 2014; Beaulieu-Laroche et al., 2021). Different species have specialized brain regions with varying neuronal densities, tailored to behavioral demands through evolution. For instance, whereas the cerebral cortex constitutes the majority of brain volume in primates, weakly electric mormyrid fish possess a greatly enlarged cerebellum that covers their entire dorsal brain surface (Sukhum et al., 2018). We collected high resolution waveforms using NP Ultra from brain areas three additional species: monkey (*Macaca nemestrina*) visual cortex, electric mormyrid fish (*Gnathonemus petersii*) cerebellum, and bearded dragon (*Pogona vitticeps*) medial cortex (an area homologous to mammalian dentate gyrus (Tosches et al., 2018)).

We observed a substantial number of units with a footprint less than 20 μm in each species (**Fig. 6A,B**). We collected 124 units from the monkey visual cortex, and found 13 units with footprints smaller than 20 μm (10.5%). This closely matches the observation in the mouse visual cortex (36/359, 10.0%). The similar distribution of footprints between mouse and monkey visual cortex is consistent with the known similar distribution of somatic sizes between these two areas (Gilman et al., 2017). A similar proportion of units in the lizard medial cortex had small footprints (2/19, 10.5%). Close to half of the units detected in the cerebellum of the electric fish had footprints below 20 μm (18/37, 48.7%). Together these observations demonstrate that units with small spatial footprints (<20 µm), which are difficult to detect with lower density probes, are consistently detected with NP Ultra across species.

**Figure 6.**
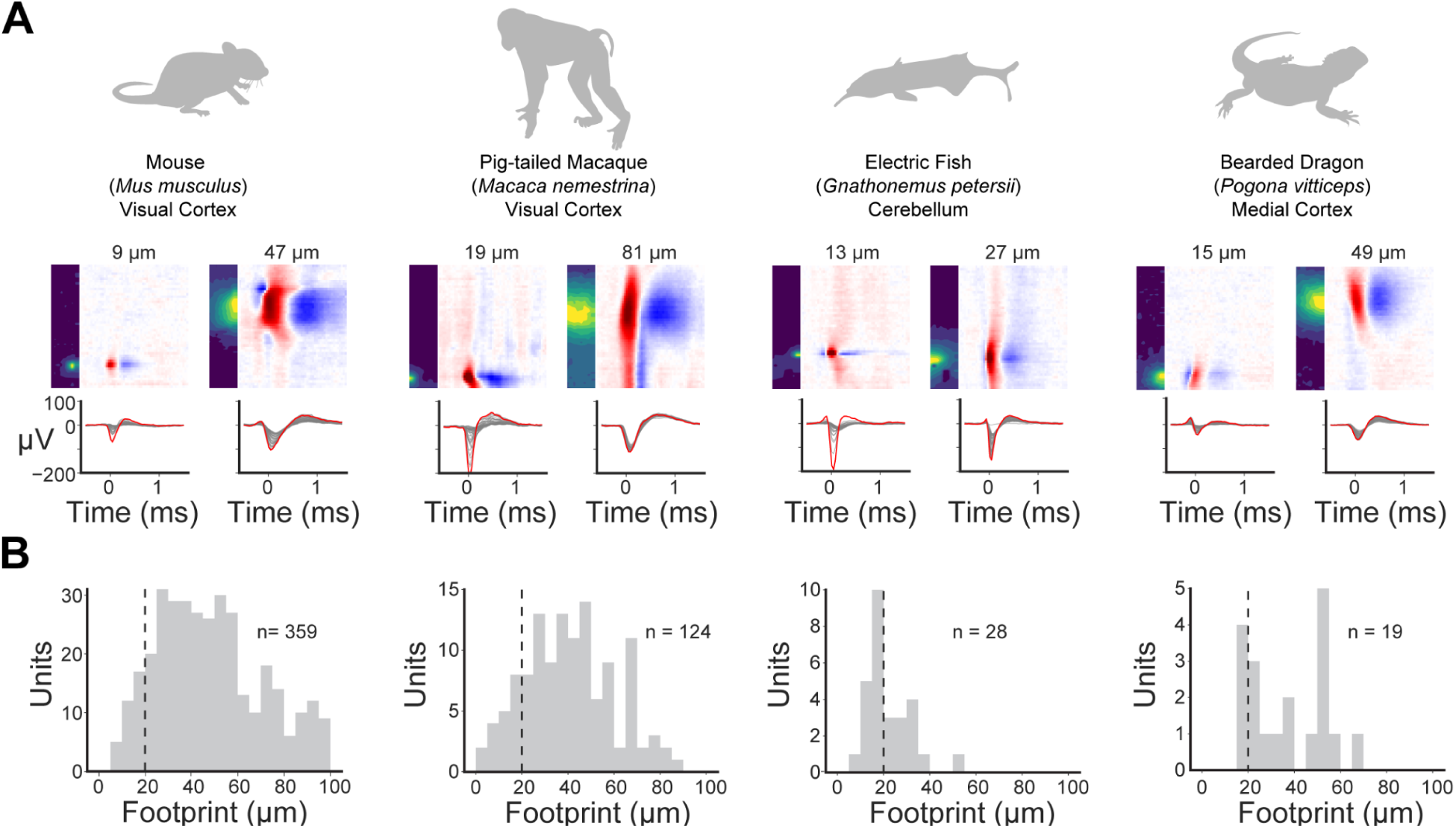
Small footprint units are detected across different species. **A**, Two example waveforms from four species (mouse visual cortex, monkey visual cortex, electric fish cerebellum, and bearded dragon medial cortex), with one large and one small spatial footprint. Two waveforms with similar peak amplitude in the same species were selected for comparison. **B**, Spatial footprint histogram of all units from each species.

### Characterizing and identifying small-footprint action potentials

The brain-wide recordings in mice described above revealed a remarkable diversity of spatio-temporal waveforms (**Supp. Fig. S17**). Even just considering units recorded in the cortex, we observed a wide variety of unique spatial and temporal profiles ranging from small compact units to larger, more spatially extended waveforms. We hypothesized that this diversity might reflect spike waveforms recorded from distinct cell types. For instance, we wondered whether some small footprint units in the cortex might correspond to a distinct interneuron type, in addition to those identified as axonal in the muscimol experiments described above. We collected a new dataset focused on visual cortex (n = 40 recordings, 20-30 minutes each, n = 8,896 units) and found that a large proportion of small footprint units had short duration waveforms, suggestive of fast-spiking interneurons (**Fig. 7A**). Furthermore, when these data were segregated by spike duration (**Fig. 7B**), we found that fast-spiking (FS, duration < 0.4 ms; n = 2,058) units displayed a distinctly bimodal footprint distribution compared to regular-spiking (RS, duration ≥ 0.4; n = 6,838) units, which were more uniformly distributed about the mean (44.8 ± 20.0 μm). The set of FS units, therefore, appeared to encompass both small and large footprint units and were potentially composed of more than one cell type.

**Figure 7.**
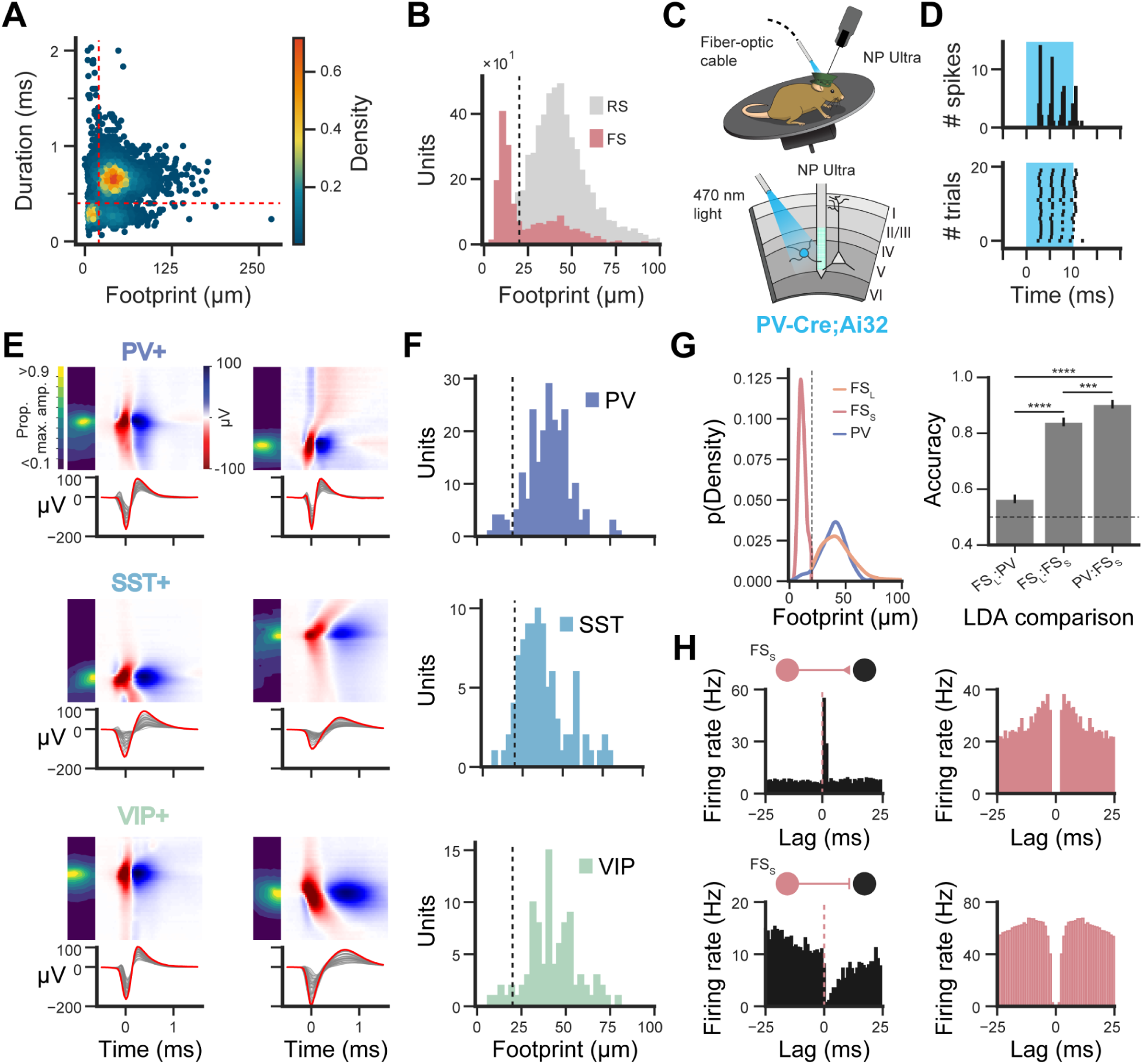
Optotagging three inhibitory neuron classes measured with NP Ultra. **A**, Scatter plot of waveform duration and footprint size, colored by the density of units within a given region of the graph. Note the existence of two maxima, one with relatively long waveform durations and large footprint, and another with short-duration waveforms and small footprint. **B**, Distributions of footprint size for all units, segregated by waveform duration (RS = regular-spiking, FS = fast-spiking). FS units displayed a bimodal distribution split at 20 μm. Black dashed line indicates the demarcation between “large” and “small” units. **C**, Schematic of experimental setup in which neurons in visual cortex transgenically expressing channelrhodopsin were recorded with NP Ultra probes and optotagged using 470 nm light. **D**, Trial-wise optogenetically-driven responses of an example PV-Cre;Ai32-expressing unit during a 10 ms 470 nm light pulse. **E**, Two example spatial, spatio-temporal, and temporal waveforms from optotagged units in each of the examined transgenic mouse lines (PV-Cre;Ai32, SST-Cre;Ai32, VIP-Cre;Ai32). **F**, Distributions of spatial footprint for each optotagged group of units. **G**, Left, Density histograms displaying the distribution of footprint between PV and FS units. FS units are subcategorized into FS_L_ (fast-spiking, large) and FS_S_ (fast-spiking, small) based on a footprint size threshold (20 μm). Right, Accuracy of Linear Discriminant Analysis (LDA) in separating different types of FS and PV units using pre-peak-to-trough ratio (prePTR) and peak-to-trough ratio (PTR) as features. **H**, Left, Cross-correlograms displaying the firing rate of example units as a function of spiking in two FS_S_ units (top/bottom rows). Dashed line indicates the time of an FS_S_ unit spike. Right, Autocorrelograms of the example FS_S_ units.

To measure and compare the spike waveforms of genetically identified cells, we performed optotagging of inhibitory neuron types in the visual cortex. We used transgenic mice to express channelrhodopsin (ChR2) in three major inhibitory neuron subclasses in the cortex: parvalbumin- (PV), somatostatin- (SST), and vasointestinal polypeptide-expressing (VIP) cells. During the recording session, we photostimulated the cortex causing these neurons to spike and thereby ‘optotagging’ them (**Fig. 7C**; (Lima et al., 2009)). Optotagged units had short latency and stereotyped spiking responses to photostimulation (**Fig. 7D**). 243 PV, 116 SST, and 126 VIP interneurons were identified as optotagged based on both unsupervised density-based clustering methods and qualitative assessment (**Fig. 7B-D, Supp. Fig. S21**, Methods). Example waveforms of each inhibitory neuron subclass are shown in **Fig. 7E** with additional examples in **Supp. Fig. S22** and **Supp. Movie S1**.

We computed the spatial footprint of each optotagged unit and found that a minority of units within each optotagged group had a small footprint. SST+ units had the largest share (9%) of units with a small footprint, followed by VIP+ (8%) and PV+ (6%; **Fig. 7F**, **Supp. Fig. S23**). Thus, small footprint units were not exclusively found within one interneuron group. In contrast, the proportion of *untagged* FS units with a small footprint was much larger (58%**; Supp. Fig. S23**). Based on this observation, we subdivided the untagged FS group into two categories based on their footprint (greater or less than 20 μm), which we call large fast-spiking (FS_L_, n = 952) and small fast-spiking (FS_S_, n = 1,133) units. FS_L_ and FS_S_ had distinct waveforms, with FS_S_ having smaller peak-to-trough ratios (**Supp. Fig. S23**).

Given that FS units have long been assumed to be mostly PV+ fast-spiking interneurons in extracellular electrophysiology, we set out to test the relationship between FS_L_, FS_S_, and PV units recorded with NP Ultra. We first compared the distributions of their respective footprint sizes and found highly overlapping distributions between FS_L_ and PV, but not FS_S_ (**Fig.7G**, *left*). We next examined waveform features that were independent of footprint size and spike duration and compared them across PV, FS_L_, and FS_S_; specifically, we computed the pre-trough peak-to-trough ratio (prePTR) and peak-to-trough ratio (PTR, see also **Fig. 8A**, **Supp. Fig. S23**, Methods). Pairwise comparisons of PV, FS_L_, and FS_S_ units were made through Linear Discriminant Analysis (LDA) using only these two features (**Fig. 7G**, *right*). FS_L_ and PV units were not well-differentiated, with an average discrimination accuracy of 0.56 ± 0.042 (mean ± SEM, just above chance, 0.5) suggesting that these could correspond to the same cell class. In contrast, discrimination accuracy between these two groups and FS_S_ units was higher (PV:FS_S_, 0.87 ± 0.022; FS_L_:FS_S_, 0.83 ± 0.026). Thus, FS units are not a uniform population and FS_S_ units mostly do not correspond to PV+ interneurons.

**Figure 8.**
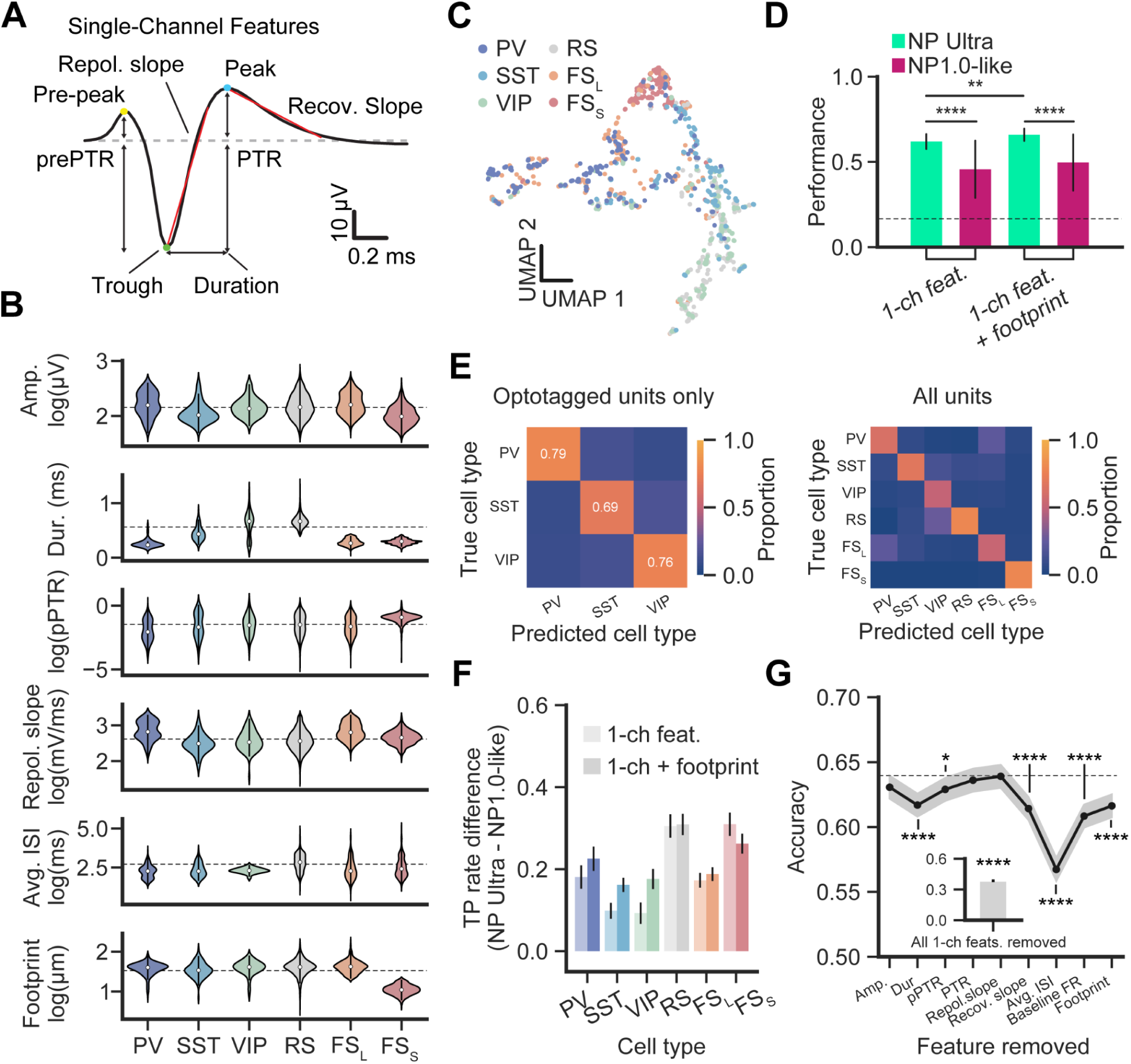
Enhanced classification of inhibitory neuron types with NP Ultra. **A**, Schematic of features extracted from single-channel waveform (1-ch features). Amplitude is the absolute difference between trough and peak. Duration is the time between trough and peak. Peak-to-trough ratio (PTR) is the ratio between absolute amplitudes of peak and trough, prePTR is the ratio of the absolute amplitudes of the pre-peak and trough. Red lines show slopes for repolarization and recovery. **B**, Comparison of waveform features extracted from the peak amplitude channel for each unit within a transgenic mouse line (all metrics except duration are log-scaled). All untagged units were split into three groups based on spike duration (RS = regular-spiking, FS = fast-spiking) and footprint size. Dots indicate the median and black vertical lines indicate the interquartile range. Dashed horizontal lines indicate the population median for each metric. **C**, UMAP projection of an equally sampled subset of NP Ultra-recorded units from each cell class. **D**, Classifier performance (out-of-bag score, mean ± S.D.) on NP Ultra (green) and NP 1.0 (blue) data using single and multi-channel features. **E**, Confusion matrices for opto-tagged only (left) and all cell class (right) classification using that 1ch features + footprint dataset. **F**, Difference in True Positive (TP) rate between NP Ultra and NP 1.0-like data for each cell class. **G**, Classifier accuracy (mean ± 95% confidence interval) across all cell classes when one feature is iteratively excluded from the dataset. Asterisks indicate significant changes in accuracy as a result of feature removal. Dashed line is the mean classifier accuracy with all 1-ch features + footprint. Inset shows the accuracy of the classifier when no 1-ch features are included in the dataset.

If FS_S_ units don’t correspond to a specific cortical inhibitory interneuron type, we considered whether they might have an excitatory downstream effect on postsynaptic partner units. To investigate this, we performed a cross-correlation analysis to identify units whose firing rate was modulated around the time of an FS_S_ unit spike. Putatively connected pairs within a recording were extremely rare (<0.02% of all observed interactions, >125,000 pair interactions), but some examples were identified (**Fig. 7H**, **Supp. Fig. S23**). Excitatory interactions were identified as those in which the post-reference spike firing rate period increased to greater than five times the pre-spike standard deviation, whereas inhibitory interactions were those in which the firing rate decreased by at least the same amount. We found instances of both putative excitatory and inhibitory interactions at short latencies (<3ms) from the time of an FS_S_ unit. FS_S_ units can therefore be either excitatory or inhibitory, further supporting the idea that these small-footprint waveforms correspond to axons of either excitatory or inhibitory neurons rather than larger, somatically initiated action potentials from any distinct class of neuron.

### Classification of cortical inhibitory neuron classes

The dense and small recording sites of NP Ultra can acquire greater resolution information about the electrical features of recorded neurons. Thus, we next sought to determine whether NP Ultra probes could differentiate cell classes (that is, broad categories of molecularly-defined neurons grouped under a larger class) better than an NP 1.0 probe configuration. We extracted single-channel waveform features using the peak amplitude channel for each unit, which revealed waveform characteristics unique to each type of neuron discussed above (**Fig. 8A**, **B**, **Fig. 7E-G**, **Supp. Fig. S23**, **Table 4**). PV+ units displayed large amplitude, short-duration spike waveforms that were highly similar to the FS_L_ units (**Fig. 7G**), especially in terms of peak-to-trough ratio (PTR) and repolarization slope. SST+ and FS_S_ units tended to have the lowest amplitude waveforms on average but were notably different in prePTR and PTR, with the former having both smaller pre-trough peaks and larger post-trough peaks relative to the magnitude of the trough than other cell classes (p_prePTR_ = 7.7 × 10^-6^, p_PTR_ = 5.43 × 10^-30^, Mann-Whitney U test, multiple comparisons corrected via Benjamini-Hochberg method, **Supp. Fig. S25**). VIP units were generally heterogenous, forming a bimodal distribution along some metrics, especially spike duration. However, VIP units as a population closely resembled untagged RS units in many respects with the notable exception of baseline firing rate, which was higher in VIP cells, and the average inter-spike interval (avg. ISI), which was lower (p_bFR_ = 4.6 × 10^-8^, p_ISI_ = 5.3 × 10^-22^, **Supp. Fig. S25**). Reducing the dimensionality of the features revealed substantial, but incomplete, segregation of the cell classes according to these features (**Fig. 8C**, **Supp. Fig. S26**). We therefore undertook a quantitative assessment of cell classes discriminability.

**Table 1.** Estimated signal attenuation of Neuropixels probes.

|  | Neuropixels 1.0<br>& 2.0 | Neuropixels Ultra | Notes |
| --- | --- | --- | --- |
| Electrode area | 144 $\mu\text{m}^2$ | 25 $\mu\text{m}^2$ | |
| $Z_{elec}$ | ~100 kΩ | 500 kΩ* | * Estimated from test structures |
| $Z_{amp}$ | 27.2 MΩ | 27.2 MΩ | |
| $Z_{elec}/Z_{amp} * 100\%$ | 0.4% | 1.8% | |

**Table 2.** Estimated noise of Neuropixels probes in saline.

|  | Neuropixels 1.0 | Neuropixels Ultra | Notes |
| --- | --- | --- | --- |
| Electrode area | 144 $\mu\text{m}^2$ | 25 $\mu\text{m}^2$ | |
| $Z_{elec}$ | ~100 kΩ | 500 kΩ* | * Estimated from test structures |
| $V_{n-elec}$ | 2.7 $\mu\text{V}_{rms}$ | 5 $\mu\text{V}_{rms}$** | ** Modeled based on estimated impedance |
| $V_{n-amp}$ | 5.4 $\mu V_{rms}$ | 5.4 $\mu V_{rms}$ | |
| $V_{n-total}$ | 6 $\mu V_{rms}$ | 7.4 $\mu V_{rms}$ | |

**Table 3.** Estimated noise of Neuropixels probes in the brain.

|  | Neuropixels 1.0 | Neuropixels Ultra | Notes |
| --- | --- | --- | --- |
| Electrode area | 144 $\mu m^2$ | 25 $\mu m^2$ | |
| $Z_{elec}$ | ~100 kΩ | 500 k$\Omega^*$ | * Estimated from test structures |
| $V_{n-elec}$ | 4.6 $\mu V_{rms}$ | 7.6 $\mu V_{rms}^{**}$ | ** Modeled based on estimated impedance |
| $V_{n-amp}$ | 5.4 $\mu V_{rms}$ | 5.4 $\mu V_{rms}$ | |
| $V_{n-total}$ | 7.1 $\mu V_{rms}$ | 9.4 $\mu V_{rms}$ | |

**Table 4.** Features used for cell type classification.

| Feature | Abbreviation | Definition | Units |
| --- | --- | --- | --- |
| Amplitude | Amp. | The absolute difference in voltage between the spike minimum (trough) and the post-trough maximum (peak). | $\mu\text{V}$ |
| Duration | Dur. | The time delay between the trough and post-trough peak. | ms |
| Pre-peak-to-trough ratio | prePTR or pPTR | The ratio between the maximum voltage prior to the trough and the trough. | ratio |
| Peak-to-trough ratio | PTR | The ratio between the maximum voltage after the trough and the trough. | ratio |
| Repolarization slope | Repol. slope | The slope of the linear fit between the trough and 0.3 ms after the trough. | mV/ms |
| Recovery slope | Recov. slope | The slope of the linear fit between the post-trough peak and 0.3 ms after the peak. | mV/ms |
| Average inter-spike-interval | Avg. ISI | The average time between spikes calculated across the recording, excluding epochs of photostimulation. | s |
| Baseline firing rate | Baseline FR | The average firing rate 1 s prior to photostimulation epochs. | spks/s |
| Spatial footprint | Footprint | The radial distance about the peak amplitude channel within which the average amplitude across channels $\leq 30 \mu\text{V}$. | $\mu\text{m}$ |

The cell class identity could be discriminated with high accuracy from recorded features, and could be discriminated better with data from NP Ultra than from NP 1.0. To analyze this, we trained a non-linear supervised classifier (random forest) to predict the identity of each neuron among RS, FS_L_, FS_S_, PV, SST, and VIP types, with balanced groups (chance performance = 0.17; see Methods). We assessed model performance when training on three different feature sets: single-channel waveform features (1-ch features; **Fig. 8A**), the single-channel waveform itself, or all 1-ch features along with the footprint radius (1-ch wf, 1-ch + footprint; **Fig. 8B,D** **Supp. Fig. S26**). Because classifier accuracy was not improved using the single-channel waveform over single-channel features using all units, we focus here only on analyses using the single-channel features and footprint. Overall, classification performance on both of these feature sets was significantly above chance (mean ± S.D. accuracy: 1-ch feats. = 0.63 ± 0.03, 1-ch feats. + footprint = 0.67 ± 0.04). To compare NP Ultra waveforms with lower resolution NP 1.0 waveforms, we spatially resampled data from NP Ultra-recorded units to provide a direct unit-for-unit comparison (**Fig. 8D**, **Fig. 2E**, see Methods). NP Ultra-recorded units had better performance than NP 1.0, regardless of the features used in classification (mean accuracy for NP 1.0-like: 1-ch feats. = 0.44 ± 0.16, 1-ch feats. + footprint = 0.46 ± 0.15, p << 0.001 for each comparison, independent t-test, multiple comparisons corrected via Benjamini Hochberg method). The improvement in performance of NP Ultra over NP 1.0-like data extended to classification using all six cell class categories as well: RS units were classified with the highest accuracy (mean ± SD = 0.83 ± 0.10) followed by FS_S_ (0.71 ± 0.08), SST (0.67 ± 0.10), PV (0.59 ± 0.09), VIP (0.49 ± 0.09), and FS_L_ (0.48 ± 0.11, **Fig. 8E,F**).

Classification performance was even higher when considering the cell class for which we have ground truth identification, and still superior for NP Ultra relative to 1.0. We demonstrate this here by performing random forest classification using only those units that had been optotagged (PV, SST, and VIP; **Fig. 8E**). The accuracy among only these three types was high (0.75 for 1-ch features; 0.77 for 1-ch waveform; 0.74 for 1-ch features + footprint). We also found that while accuracy was similar for NP Ultra and NP 1.0-like data using 1-ch features (NP Ultra = 0.75 ± 0.05, NP 1.0-like = 0.76 ± 0.11, p = 0.57, **Supp. Fig. S26**), NP Ultra outperformed NP 1.0 when including spatial footprint (1-ch waveform: NP Ultra = 0.77 ± 0.05, NP 1.0-like = 0.68 ± 0.05, 1-ch features + footprint: NP Ultra = 0.74 ± 0.05, NP 1.0-like = 0.56 ± 0.07, p_1-ch feats + footprint_ = 3.6 × 10^-42^), likely because NP Ultra was better able to estimate spatial footprint due its increased site density.

The importance of spatial footprint as a feature useful for discriminating cell class was reinforced by further analyses (**Fig. 8D,E,F**). Mean classification accuracy using NP Ultra data, but not NP 1.0-like data, increased significantly when including footprint as a feature relative to performance without footprint (p = 0.0003). Classification of PV, SST, VIP, and FS_L_ improved when footprint was included and by a greater margin when compared to NP 1.0-like classification (**Fig. 8F**). Next, we iteratively removed each feature and assessed the difference in overall performance, a measure of feature importance (**Fig. 8G**). Doing so revealed that removing footprint impacted classification performance as much as spike duration (mean accuracy ± S.D. = 0.61 ± 0.04 for each) and more than the majority of other single-channel features, with the exception of average inter-spike interval (0.57 ± 0.05). Classifier accuracy was still above chance levels even when only decoding from footprint, average ISI, and baseline FR (**Fig. 8G**, inset; mean accuracy = 0.38 ± 0.03). In summary, we demonstrate that NP Ultra probes offer an enhanced ability to discriminate cell classes recorded from the visual cortex compared to classifiers using NP 1.0-like data, and that they do so in particular because of the high resolution spatial information captured by NP Ultra.

## Discussion

Here, we introduce Neuropixels Ultra, a device capable of recording extracellular neural activity with an unprecedented site density (1.3 sites/μm). These new probes have small site size and spacing, resulting in a trade-off in recording span compared to NP 1.0 probes (288 μm vs. 3840 µm vertical span), but allowing the detailed spatial structure of electrical fields to be sampled with unprecedented resolution. Therefore, NP Ultra probes are particularly suitable for targeting thin layers or small structures, such as individual cortical layers, the CA1 pyramidal layer, or subcortical nuclei including the claustrum and parts of thalamus, basal ganglia, midbrain, and hindbrain. By harnessing this improved spatial resolution for recording extracellular data and employing tailored spike sorting approaches, we demonstrate significant improvements in extracellular data quality, including higher spike amplitudes and improved signal-to-noise ratios, resulting in increased yield of functionally responsive neurons. Recordings made with NP Ultra probes capture subcellular features – axonal and dendritic signals – and provide a new window into the biophysics of action potential initiation and propagation. Finally, recordings at this density offer enhanced discrimination between cell classes, in particular for interneuron cell classes within the visual cortex. These devices therefore enable high-resolution measurements across multiple brain regions and species, and the resulting datasets will provide a useful resource for spike sorting algorithm development, for electrode array design, and for biophysical modeling.

We observed higher spike amplitudes and improved neuron yield with NP Ultra relative to previous probes. Larger amplitudes can be attributed to the interaction of two factors: 1) higher spatial sampling, which increases the likelihood of recording closer to the site of action potential initiation (Fiáth et al., 2018), and 2) smaller recording sites average signals over a smaller area, avoiding underestimation of peak amplitudes as much as large sites do (Canakci et al., 2017; Viswam et al., 2017; Hill et al., 2018). Additionally, NP Ultra probes exhibited a significantly higher signal-to-noise ratio compared to lower density probes. Although smaller sites incur a penalty in noise level, the increased site density of NP Ultra allows for the combination of information from multiple sites, effectively reducing the averaged noise level. Finally, NP Ultra has more precise localization of spikes for improved discrimination of nearby units. Combined, these benefits in spike sorting resulted in NP Ultra having a consistently higher yield of stably firing neurons with reliable visual responses recorded from mouse visual cortex compared to NP 2.0 and NP 1.0. Much of this improvement in yield results from an increased ability with NP Ultra probes to capture low amplitude spikes, which would have undetectably low SNR on probes with lower spatial densities. Importantly, the improvement in the yield of well-isolated neurons passing rigorous quality metrics (International Brain Laboratory et al., 2023a) was more modest (Ultra vs. 1.0, 159%) than the improvement in yield of all visually responsive neurons (Ultra vs 1.0, 305%). Though many of the extra neurons may not be well-isolated, they would nevertheless be useful for analyses such as dimensionality reduction or decoding (Trautmann et al., 2019; Zhang et al., 2024).

Recordings obtained from various brain regions in four species unveiled a remarkable diversity of extracellular neural waveforms. Particularly noteworthy was the presence of a population of “small footprint” units detected using NP Ultra probes across brain regions in different species, operationally defined here as units with an electrical waveform extent of less than 20 μm (smaller than the smallest distance between contacts on an NP 1.0 probe). In the mouse visual cortex, we observed that recordings from identified axons all exhibit small footprints. Despite the commonly accepted wisdom that axonal spikes have early positive peaks (Cooper et al., 1969; Deligkaris et al., 2016; Someck et al., 2023), we observed a fraction of axonal waveforms lacking such positive peaks, consistent with simulations of recordings from nodes of Ranvier in myelinated axons (Halnes et al., 2024). Our high resolution recordings therefore reveal that the spatial footprint, rather than a particular peak direction, is a more reliable signature of axonal recordings. Moreover, while recordings of axonal spikes have been reported with Neuropixels probes (Schröder et al., 2020; Sibille et al., 2022) and with other devices (Merrill et al., 1978; Goldberg and Fee, 2012; Robbins et al., 2013; Barthó et al., 2014; Sun et al., 2021), it has been assumed that this is only possible in limited situations when axons have particularly favorable properties (McColgan et al., 2017; Barthó, 2021). Here, we instead demonstrate these putative axonal recordings in all 15 brain regions and 4 species that we tested, establishing that NP Ultra probes can reliably achieve these recordings. Careful confirmation of the identity of these units will be required in each brain area in the future.

In mouse dentate gyrus (DG) and fish cerebellum (CB), we observed a significant fraction of small footprint units that might represent action potentials from small neuron types, axons, or both. Both DG and CB are notable for their populations of granule cells with very small somata. Considering DG granule cells, however, previous studies have indicated that they are relatively quiescent (Senzai and Buzsáki, 2017; Neubrandt et al., 2018; McHugh et al., 2022), firing at frequencies of about 0.15 spikes per second on average *in vivo* during wakefulness. The short duration of our DG recordings (approximately 5 minutes) would make it difficult to isolate such sparsely firing neurons and might have resulted in failure to record this cell class in general, regardless of the size of its extracellular action potential. Furthermore, we note that the majority of units recorded from the dentate gyrus were fast-spiking, an attribute not typically associated with mature granule cells (Pedroni et al., 2014), but which we observed in the small proportion of small footprint units in regions such as visual cortex and in small footprint units of corpus callosum. Therefore, while it appears plausible that our recordings here represent extracellular measurements of granule cells in DG and CB, this cannot be decisively concluded without further ground-truth experiments, such as recording genetically identified granule cells via opto-tagging.

In the visual cortex, the narrow action potential waveform (≤0.4 ms) spikes that we observed for most small-footprint neurons has traditionally been associated with putatively PV-expressing fast-spiking (FS) inhibitory interneurons (McCormick et al., 1985; Barthó et al., 2004; Kvitsiani et al., 2013). We discovered that small-footprint fast-spiking units (FS_S_) made up only a small fraction of optotagged units from genetically identified inhibitory subclasses – PV, SST, and VIP neurons – and that large footprint fast-spiking units (FS_L_) corresponded most closely to PV+ optotagged neurons. Indeed, more than half of all small FS cells were not associated with PV, SST or VIP tagged neurons. That is, FS_S_ units constitute a small, yet significant portion of the total units and almost one-third of all fast-spiking units in NP 1.0 data. In combination, these results suggest that FS_S_ units most likely represent axon segments of both excitatory and inhibitory neurons and are distinct from genetically identified PV+ fast-spiking units. Reliable identification of putative PV+ interneurons therefore requires considering both waveform duration and footprint, without which approximately one third of those units identified by waveform duration alone may be misidentified.

Our ability to decode cell class identity was significantly improved using NP Ultra compared to 1.0 probes. Nevertheless, our findings are applicable to existing recordings made with previous probes, enabling the identification of recorded units as PV+, VIP+, SST+, or other based on single-channel waveforms with a reasonable level of accuracy. In fact, our category of ’untagged’ units includes neurons from the PV, VIP, and SST cell classes, which likely leads to an underestimation of decoding accuracy (i.e., some untagged units labeled by the classifier as VIP might in fact be VIP+, though this is counted as an error). In the future, unit classification using a variety of additional metrics, such as non-linear waveform features (Lee et al., 2021), autocorrelograms (Barthó et al., 2004; Yamin et al., 2013; Senzai and Buzsáki, 2017), and oscillatory phase locking (Reifenstein et al., 2016), as well as alternative machine learning approaches (Crawford and Pineau, 2019; Tymochko et al., 2020; Kingma and Welling, 2022; Beau et al., 2024; Lee et al., 2024), may yield even greater performance.

Taken together, our findings collectively highlight the advantages of electrophysiological probes with increased site density for a wide range of important neuroscience applications. While sacrificing recording span, Neuropixels Ultra probes offer improved signal quality, making them ideal for recording from small neuron types and subcellular compartments, maximizing yield in dense brain regions, and accurately distinguishing cell classes of recorded neurons.

## Methods

### Probe noise model

When using neural probes with planar microelectrodes, the quality of the neural recording (e.g. signal to noise ratio) will depend on several factors, including the electrode size; the electrode impedance and the input impedance of the recording amplifier; the electrode noise and the noise of the recording amplifier; and finally the distance and alignment between the neuron and the electrode. These factors contribute to different signal-degradation effects that are explained as follows.

#### Signal attenuation due to electrode impedance

The electrode impedance is inversely proportional to the electrode size. Small electrodes can exhibit larger signal attenuation due to the impedance ratio (i.e. voltage divider) at the input of the amplifier. This attenuation is given by: *Z_elec_/Z_amp_*. In the Neuropixels 1.0 design, *Z_amp_* corresponds to 27.2 MΩ (at 1 kHz). Thus, the signal attenuation can be calculated as:

Although the signal attenuation is increased by almost 5 times, it continues to be negligible due to the much higher input impedance of the recording amplifier.

#### Noise performance

The total noise affecting neural recording has two components: i) the thermal noise generated by the electrode-tissue interface (*V_n-elec_*) and ii) the noise of the readout electronics (*V_n-amp_*). The total noise can be calculated as:

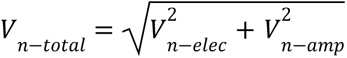

The noise generated by the electrode-tissue (or electrode-electrolyte) mostly depends on the electrode area, the double-layer capacitance formed at the electrode-electrolyte interface (i.e. electrode impedance) and the resistivity of the saline solution, medium or tissue (Yang et al., 2009; Lopez et al., 2012; Viswam et al., 2017).

In the Neuropixels 1.0 design, *V_n-amp_* corresponds to 5.4 µV_rms_ (for the action potential band: 300 Hz to 10 kHz). For the low-impedance TiN electrodes, the electrode noise was modeled and measured in saline as 2.7 µV_rms_. Based on estimated electrode impedance values, it is possible to predict the noise performance of smaller electrodes for different solution/tissue resistivities (Lopez et al., 2012). In saline, the total noise can be calculated as:

If we consider the resistivity of the brain tissue, which is at least 4 times higher than the resistivity of saline, the total noise can be calculated as:

Note that in Neuropixels 1.0, the noise is dominated by the readout-electronics noise, while in Neuropixels-Ultra the electrode noise has a bigger impact on the total noise.

Although the electrode noise increases by a factor of ∼2, the increase of the total recording noise is only 23% in saline or 32% in brain tissue. This increase is modest relative to the amplitude of ‘biological’ noise (i.e. spiking activity of distant neurons). This indicates that, thanks to the low-impedance of the TiN electrode material, the reduction in electrode size does not significantly impact the quality of the recordings.

### Noise and gain characterization

*In vitro* noise measurements are performed in the standard self-referenced configuration as described in the Neuropixels manual, with the reference and ground connected together and to a Pt wire electrode in the saline bath. Noise on each channel is measured by averaging Fourier power spectra from 5x 3-second-long sections of data, and estimating rms from the integral over 300-10000 Hz:

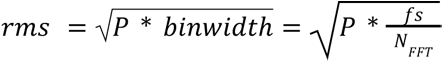

Where P is the sum over the frequency range in the power spectrum, F_s_ is the sampling frequency, and N_FFT_ is the number of points in the FFT calculation.

Noise is estimated from *in vivo* data for “low activity” channels with event rates < 0.01 Hz using (1.4826*median absolute deviation). For the gain measurement, the probe ground and reference are connected to the Faraday cage ground, and a 2 mV, 3 kHz sine wave is applied to the saline bath through a Pt wire electrode. The amplitude of the sine wave on each channel is measured by averaging over a 15 Hz window about the 3 kHz peak in the Fourier power spectrum.

### Crosstalk estimation

To estimate the crosstalk between channels, we took advantage of our *in vivo* recordings which provide a situation in which a small voltage source is located immediately adjacent to the probe. We reasoned that for neurons with spatially restricted extracellular action potentials, distant sites on the probe should not reflect the waveform of the neuron unless by crosstalk. To the extent that this assumption might not be true (i.e. distant sites really do detect the actual extracellular action potential), we would measure signal at the distant site and incorrectly attribute it to crosstalk; therefore, this estimation strategy produces an upper bound on the true amount of crosstalk.

Specifically, we selected units for which at most 20 sites had significant amplitude of the average waveform (defined as peak-trough amplitude > 25% of the amplitude on the peak site), and averaged the amplitude at the 10 most physically distant sites relative to the peak site. The amplitude on the distant sites was calculated using the same peak and trough timepoints as the peak channel, since any crosstalk should be temporally instantaneous. We report the ratio between this average amplitude on the distant 10 sites and the amplitude on the peak site. This measurement was subject to biological noise, but across an average of n=519 small neurons, we measured 0.0007% ± 0.0736% (mean ± s.e.m.; **Supp. Fig. S4**).

### Light artifact tests

We tested the sensitivity of NP Ultra probes to 470 nm light (the wavelength used for channelrhodopsin activation). We used a fiber-coupled laser (200 µm core fiber) to shine light directly on probes submerged in phosphate-buffered saline. Light was incident on the front side of the probes. During the tests, we simultaneously recorded from two NP Ultra probes (passive, non-switchable version) and one NP 1.0 probe to compare light artifacts across probe variants. Two different stimulation waveforms were tested (10 ms pulse and 1 s raised cosine stimulus) at three light intensities (0.5, 4.0, 10.0 mW/mm2). We quantified photo-artifacts in both the spike (0.3-10 kHz) and LFP bands (0.5-500 Hz) by computing trial-averaged photo-evoked responses (100 trials) time-locked to the start of photo-stimulation (**Supp. Fig. S5**). Artifact magnitudes were computed as the absolute value of the light-evoked voltage deflection peak during the photo-stimulation period.

### Recordings from diverse brain regions in the mouse

#### At UW

All experimental protocols were conducted according to US National Institutes of Health guidelines for animal research and approved by the Institutional Animal Care and Use Committee at the University of Washington.

For awake, head-fixed acute recordings, wild type C57BL/6 mice of both sexes, between 2 and 8 months of age, were first implanted with custom-made steel headplates and 3D-printed plastic recording chambers. Following recovery, mice were acclimated to head-fixation for at least two sessions before recording. Head-fixation mice were seated on a plastic body restraint tube with forepaws on a rotating rubber wheel. Three iPad screens were positioned around the mouse at right angles. On or before the first day of recording, a 1-2 mm diameter craniotomy with intact dura was prepared with a dental drill over the target brain area under anesthesia. The craniotomy was either protected with removable silicone sealant (Kwik-Cast, World Precision Instrument) or covered with transparent dura-gel (Dowsil 3-4680) with a further protective plastic cap sitting on top of the recording chamber, after the surgery and before recording procedure starts. After several hours of recovery, mice were head-fixed in the recording setup. We used single-ended configuration for all recordings, with ground and reference pins on the probe tied together. We used either external or internal reference methods. For recordings with external referencing, an Ag wire was connected to the probe ground/reference wire and positioned above the skull. After peeling off the silicone sealant, the craniotomy and Ag wire were then submerged in a bath of Ringer’s lactate solution. For recordings with internal reference, a probe was directly inserted through the dura gel without solution bath and the reference site on the probe tip was used instead. Prior to each insertion, the electrode shank was coated with CM-Dil (Invitrogen), a red lipophilic dye, for later histological reconstruction of probe tracks.

To sample waveforms from diverse brain areas in a single penetration, we automated the advancement of probes through manipulators (Sensapex Inc., uMP-4) using the Sensapex API via Matlab, following every 5 minutes of stable recordings. For each step, we advanced the probe by 300 μm at 5 μm/sec speed, followed by 50 μm retraction, to help reduce the tissue compression caused by the probe. Recordings normally stabilized within one minute after probe motion, as visualized by post-processing drift maps. We typically repeated this for 8-16 steps from the surface of the penetration, depending on the final depth of target brain area. For most sessions, we recorded with two probes simultaneously on two hemispheres in a single session.

After recording, mice were perfused with 4% paraformaldehyde. The brain was extracted and then fixed in 4% paraformaldehyde for a further 24 hours at least at 4 °C. We cleared the brain tissue with iDISCO before 3D imaging with lightsheet microscopy (UltraMicroscope II, LaVision BioTec). We used 561 nm channel for imaging probe track with DiI and 488 nm channel for brain autofluorescence, with 10 μm isotropic resolution. We registered the imaging volume to Allen Mouse Brain Common Coordinate Frameworks (CCF) Atlas with ARA tools (https://github.com/SainsburyWellcomeCentre/ara_tools), which calls Elastix via a Matlab wrapper. Probe tracks were traced in the atlas-registered brain volume with open-source package Lasagna (https://github.com/SainsburyWellcomeCentre/lasagna). While the DiI track provides a good estimate of the probe trajectory, the depth information between DiI track and electrophysiological recordings usually do not match with high precision, due to the warping and shrinkage of the brain sample during perfusion and tissue clearing steps, and due to uncertainty about the final probe tip location along the track. Therefore, we used multiple electrophysiological landmarks (e.g., cortical layer 5 neuron layer, CA1 pyramidal layer, white matter boundaries, etc) to match the recording sites with the probe trajectory by linear interpolation (Steinmetz et al., 2019). After this manual curation process, we assigned each probe site and therefore each spike cluster to designated CCF coordinates.

The NP Ultra brain-wide data was sorted with Kilosort 2.0, using a 96 site template. The timing of all recording epochs were extracted from a manipulator motion start/stop channel synchronized to the electrophysiology data. The first minute of the recording after manipulator motion was discarded due to probe-tissue motion, leaving 4 minutes of stable recording data for each epoch. Kilosort was then batch run based on epoch timing, and units were selected with an automated quality metric within Kilosort 2.0 after spike sorting.

#### At Janelia

Acute recordings were conducted at the HHMI Janelia Research Campus, following guidelines set by the Institutional Animal Care and Use Committee. Three male VGAT-ChR2-EYFP mice, aged 6 months, underwent a brief stereotaxic surgery to implant a titanium headpost and build a recording chamber using dental cement for acute head-fixed recordings. Postoperative analgesia was provided using Buprenorphine (0.1 mg/kg, intraperitoneal injection), and Ketoprofen (5 mg/kg, subcutaneous injection) was used at the time of surgery and postoperatively for two days. Following surgery, mice were habituated under head restraint in the recording setup for two days with incremental acclimatization duration (30 minutes to one hour).

On the first day of recording, two to four craniotomies of diameter 1-1.5mm were made with a dental drill over the target areas of interest under anesthesia, while leaving the dura intact. The craniotomies were protected with removable silicone sealant (Kwik-Cast, World Precision Instrument) sitting on top of the recording chamber between recording sessions. After a three-hour recovery period, the animal was head-fixed in the recording setup, and two or four NP Ultra probes were inserted into separate craniotomies and to the target depths, inferred from manipulator readings. The insertions covered brain regions such as the cortex, striatum, thalamus, midbrain, cerebellum, and medulla. The tip of the electrode was coated with CM-Dil (Invitrogen), a red fixable lipophilic dye, before each insertion to enable histological reconstruction of probe tracks. Probes were slowly lowered at a speed of 10 μm/s to the first intended depth using micromanipulators (uMP-4, Sensapex, Inc), and after reaching the desired depth, the inserted probes were allowed to settle for approximately 5 minutes before recording commenced. Multiple probes were recorded simultaneously for 20-30 minutes. Since the NP Ultra sites span a length of 288 μm only, after each recording, probes were further inserted by 300 μm to make a successive recording. As such, over the course of a session, probes were advanced 2-6 times to collect data at multiple depths from each penetration, and brain tissue was allowed to relax for a few minutes after each probe advancement. All probes had a wire soldered onto the reference pad, which was shorted to the ground pad. This wire was then connected to an Ag/AgCl wire positioned above the skull. During recordings, the craniotomies and reference wire were submerged in cortex buffer (NaCl 125 mM, KCl 5 mM, Glucose 10 mM, HEPES 10 mM, CaCl_2_ 2 mM, MgSO_4_ 2 mM, pH 7.4). Daily recording sessions lasted approximately two hours and were repeated for multiple consecutive days in a craniotomy, with each penetration separated by at least 200 μm at the point of entry. All recordings were made with open-source software SpikeGLX (http://billkarsh.github.io/SpikeGLX/) in external or tip reference mode, and the animal was awake and sitting quietly in a body restraint tube throughout the recording sessions. The dataset was sorted offline with Kilosort 2.0 using a 96 site template.

To visualize the recording locations, the mice were perfused transcardially with PBS followed by 4% PFA, and their brains were fixed overnight and then cleared with SPiB solution using an aqueous-based clearing method called ’Uniclear’ to facilitate whole-brain dilapidation and refractive index matching (Chen and Svoboda, 2020). The cleared brains were then imaged with a light sheet microscope (Zeiss Z1) using a 5x objective with a voxel size of 1.2 × 1.2 × 6 μm. Whole-brain autofluorescence was captured with a 488 nm channel, while Dil tracks were imaged with a 561 nm channel. The resulting tiled images were stitched together using IMARIS software, and the entire brain stacks were registered to the Allen Institute Common Coordinate Framework (CCFv3) of the mouse brain based on anatomical landmarks, allowing each spike cluster to be assigned to a specific CCF coordinate.

Data was preprocessed using CatGT (https://billkarsh.github.io/SpikeGLX/#catgt) for multiplex correction, filtering, artifact removal, and common average subtraction. Spike sorting was performed with Kilosort 2.0, followed by manual curation. We classify units as either ‘unimodal’ or ‘overlapped’. Unimodal units have distributions in feature space visually that appear to come from a single distribution that does not overlap other units, and refractory period violations are low. ‘Overlapped’ units have distributions in feature space that overlap other units, and generally have higher refractory period violations. In the analysis of signal and physical distance, all units with > 500 spikes are included, to provide as complete a picture of the activity as possible. For the waveform shape analysis, only ‘unimodal’ units are included, so that mean waveforms are as close as possible to single units.

#### At University College London

All experimental procedures were carried out under license from the UK Home Office in accordance with the UK Animals (Scientific Procedures) Act (1986). Acute recordings were performed on adult (>P60) Rbp4-Cre male and female mice kept on a 12 hours dark/light cycle. At the time of surgery, mice were subcutaneously injected with Rimadyl (5mg/ml) and Lidocaine (5mg/ml) to numb the area above the skull prior to headplate implantation. A 0.5 g customized metal headplate was attached to the skull using super glue and dental acrylic (Super bond C&B polymer mixed with Super bond monomer and catalyst). After headplate implantation, mice were allowed to recover at 37 °C inside an incubator and were closely monitored for 5 days post-surgery. On the first day of recording, mice were administered dexamethasone (Dexadreson, 2 mg/ml, intramuscular injection) to reduce brain swelling. On the first day of recording, one 2 mm-diameter craniotomy was made over the region of interest under anesthesia using a surgical motorized drill. The skull flap was carefully removed to leave the dura mater intact and the exposed brain surface was cleaned using IVE and subsequently protected with transparent Duragel and removable silicone sealant (Kwik-Cast, World Precision Instrument) placed on top of the recording chamber. Finally, the animal was allowed to recover for 3-hours before starting any further experimental procedure.

Before recording sessions, mice were acclimated to the experimental rig for at least 3 days: animals were head-fixed above a static 3D-printed holder using a head bar clamp, and habituated to head restraint of increasing duration (starting from 10 minutes and up to 1h). On recording days, the shank of Neuropixels probes was covered with red fluorescent dye DiI (Invitrogen) to allow for histological probe tracing. Then the probe was inserted perpendicularly to the cortical surface using a micromanipulator (Sensapex uMP-4). Probe insertion speed was kept at 20 µm/s to minimize cortical damage, terminating at 2000-2800 µm below the brain surface for NP 1.0 and 1200 µm for NP Ultra. Probes were allowed to settle for about 10 minutes before the start of every recording. All recordings were carried out using the acquisition software SpikeGLX (http://billkarsh.github.io/SpikeGLX/) using tip reference mode, and lasting about 1.5-2 hours. Every animal was recorded for a maximum of three consecutive days, with recordings taking place at different locations within the craniotomy.

All recordings were processed to align channel sampling using the common average referencing software CatGT (https://billkarsh.github.io/SpikeGLX/#catgt); putative neuronal units were spike sorted with Kilosort 2.0 (https://github.com/MouseLand/Kilosort2) and Phy2 (https://github.com/cortex-lab/phy) , and manually curated in Phy2. For every unit, we computed spike-triggered voltage maps showing the extracellular signature of each action potential across time and channels. We then averaged 2000 of these maps to compute a template spike voltage map for each unit. We located the putative soma as the channel with the largest sink recorded at the time of spike in the template maps.

To correct for vertical probe motion during the recording we used a two-step strategy. First, we used Kilosort 3.0 drift estimation to correct for slow probe movements at the second scale. Second, to correct for fast movements on a spike-by-spike basis, we used 1D cross correlation to compute the vertical shifts needed to register spike-triggered voltage maps to a target reference map obtained from 1000 long-ISI spikes selected randomly throughout the recording.

To identify units with significant dendritic back-propagation, e.g. those for which the apical dendrite was best aligned with the probe, we used a permutation test. Starting from the soma, we compared the average spike amplitude of every channel in the template spike map to a null distribution of amplitudes obtained from 500 shuffled template maps, each computed from a random linear shift of the spike times. Moving from towards the cortical surface, we considered as significant dendritic recordings those channels bearing signals larger than 95th percentile of the null distribution. We included for further analysis neurons which displayed significant dendritic back-propagating signals extending beyond 250 µm from the soma.

### Muscimol experiments

To verify the identities of small footprint units in the mouse cortex, we topically administered muscimol to the surface of the dura through the craniotomy, while acutely recording cortical activity with NP ultra probes in head-fixed mice. Acting as a GABA agonist, muscimol will silence neural activity of local cortical neurons, but leaving activity of long range axons from elsewhere intact.

Headplate implant surgery, craniotomy surgery and acute NP ultra recordings were performed mostly as described in the Methods “Recording from diverse brain regions in the mouse (At UW)” section, with additional details as below. During the craniotomy surgery procedure, a cement well was built around the craniotomy after the craniotomy was made and covered with removable silicone sealant, to constrain the spatial spread of drug application within the craniotomy during recording. For acute NP ultra recordings, the craniotomy was submerged in a bath of Ringer’s lactate solution during control conditions. After a period of baseline spontaneous activity recording for around 10 minutes, a set of 5-minute sparse noise stimuli were presented on the iPad screens for receptive field mapping.Thereafter, 5mM muscimol in Ringer’s solution was applied to the craniotomy and “bathed” the surface of the brain for the remainder duration of the recording. The majority of unit activity were silenced, 3-10 minutes after the muscimol application. A second set of 5-minute sparse noise stimuli were then presented for receptive field mapping for units that survived muscimol.

### Spike sorting methods

To take advantage of the high spatial resolution of NP Ultra sites, we utilize a spike sorting pipeline, DARTsort (Boussard et al., 2023), that takes advantage of recent progress in spatial localization of spikes. After an initial detection step, we denoise (Lee et al., 2020) and localize the spikes (Boussard et al., 2021), and then estimate the motion of the probe (i.e., drift) relative to the recorded neurons. For this registration step we use the decentralized approach from (Windolf et al., 2023), which is more robust to boundary effects (i.e., neurons moving on or off of the short NP Ultra probe) than the template-based registration method used in Kilosort2.5 (see (Windolf et al., 2023) for further analysis). After registration, we have a robust estimate of each spike’s amplitude and position relative to the probe. Then, by subtracting the estimated drift from the position of each spike, we obtain a set of 3D features for each unit: registered 2D position and spike amplitude (Hilgen et al., 2017). We cluster in this 3D feature space using HDBSCAN (Malzer and Baum, 2019), and then recursively re-split these clusters using additional features computed by principal components analysis applied within each cluster (Swindale and Spacek, 2014). Even after registration, we find that some units have some remaining drift (changes in mean registered location and amplitude) over time. Therefore we divide the recording into 5-minute segments that we separately cluster and then merge, to make the clustering step more robust to these individual-unit drift effects.

Next we run a template-matching deconvolution step to resolve “collided” spikes that overlap temporally and spatially. To enable drift-aware template matching, we construct a set of denoised “super-resolution” templates for each unit by dividing the depth of the probe into 2 µm bins and then averaging the spikes localized to each bin separately. During template matching, we can then shift these super-resolved templates according to the drift to efficiently detect spikes for each unit. One major benefit of this approach is that we do not need to apply a shift/interpolation step to the raw data (as in Kilosort 2.5); in practice, we found that this raw-data interpolation removed signal near the edge of the probe due to boundary effects, which are particularly problematic for the small dense NP Ultra probes. In our approach, instead of shifting the raw data, we shift the templates, allowing us to robustly track neurons moving on or off the probe.

As the initial clustering is usually imperfect, we perform the above steps iteratively: cluster, then deconvolve, then re-cluster on the deconvolution output. We use the post-deconvolution collision-subtraction method from (Lee et al., 2020) to denoise the spikes obtained in the deconvolution step, to enable improved iterative clustering.

### Methods for site pattern comparison using spatial resampling

We resampled data recorded with NP Ultra to predict what signals would have been recorded from the same brain location and time period with a lower density probe. To do this, we simply average together the four 5×5 µm NP Ultra sites that are co-localized with a single 12×12 µm site in the resampled pattern (**Fig. 2E**; due to the 1 µm gaps between sites in NP Ultra, the four 5×5 sites fill the 12×12 µm area almost exactly, with small missing gaps). We used this method to measure the expected changes in amplitude and waveform shape for the Neuropixels 1.0 and 2.0 patterns, and a “large dense” pattern of 12×12 μm sites with 12 μm pitch (**Fig. 2G-K**, **Fig. 3D-G**, **Fig. 8D,F**).

For measurements of the site size and pattern dependence that include the effects of noise - specifically, for computing the ‘Template SNR’ (**Fig. 2I**) and for analyses involving spike sorting the resampled raw data (**Fig. 3D-G**) - we needed to create data with realistic noise and waveform variation, which in real data combines electronic noise with signals from distant units.

Averaging the four raw traces together as described above reduces the noise by 1/ 2; to boost the noise back to normal levels, simulated noise with a matched frequency spectrum is added to the averaged traces (code at: https://github.com/jenniferColonell/NP_Ultra_downsample).

An interpolation method was used for the analyses on waveform shape and peak amplitude in **Supp. Figs 8, 9**. The mean waveform from the Ultra data was used to create a gridded interpolant (MATLAB, using ‘makima’ interpolation). The waveforms for an arbitrary pattern with any site size and position can then be estimated by averaging together interpolated waveforms at sets of points that span the model sites. This method has the advantage of allowing a test site pattern that is slightly better matched to the true NP 1.0 and NP 2.0 geometry than in **Fig. 2E**.

### Analyses of sorting quality

Spike amplitude was calculated as peak-to-peak voltage, i.e. the maximum voltage of the waveform minus the minimum (**Fig. 2G-H; Supp. Fig. S8A, S9A**).

In template matching methods, a multi-site template is convolved with the filtered data to detect spikes. To quantify the impact of site density on detectability in template matching, we calculate the ‘Template Matching Signal to Noise Ratio’ for each unit (**Fig. 2I; Supp. Fig. S8B, S9B**) defined as:

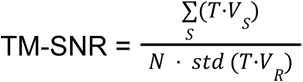

Where T is the template vector of 40 time samples * number of sites in the template, and V is measured voltage on those sites centered either at a spike time (to measure signal) or a randomly chosen time (to measure noise). S indexes spike times of the unit, R indexes random times, and N is the spike count of the unit. This is an estimate of the signal to noise ratio in data filtered by the template. Spike times for each unit are drawn from the sorted Ultra data, and the TM-SNR is calculated for the Ultra data and spatially resampled data obtained from averaging sites (see exact patterns in **Fig. 2E**).

The spatial spread of spikes was calculated as the standard deviation of the x-coordinates (lateral dimension, across the face of the probe) or of the z-coordinates (axial dimension, down the length of the probe) across all spikes in a given unit (**Fig. 2J-K**). For the resampled patterns, these coordinates were computed by resampling each spike individually and then computing that spike’s estimated coordinates using the same method employed in the main DARTsort pipeline.

To understand the possible effects of this spatial scatter in more detail, we computed the average distances between neurons, as well as the minimum distance that each site pattern should be able to resolve, in a simple simulation (**Supp. Fig. S8C, S9C**). We use a Monte Carlo model to generate sets of peak-to-peak voltages over sites for a point source with 1/R amplitude falloff, placed 15 μm from the probe, with peak amplitude at the probe of 50 μV and rms noise of 14 μV. For each set, all sites within 60 μm of the peak site are fit to obtain an estimated XZ position (Boussard et al., 2021). Modeled noise in the position estimate is the standard deviation of the Monte Carlo trials, and is 5 μm for a NP Ultra probe, and 25 μm for the NP 2.0 pattern. The increased accuracy is due to the inclusion of more sites in the 60 μm radius of fit points, which decreases the impact of noise. These values are the upper limit of position resolution, and can be compared to the measured distribution of nearest neighbor unit distances from the Ultra data. Units with nearest neighbor distances below the spatial resolution of the probe will be more difficult to sort. The experimental nearest neighbor distributions are calculated including both ‘unimodal’ and ‘overlapped’ units to build as complete a set as possible to assess the potential impact of this difference in spatial resolution on sorting. Note that standard deviations of positions in experimental data can be larger (see **Fig. 2J-K**); this is likely due to variability in the spikes (background firing, incomplete drift correction) not captured in the simple model.

To measure how more detailed spatio-temporal waveforms could affect sorting, we created a waveform distance metric and compared the waveform distances measured at varying site density (**Supp. Fig. S8D, S9D**). Higher site density will always provide more detail, and better confirmation of small differences, but the engineering question is whether the waveforms are ‘mostly’ smoothly varying and distinguishable with sparse sampling. To isolate the waveform distance from difference in z position and amplitude, a neighborhood of rows centered about the peak row is taken for each template, and the amplitude is normalized. For the spatially resampled patterns, the expected waveforms are calculated using interpolation of the full density waveform, with the sparser pattern centered at the position of the peak row for each template. Centering the sparse pattern on the peak row ensures the two templates look as similar as possible, by ensuring that the same part of the footprint is sampled.

A simple metric to compare the two normalized templates [A,B] is the L1 norm of the difference between the two divided by the sum of the L1 norm of both waveforms, summed over sites, that is:

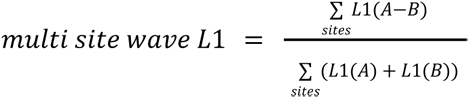

This quantity is essentially the fraction of detected signal that is different between the two templates. It is a characterization of the waveform shape that does not account for noise in individual spikes.

### Yield and stability testing with imposed probe-brain motion

#### Recordings

To make recordings with imposed probe motion in head-fixed mice (n=2 mice, n=6 sessions), we automated the probe motion through manipulators (Sensapex Inc., uMP-4) using the Sensapex API via Matlab. Manipulators were programmed to alternate between moving forward and backward along the probe axis, for a total of 10 steps forward and 9 alternating steps backwards with 25 μm travel in each direction at 1 µm/sec speed, which results in 25 µm net total forward displacement disposition of the probe by the end of imposed motion. A digital synchronization signal was issued at the start and end of each motion step for later alignment with neural data. To match neuron identities before and after imposed motion, a battery of 118 natural images with 60 repetitions per image were presented to ‘fingerprint’ visual responses. Images were presented in a random order. An average of 30 repetitions of the image set were presented before imposed motion, and an average of 15 repetitions were presented after imposed motion. The duration of each image follows a randomized exponential distribution (mean: 0.5 seconds; minimum: 0.2 seconds; maximum: 3 seconds). NP Ultra probes were inserted into the visual cortex at 700-900 µm depth to sample deep layer neurons.

The NP Ultra imposed motion dataset was spike-sorted with DARTsort (see “Spike Sorting Methods”) (Boussard et al., 2023). The original full pattern data, as well as the same data resampled to NP 1.0-like, NP 2.0-like, and 12 μm large dense site patterns (see “Methods for pattern comparison using spatial resampling”, **Fig. 2E**), were sorted. The timing of imposed motion epochs was extracted from a manipulator motion start/stop channel synchronized to the electrophysiology data.

#### Yield and stability analysis

To assess and compare the yield and tracking stability of recorded neurons across different channel densities (full density NP ultra, NP 1.0-like, NP 2.0-like, and 12 μm large dense), visual response fingerprints were created for each unit by concatenating average PSTHs (window 0 to +250 ms aligned to image onset, bin size 10 ms) for all 118 images into a single vector. PSTHs were normalized for a given unit by subtracting its baseline activity (-100 to 0 ms) averaged across trials prior to image onset. PSTHs were denoised by reconstructing each PSTH from the top two principal components taken on the average PSTH per image × time points matrix for each unit (dimensions 118×35).

To determine whether a unit had a reliable visual fingerprint, a null distribution for each unit was created by randomly permuting the image identities before concatenating the PSTHs into a vector (**Fig. 3C and Supp. Fig. S12A**). This distribution preserves the noise and average PSTH time-course of a given unit while disrupting its selective visual response fingerprint. Average PSTHs were calculated from odd-numbered pre-motion trials and even-numbered pre-motion trials. Image identities were randomly permuted 1000 times, and odd-numbered pre-motion trials were used to create the null distribution. Units were considered to have a reliable visual response if their original correlation score (i.e. pre-motion odd vs. pre-motion even trials) was greater than all scores in their null distribution (i.e. 1000 pre-motion odd shuffles vs. pre-motion even trials). This ensures that included neurons have both a reliable fingerprint (i.e. responses are consistent between trials of the same image) and a selective fingerprint (i.e. responses to each image significantly vary).

To create a distribution of pre- vs. pre- and pre- vs. post-imposed motion fingerprint correlations (Pearson correlation coefficient), 1000 unique sets of visual fingerprint vectors were constructed for each neuron by randomly partitioning the image repetitions for each image before imposed motion into 2 non-overlapping subsets (15/30 trials in each subset, for all 118 images). In each of these 1000 sets, fingerprints from each pre-motion subset of trials (i.e. the average across trials) were correlated, and the fingerprint from one pre-motion subset was correlated with the fingerprint from the post-motion trials (**Supp. Fig. S12B-C**). The fingerprint stability ratio for a given neuron was calculated by dividing the mean of the 1000 pre- vs. post-motion correlations by the mean of the 1000 corresponding pre- vs. pre-correlations. To compare fingerprint stability tracking across the 4 site patterns, stability ratios for every neuron sorted from a given pattern were pooled across all sessions (**Fig. 3G**). Neurons were only included in this analysis if they had a reliable visual response and had a stable firing rate between the pre- and post-motion periods (neurons were dropped if the mean firing rate across all PSTHs in pre-motion period was >4x different than that of the post-motion period).

### Biophysical simulations

All neural simulations (**Supp. Figs. S7, S15, S16**) were done through LFPy 2.3 (Hagen et al., 2018) running on NEURON 8.1 (Carnevale and Hines, 2006). Action potentials were evoked by a step current injection (POINT_PROCESS in NEURON), where the amplitude of the current was adjusted for each cell model to evoke an action potential. Calculations of extracellular potentials were done with LFPy, assuming an electrode diameter of 5 µm, incorporated through the disc-electrode approximation. Simulations in **Supp. Fig. S7** used the rat layer 5 pyramidal cell model from (Hay et al., 2011). Simulations in **Supp. Fig. S16** used the rat layer 5 pyramidal cell from (Hallermann et al., 2012) (available from https://modeldb.science/144526), and the parameters for the passive and active conductances for the axon models used in **Supp. Fig. S15** were extracted from the same neuron model.

### Recordings from species other than mouse

#### Recordings from monkey

One macaca nemestrina weighing 6.6 kg (female, 9 yr) participated in the experiments which were conducted in the anesthetized, paralyzed preparation. At the start of the experiment the animal was placed in the stereotaxic instrument and then a craniotomy and a durotomy were performed to target the primary visual cortex (V1). Anesthetic and paralytic regimens are as described elsewhere (McLelland et al., 2015). All animal procedures conformed to National Institutes of Health guidelines and were approved by the Institutional Animal Care and Use Committee at the University of Washington.

The NP Ultra probe was advanced into the cortex using a hydraulic Microdrive (MO-97A, Narishige) which was mounted onto the stereotaxic arm. Once spike waveforms first appeared on the spikeGLX activity map, we retracted the probe 50 μm and mapped the receptive field (RF) of recorded neurons using achromatic bars under experimenter control. Then data collection on the main experiment began, during which we presented images of naturalistic objects (200 images from (Kiani et al., 2007)) repeatedly (presentation duration: 200 ms). Images were sized to cover mapped RFs.

As the data were continuously collected we completed the following sequence of operations 11 times: (1) probe at rest for 10 minutes, (2) advance probe 300 μm deeper over a 2 minute duration, (3) retract probe for 50 µm. This procedure allowed us to record spike waveforms at 11 different depths where every depth was 250 μm deeper than the previous recording site. Following completion of the recording session, we retracted the probe and confirmed that spikes completely disappeared at a depth within 200 μm from where spikes had first appeared during probe insertion.

Stimuli were presented on a liquid crystal display monitor (24 inches; 100-Hz frame rate; 1920×1080 pixel size, XL2430-B, BENQ) using custom experimental control software, Pype2 (Mazer, Jamie, 2013). Neural data were acquired using spikeGLX with internal tip reference.

The data were batch sorted based on epoch timing, with Kilosort 2.0 using a 96 site template, and units were selected with an automated quality metric within Kilosort 2.0 like recordings from other species.

#### Recordings from lizard

All experimental procedures were performed in accordance with German animal welfare guidelines: permit no: V54-19c20/15-F126/2006 delivered by the Regierungspraesidiµm Darmstadt (E. Simon).

A day before surgery, the lizard was administered analgesics (butorphanol, 0.5 mg kg−1 subcutaneously; meloxicam, 0.2 mg kg−1 subcutaneously) and antibiotics (marbofloxacin, marbocyl, 2 mg kg−1). On the day of surgery, anesthesia was initiated with isoflurane in an induction box and maintained with 1–4% isoflurane after intubation (Hallowell EMC anesthesia Workstation AWS). The lizard was placed in a stereotactic apparatus (Kopf 963) after ensuring deep anesthesia (absence of corneal reflex). Body temperature during surgery was maintained at 30 °C using a heating pad and esophageal temperature probe (Harvard apparatus part number 50-7213). Heart rate was monitored using an Ultrasonic Doppler flow detector (PARKS medical electronics, INC. model 811-B). The skin covering the skull was disinfected using a 10% povidone-iodine solution before removal with a scalpel. A fine layer of UV-cured glue (Oxford Flow Light Cure Flowable micro-hybrid composite A2) was applied around the exposed surfaces of the skull. A 3×2 mm craniotomy was then drilled around the parietal eye and the pericranium was removed. The parietal eye was retracted and fixed to the posterior edge of the craniotomy using histo-acryl tissue glue. The dura and arachnoid layers covering the forebrain were removed with fine forceps. A silver wire was inserted into the CSF fluid next to the olfactory tract, acting as both reference and ground. Pia was removed over the area of probe insertion (medial cortex). Prior to insertion, the probe was mounted on a movable drive (R2Drive, 3DNeuro), and secured to a stereotactic holder. The probe was stained with DiI (Invitrogen™, Vybrant™ DiI cell labeling solution) and implanted 500 µm deep in the medial cortex (MC) at a speed of around 100 µm/second.

After insertion of the probe, the brain was covered with Duragel (Cambridge Neurotech), and the craniotomy sealed with sterile vaseline. The microdrive base was fixed to the skull with UV glue and the microdrive was released from the stereotactic holder. Finally, a 3D-printed cap was mounted around the skull. The cap provided mechanical protection, carried the head-stage, as well as colored strips of tape for head-direction and position tracking. A silicone sealant (Kwik-Cast, World Precision Instruments) was applied around the edges of the cap and skin.

After surgery, the lizard was released from the stereotactic apparatus and left on a heating pad set to 30 °C until full recovery from anesthesia. On the days after the surgery, probes were slowly lowered into the tissue (up to 280 µm a day).

During recording sessions, lizards were freely moving in a 150 cm diameter circular arena, with two salient opposing visual cues in a brightly lit room. The room was heated to 29 °C, allowing lizards to stay active for an extended duration. Live prey items (mealworms) were dropped into the arena at random times and locations, motivating lizards to explore and maximize arena coverage.

After the end of the experiments, the lizard received intramuscular injections of ketamine and Dormicom, followed by induction and intubation with Isoflurane as described in the surgical method. The 3D printed cap, head-stage, and microdrive carrying the probe were retrieved. Lizard was then decapitated and immediately perfused through the carotid arteries with PBS, followed by 4% formaldehyde. The brain was removed and stored in 4% formaldehyde for one day, followed by an additional day in a 30% sucrose solution. The brain was frozen with dry ice and cut into 70 µm coronal slices. Slices were mounted on glass slides, and stained with DAPI. The final position of the tip of the probe was identified based on tissue damage and a DiI signal, imaged using a slide-scanner (Zeiss, Axio scan Z.7). The data were batch sorted with Kilosort 2.0 using a 96 site template, and units were selected with an automated quality metric within Kilosort 2.0 like recordings from other species.

#### Recordings from electric fish

All experiments adhered to the American Physiological Society’s Guiding Principles in the Care and Use of Animals and were approved by the Institutional Animal Care and Use Committee of Columbia University. Weakly electric mormyrid fish (7-12 cm in length) of the species *Gnathonemus petersii* were used for recordings. Fish were housed in 60 gallon tanks in groups of 5-20. Water conductivity was maintained between 65-100 microsiemens both in the fish’s home tanks and during experiments. For surgery to expose the brain for recording, fish were anesthetized (MS:222, 1:25,000) and held against a foam pad. Skin on the dorsal surface of the head was removed and a long-lasting local anesthetic (0.75% Bupivacaine) was applied to the wound margins. A plastic rod was attached to the skull with Metabond (Parkell) to secure the head and a craniotomy was performed over the C1 region of the cerebellum. Gallamine triethiodide (Flaxedil) was given at the end of the surgery (∼20 μg/cm of body length) to immobilize the fish and fresh aerated water was passed over the fish’s gills for respiration. The rate of the electric organ discharge motor command was monitored continuously by electrodes positioned near the electric organ in the tail. Electrosensory stimuli were delivered (0.2 ms duration square pulses) between an electrode in the stomach and another positioned near the tail. Probes were inserted vertically into the cerebellum along the midline and lowered slowly (∼10 μm/s) to a final depth of ∼3.5 mm. The data were batch sorted with Kilosort 2.0 using a 96 site template, and units were selected with an automated quality metric within Kilosort 2.0 like recordings from other species.

### Optotagging

#### Optotagging experiments

For optotagging experiments, four different transgenic mouse lines were used: 1.) Sst-IRES-Cre;Ai32, 2) Vip-IRES-Cre;Ai32, 3) Pvalb-IRES-Cre;Ai32, and 4) Sim1-Cre;Ai32. Ai32 is a reporter line that expresses channelrhodopsin in Cre+ cells. In each experiment we made recordings in parallel from 2-3 NP Ultra probes mounted on separate NewScale micromanipulators. Electrophysiological signals were acquired with OpenEphys software as described previously (Siegle et al., 2021). During the experiment, we recorded sequentially at 3-4 depths within the cortex. Probes were first inserted superficially in the cortex (∼200 μm deep) and allowed to settle for 10 minutes. We then ran the optogenetic protocol (described below). Next, probes were inserted deeper into the cortex (200-300 μm), allowed an additional 10 minutes to settle, and the optogenetic protocol was repeated. This process was repeated for up to 4 probe insertion depths. Recordings were made on a single day or on consecutive days from the same mouse.

We used a 470 nm laser to stimulate and optotag neurons expressing Cre and channelrhodopsin in each transgenic mouse line and experiment. For interneuron photostimulation, lasers were coupled to a 200 μm optic fiber. For Sim1-Cre;Ai32 L5 pyramidal neuron photostimulation, the laser path was guided by a pair of galvo-coupled mirrors to focus light to a precise point on the brain surface to minimize recurrent excitation. We used two different photostimulation patterns: a 10 ms square-wave pulse and a 1 s raised cosine ramp. On each trial, light was delivered at one of three light intensities levels (0.2, 4.1, and 10.0 mW/mm^2^). These six photostimulation inputs were randomly interleaved, and each was repeated for 100 trials. The average inter-trial interval was 2 s. In total, the photostimulation protocol lasted ∼20 minutes and recordings were conducted over a total of ∼30 minutes.

#### Identification of opto-tagged units

First-pass identification of opto-tagged units used an algorithmic approach to find units of similarly shaped peri-stimulus time histograms (PSTH) during a 1 second raised cosine ramp stimulation at the highest light intensity (see above). This method did not disambiguate between units that were directly stimulated and those that showed an indirect increase in firing rate. Identified units were used as a baseline comparator for further classification.

Unsupervised identification of opto-tagged units in each transgenic mouse line was performed similarly to (Jia et al., 2019). First, the PSTHs for all stimulation patterns within a given photostimulation intensity were averaged across all trials, then the average PSTH for each pattern and intensity was concatenated, forming a neurons × response-vector matrix. This matrix was then normalized and PCA was applied to reduce the dimensionality of the dataset. All units were then visualized in a low-dimensional space via Uniform Manifold Approximation and Projection (UMAP) using all timeseries PCs. Doing so allowed us to determine whether units with a similar PSTH structure aggregated to the same regions (**Supp. Fig. S21**).

Clustering of units was performed using Density-based Spatial Clustering of Applications with Noise (DBSCAN). We chose this method of clustering units as it does not require an *a priori* knowledge of the number of clusters expected and is robust against clusters of varying shapes and sizes, like those produced by UMAP. Hyperparameters for DBSCAN clustering were specific to experiments performed within each transgenic line and were obtained using K-Nearest Neighbors (KNN) to estimate the distance of each data point to the next closest point, then expanding this distance to include the 10th NN. The inflection point in the distribution of these distance values was used as the cluster radius (*ɛ*) for DBSCAN. The resulting clusters of units were then compared against those found in our first-pass sorting approach and validated by ensuring a short latency response during a 10 ms square wave pulse photostimulation.

### Supervised waveform classification of inhibitory neurons

#### Linear Discriminant Analysis

Linear Discriminant Analysis (LDA, **Fig. 7G**) was performed via its implementation in SciKit-Learn (Python version 3.9.12). Briefly, pairwise comparisons between cell classes proceeded first by randomly subsampling each dataset, without replacement, to the lowest number of units among the cell classes to be compared (PV, n = 238). Only prePTR and PTR single-channel 1D features were used in this analysis. Bootstrapping of the LDA model to subsamples of each cell class over 100 iterations with 100 random initial states and each iteration cross-validated five-fold. Performance of the model was quantified as how accurately the model classified each unit, given a label matrix. Accuracy scores were averaged across all iterations for each comparison to give a mean classification accuracy score and compared against chance levels of classifying a unit correctly (two classes per comparison, chance = 0.5).

#### Random Forest Classification

Validated opto-tagged units and untagged units formed the basis of a label matrix that was then used to classify neuronal types using random forest classification. Because significant differences could be found between RS, FS_L_, and FS_S_ groups (**Fig. 7G**, **8B**, **Supp. Fig. S26**), these units were included as distinct categories during classification.

Random forest classification was performed via its implementation in SciKit-Learn (Python version 3.9.12) using three different feature sets computed independently on mean electrophysiological waveforms gathered from NP Ultra probes and interpolated, “NP 1.0-like” unit-matched data. These feature sets were 1) 1-dimensional scalar values computed from the peak amplitude channel (**Fig. 8A**, **Table 4**) 2) the peak amplitude waveform and 3) All 1D scalar features and spatial footprint.

The process of random forest classification proceeded as the following: NP Ultra data from the entire population of recorded units was spatially resampled to a NP 1.0-like geometry (**Fig. 2E**). Units for which the peak channel amplitude dropped below 50 μV were then removed from both datasets for the purposes of this analysis. Data (NP Ultra and NP 1.0-like) from each cell class category was then randomly subsampled, without replacement, to the lowest number of units among optotagged cell classes (SST, 116 units) and pooled. This process was repeated through 100 iterations using 100 initial states. Principal components analysis was then performed on the pooled data for each probe type prior to classification and classifier hyperparameters were then optimized via grid search using five-fold cross validation. Classification proceeded via five-fold cross-validation where the classifier was trained on 80% of the input data and performance was evaluated using held-out test data (20%). Classification performance was evaluated as the prediction accuracy of the classifier on left-out data over each bootstrapped iteration. Confusion matrices were computed as the comparison of predicted classes to true classes for each subsampled dataset under 100 random initial states.

## Acknowledgements

We thank NIH program officers Ned Talley, Sahana Kukke, and Michele Pucak for their support, and we thank Tim Gardner and Cindy Chestek for their advice and guidance.

## Author contributions

ZY collected brain-wide acute mouse recordings and pharmacology experiments. AMS and SM collected data with opto-tagging. ZY and JRS collected data with imposed probe motion and visual stimuli. JC performed noise and gain measurements. AMS performed light sensitivity measurements. SC collected acute mouse recordings. JHS wrote software for data acquisition. TN and WB collected non-human primate recordings. FP and NBS collected recordings from electric fish. SW collected recordings from bearded dragon lizards. LFR and GM collected and analyzed data on dendritic back-propagation. JB, CW, CH, and LP developed the spike sorting methods. ZY, AMS, JC, JB, JRS, and NAS analyzed data. DB developed the data exploration website. CML, BR, and BD performed engineering and fabrication of NP Ultra probes. TVN wrote the computational model and NAS analyzed the output. ZY, AMS, JC, JB, JRS, LP, SRO, and NAS wrote the original draft manuscript. All authors reviewed and edited the manuscript. SRO, TDH, and NAS obtained primary funding. XJ, MC, MH, LFR, GTE, GL, NBS, WB, AP, CML, BD, LP, JHS, CK, SRO, TDH, and NAS supervised aspects of the work.

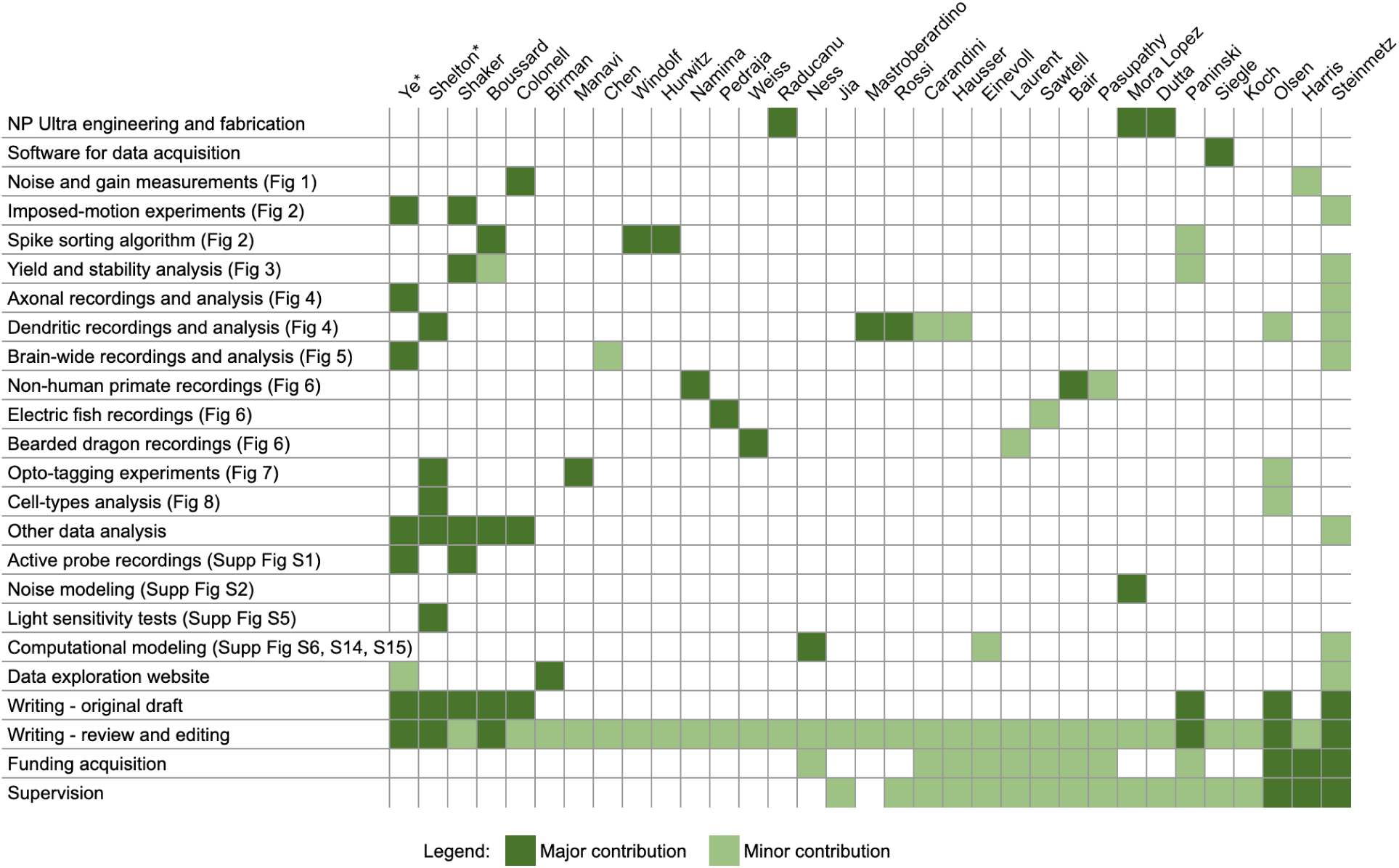

## Data and materials availability

Spatiotemporal waveforms of 4,666 single units recorded across cortex, hippocampus, thalamus, striatum, and midbrain areas in awake mice, along with spike times are available at: https://doi.org/10.6084/m9.figshare.19493588; and the waveform movies are browse-able at https://npultra.steinmetzlab.net/.

NP Ultra dataset for imposed motion with visual responses pre- and post-motion in the visual cortex is available at: https://dandiarchive.org/dandiset/000957.

DARTsort code is available at: https://github.com/cwindolf/dartsort.

## Supplemental Figures

**Supplemental Figure S1.**
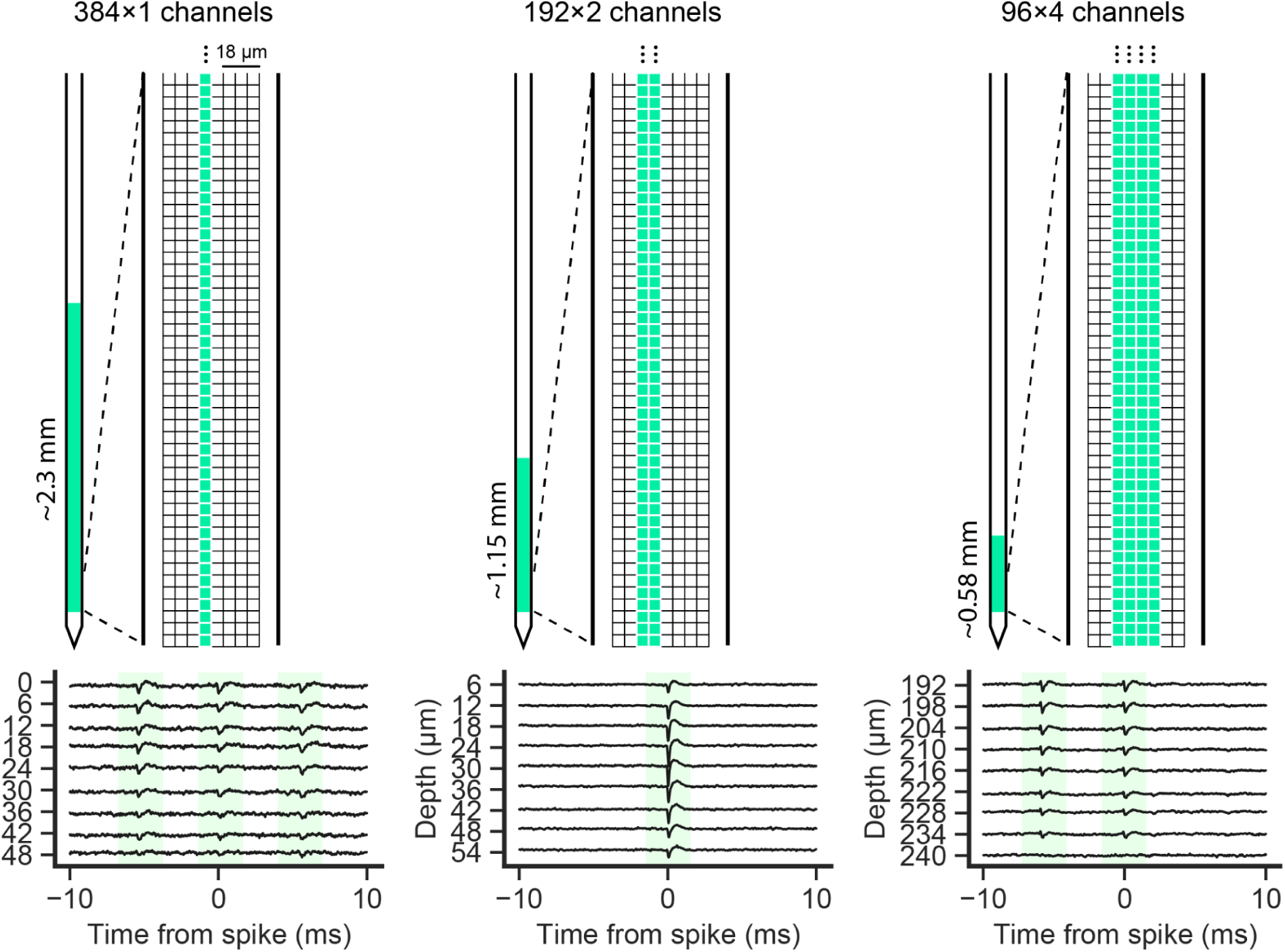
Schematics of site layout and raw data traces from three different configurations of the NP Ultra probe. (Top, left to right) 384 × 1 configuration of sites, 192 × 2 sites, and 96 × 4 sites. For both the 384 × 1 and 192 × 2 configurations, not all recorded sites are in the same column, instead there are lateral shifts in the columns of the selected sites (not shown). (Bottom) Raw data traces 20 ms about a detected spike excerpted from ∼5 minute-long recordings in mouse cortex. Detected spikes highlighted in green.

**Supplemental Figure S2.**
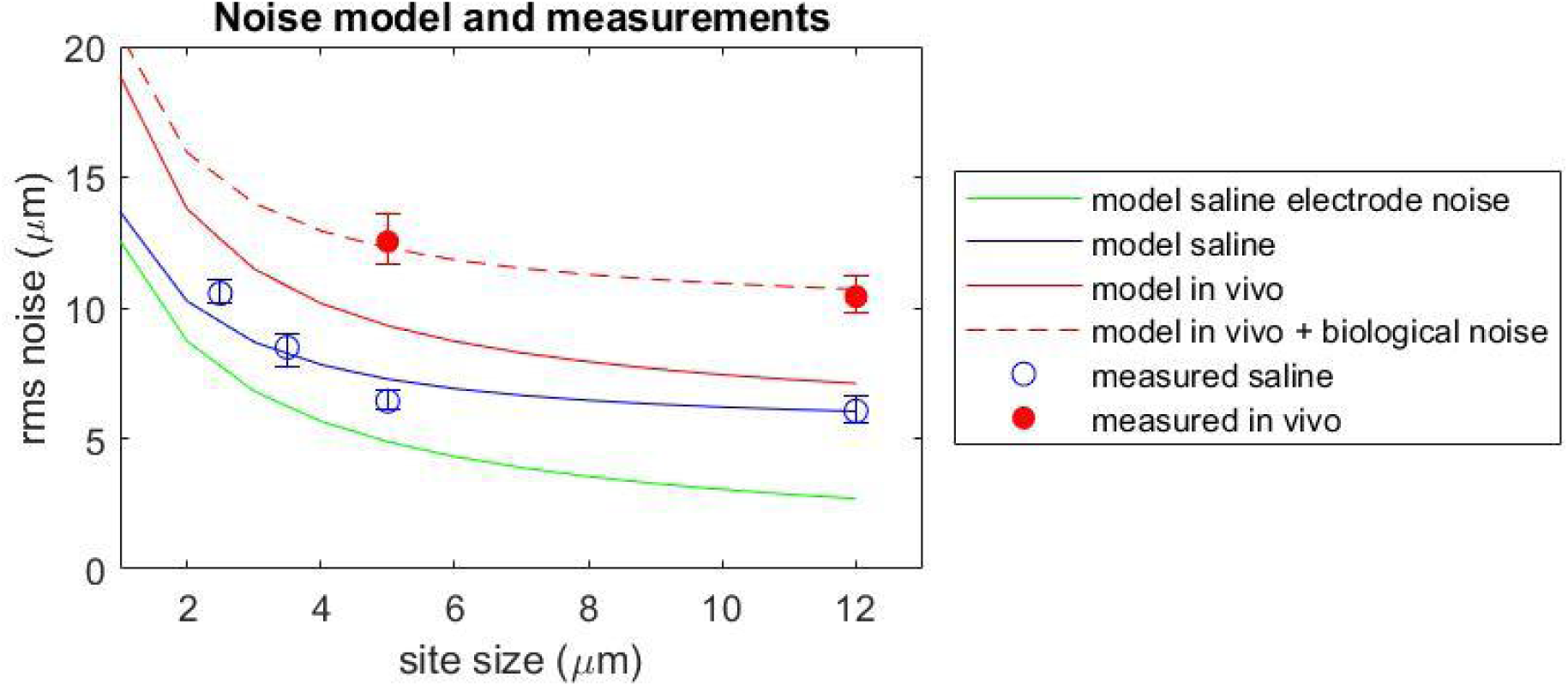
Modeled and measured noise as a function of electrode site size. The solid curves show the predicted noise as a function of site size. The model uses estimated impedances given the characteristics of the TiN recording site material (see Methods). The green curve shows the predicted thermal noise at the TiN electrode in saline; the blue curve shows the predicted total noise in saline, which includes the thermal noise and the noise in the Neuropixels electronics. The solid red curve shows the predicted total noise in vivo, accounting for the higher impedance (taken to be 300 Ohm-cm), but not including biological noise; see Lopez, et al., 2012 for details). Data points show experimental measurements of noise in saline and in vivo; error bars show the 25th and 75th percentile of all sites measured. The measured in vivo noise is expected to be higher than the model because it includes biological noise, i.e. small signals from distant units. The dashed red curve shows the predicted in vivo total plus biological noise, fit to the two in vivo data points. The inferred biological noise is 8 µV.

**Supplemental Figure S3.**
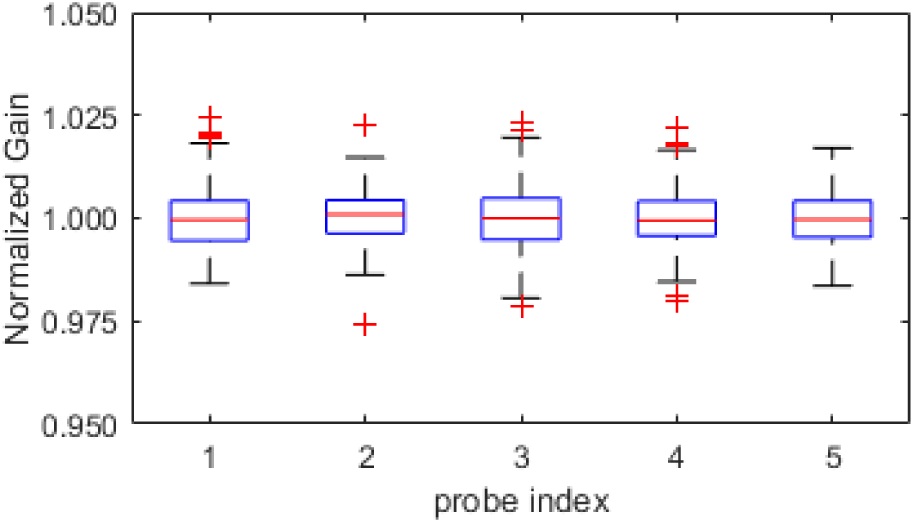
Channel-to-channel uniformity in gain in NP Ultra. Peak to peak voltage was measured at each site when applying a 3 kHz, 2mV sine wave to a saline bath for 5 probes.Standard box and whisker plots show distribution of normalized gain measured across the 383 recording sites on the probe (reference site is excluded). Box limits = 25th-75th percentile; whiskers = max/min within 1.5X the 25-75th percentile; points = outliers outside the whisker range. Note that these data show that error intra-site voltage measurements is < 5% for all sites.

**Supplemental Figure S4.**
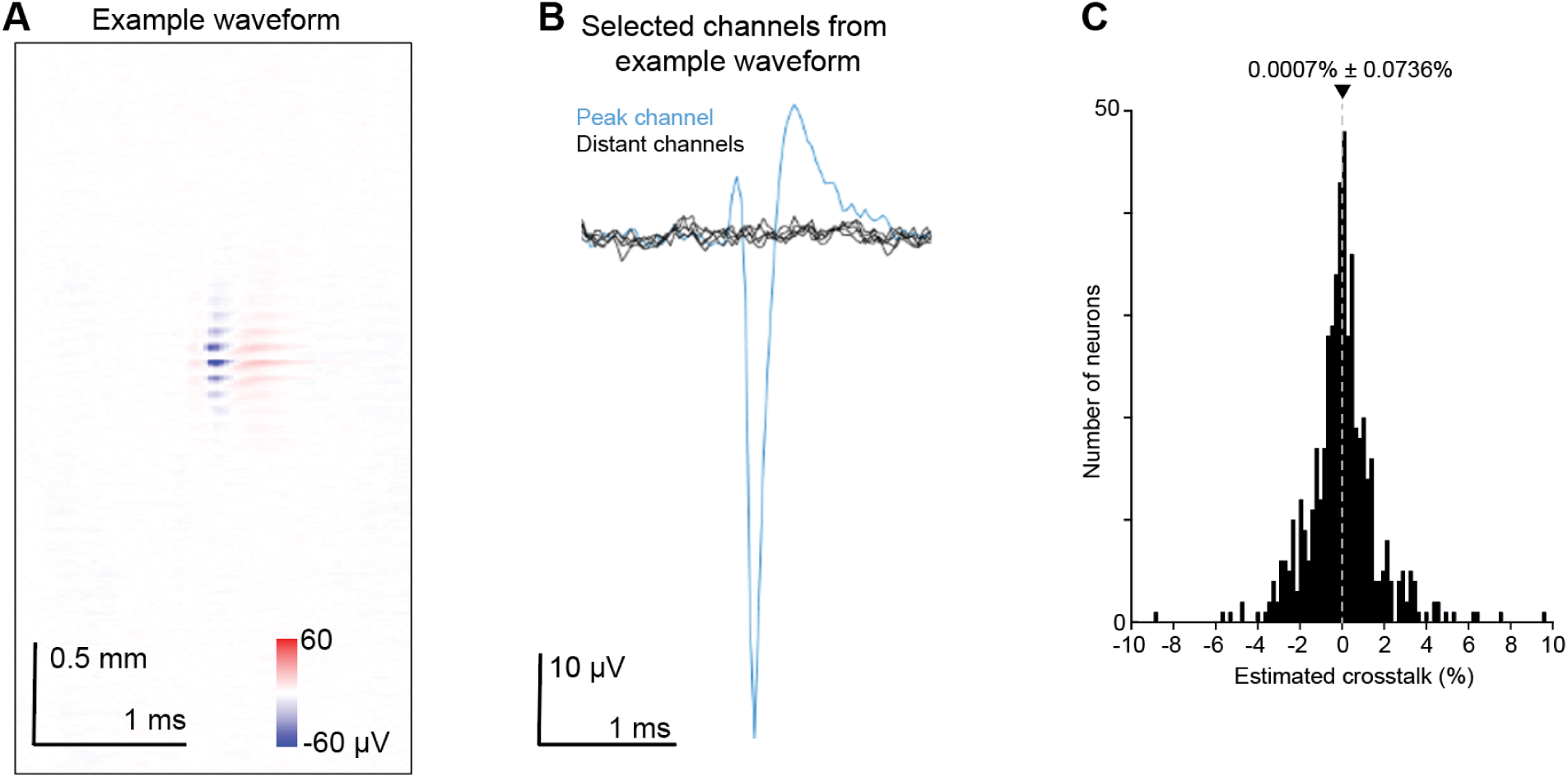
Crosstalk analysis of NP Ultra recordings. **A**, An example neuron’s average waveform across channels (y-axis) and time (x-axis). **B**, The waveform on the peak channel and on distant channels. For this example neuron, the amplitude at the distant channels was 1.25% of the peak amplitude. **C**, Summary histogram of estimated crosstalk across the n=519 neurons included in the analysis. The estimated crosstalk was on average 0.0007% ± 0.0736% (mean ± s.e.m.).

**Supplemental Figure S5.**
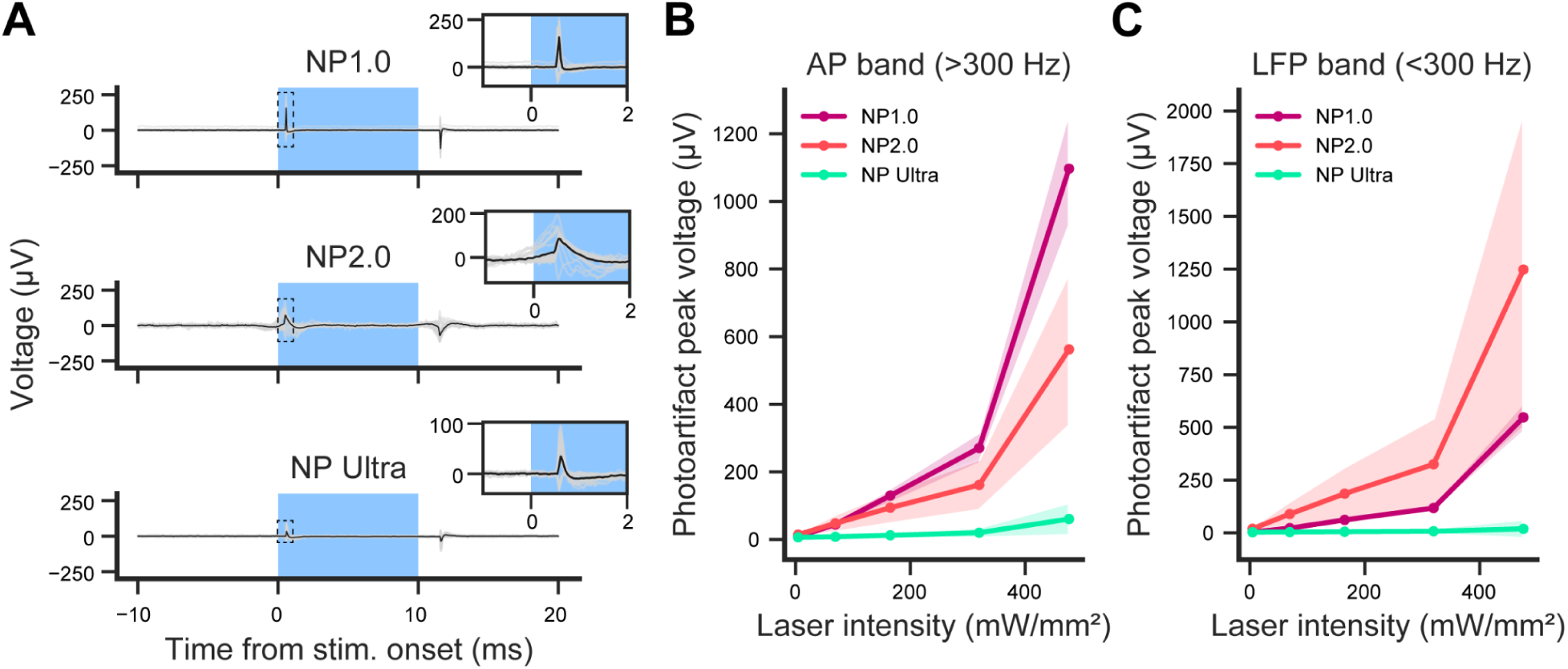
Light sensitivity tests. **A**, Example experiment showing the light-induced artifact measured on NP 1.0, NP 2.0, an NP Ultra probes. Gray traces are all channels, black line is the average across channels. Insets focus on the initial positive peak of the photoartifact occurring shortly after light onset. **B**, Peak photoartifacts for each probe type in **A** averaged across trials in the AP band (>300 Hz). **C**, Same as in **B** for LFP band data. The physical explanation for smaller artifacts in NP Ultra probes is unclear, but it is important to note that the NP Ultra probes were the passive, non-switchable version whereas 1.0 and 2.0 probes included CMOS switching circuitry.

**Supplemental Figure S6.**
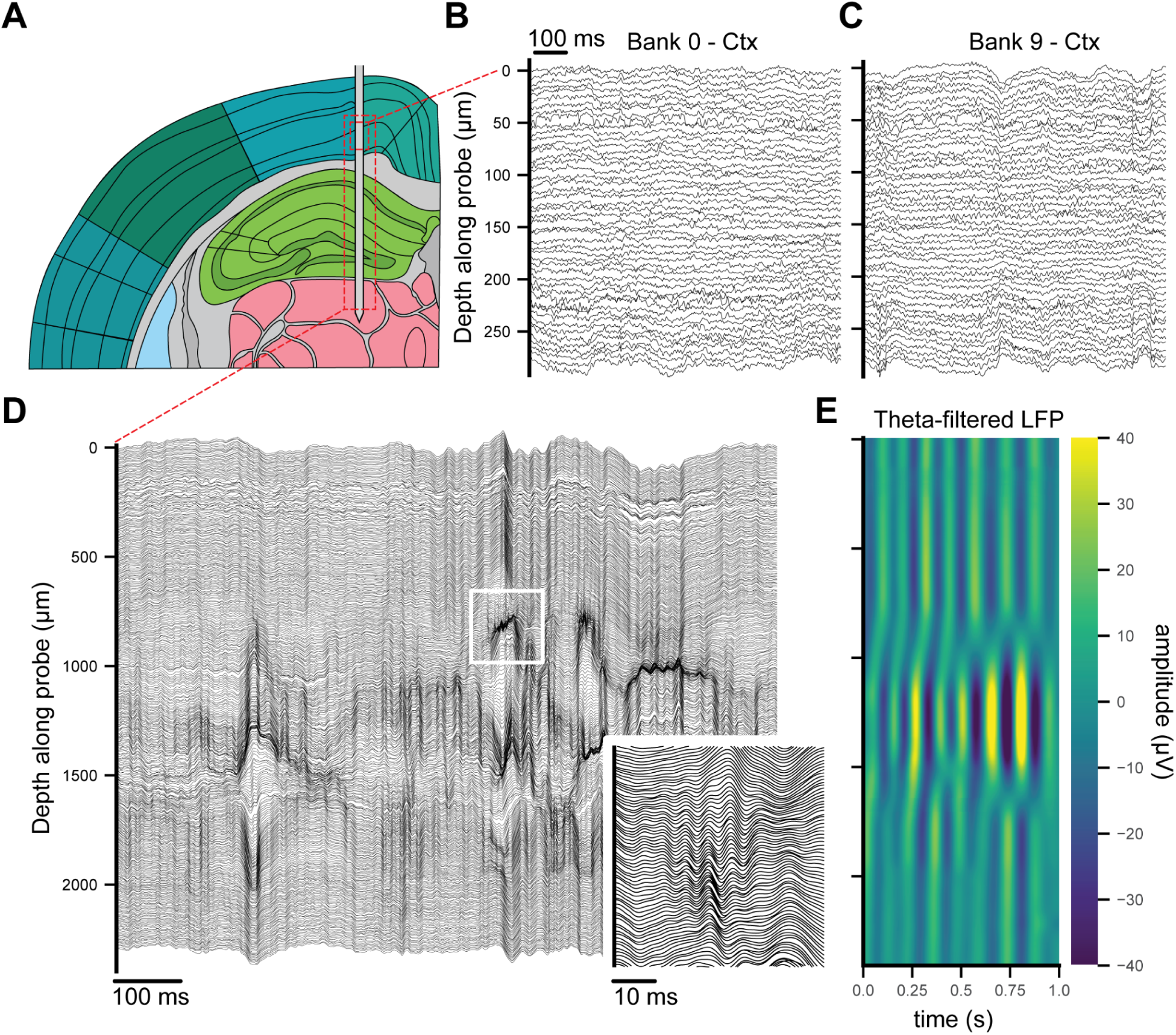
LFP signal in cortex, hippocampus, and thalamus using standard and linear NP Ultra probe configurations. **A**, Schematic of insertion location. A switchable NP Ultra probe was inserted sequentially through cortex, hippocampus, and thalamus and recordings were made in the standard 48 × 8 and linear 384 × 1 probe configurations. **B**, Initial LFP recording from cortex made using the 48 × 8 probe configuration. LFP data was filtered using a 3rd order Butterworth filter between 0.5 Hz and 300 Hz. **C**, Final LFP recording from cortex using a bank of contacts (48 × 8 configuration) higher up on the probe after full insertion to thalamus. **D**, LFP recording from thalamus to cortex in the linear probe configuration. Inset shows sharp wave ripple localized to a subset of channels, filtered between 20 Hz and 250 Hz. **E**, Heatmap showing the theta-band LFP signal filtered between 4 Hz and 15 Hz across the span of the probe in the linear configuration as in D.

**Supplemental Figure S7.**
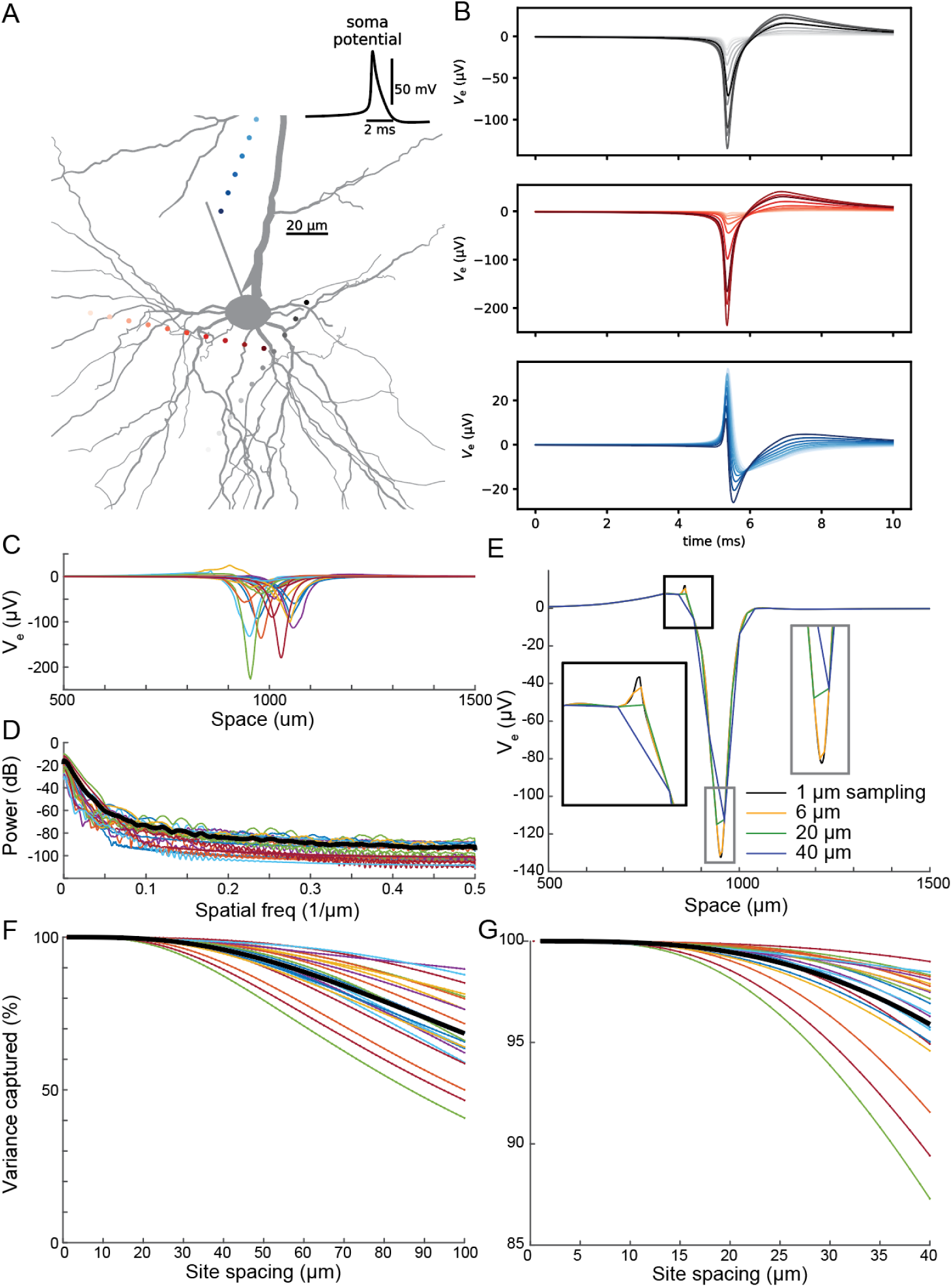
Computational modeling shows that most spatial features of extracellular action potentials are captured by measurements with the site spacing of NP Ultra and NP 2.0. **A**, A detailed biophysical simulation of the extracellular action potential (EAP) of a multi-compartmental single neuron (gray) was sampled along random linear trajectories through the 3D space near the neuron. The somatic intracellular waveform is shown in the inset. Colored dots indicate example points where the EAP was computed. **B**, Waveforms (voltage, V_e_, over time) from these example EAP sample points, with colors matching the dots in A. **C**, The spatial profile at the peak time point was extracted from each random trajectory, at 1 µm resolution. Each colored trace represents a different trajectory from near the same neuron. The exact coordinates of each trajectory were chosen randomly, so the peak of each profile (representing the point of closest approach to the neuron) has a different position on the x-axis. **D**, The spatial power spectrum of each trajectory (colors) and mean power spectrum (thick black line) reveal a gradual fall-off in power to higher frequencies. **E**, To visualize and quantify the impact of sampling at coarser spatial resolution, the original spatial voltage profile at 1 µm resolution was downsampled and reconstructed by linear interpolation at 6, 20, and 40 µm sampling resolutions (roughly corresponding to the sampling resolution of NP Ultra, NP 2.0, and NP 1.0 along the axis of the probe), here shown for a single example spatial profile. The peak amplitude is underestimated and some small features are missed with coarser sampling, as highlighted in the insets. **F**, Success of reconstruction quantified as the percent variance of the original spatial profile that is maintained by the downsample, for each trajectory sampled in panel C. **G**, Zoom in over the range 1 to 40 µm sampling resolutions. The spatial profiles are reconstructed with high fidelity (∼99% on average) with sampling of NP 2.0 but begin to lose significant information by the 40 µm spacing of NP 1.0, especially along certain trajectories. This failure to capture the spatial profile well presumably corresponds to the failure to track neurons sampled with 1.0-like patterns across probe motion as reliably as with NP Ultra and NP 2.0 (**Fig. 3**).

**Supplemental Figure S8.**
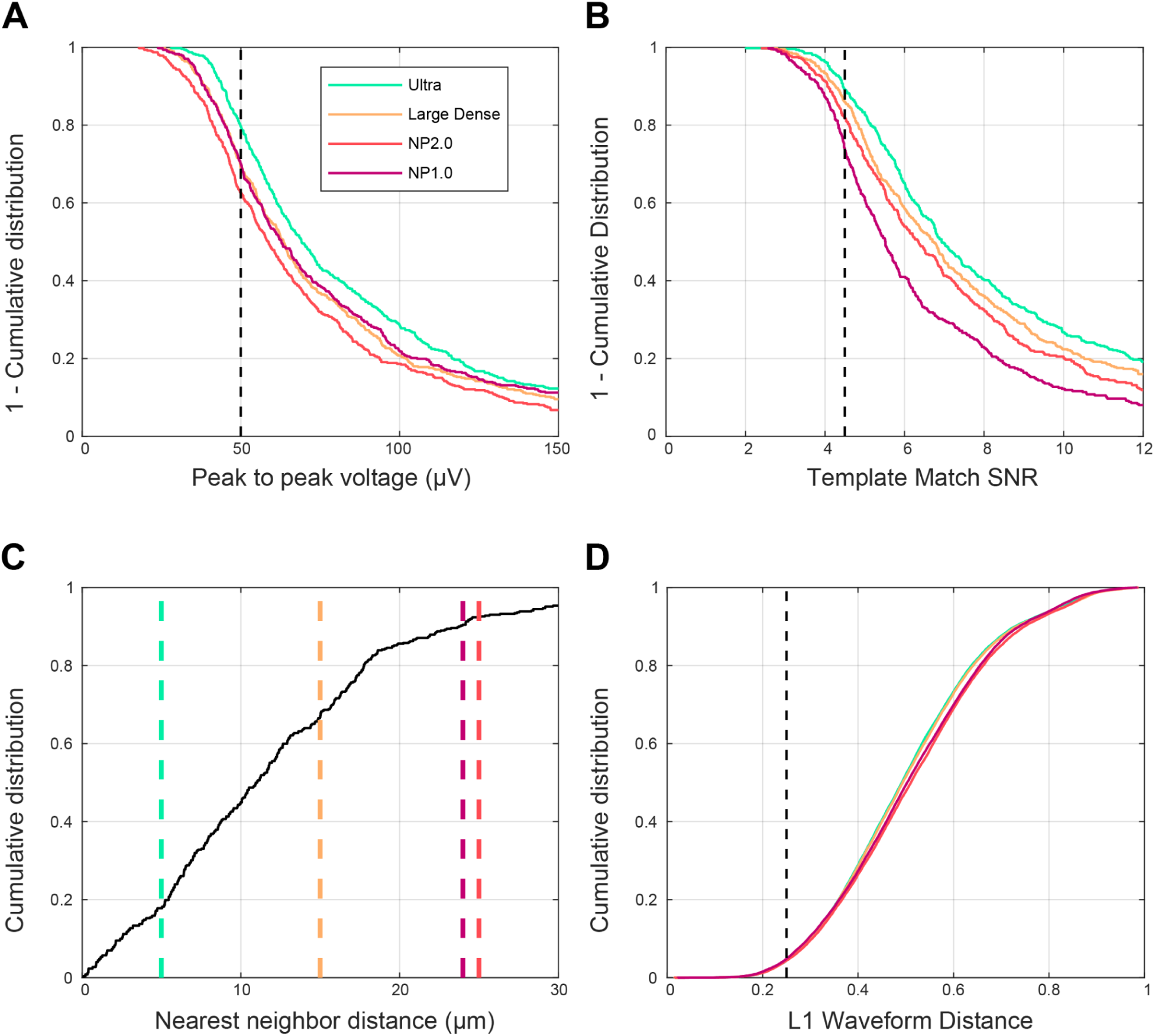
Detailed spike sorting analysis. The goal of this ‘sortability’ analysis is to parse the effects of increased site density on the ability to detect and sort spikes. To optimize for the largest spatial range while detecting the largest number of sortable units, we want to understand what aspects of the data make spikes more detectable – that is, more readily found by thresholding or template matching – and then correctly assigned to a specific unit. The ideal sampling is a function of the signal: Very small footprint units require dense sampling to detect them, while units with larger footprints, spaced well apart from each other, will be adequately detected at lower site density. The distribution of unit characteristics is expected to vary across brain regions. With curated NP Ultra data, we can examine the most complete set of units available and estimate the impact of lower spatial sampling. Measurements of the features of extracellular electrophysiology signals that contribute to sorting are summarized in this and the following figure. See **Fig. 2E** for site patterns. Panels **A-D** show distributions of these features for units measured in 11 separate NP Ultra recordings in mouse ALM. **A,** Cumulative distributions of peak to peak voltage, which is the most important feature for detecting spikes via thresholding. The shift to lower Vpp for sparser patterns is due to sampling further from the peak signal, so that a larger fraction of units will not be detected at a given threshold. Although NP 1.0 is slightly sparser than NP2.0, it covers a larger area on the probe, so its Vpp distribution is slightly higher than NP 2.0. The dashed vertical line at 50 μV indicates the threshold used in **Supp. Fig. S9**. **B,** Cumulative distributions of Template Match SNR showing significant shifts to lower SNR at sparser patterns; as discussed in the Methods section, TM-SNR is an estimate of the SNR for spike detection via thresholding of data filtered by a template. The dashed vertical line at 4.5 indicates the threshold used in **Supp. Fig. S9**. **C**, Cumulative distribution of physical nearest neighbor distances, including ‘unimodal’ and ‘overlapped’ units (see Methods for Analysis of sorting quality ), to build as complete a picture of the activity as possible. The effect of the lower spatial resolution of sparser patterns is that more units have overlapped distributions of spike locations. Pairs of units separated by less than spatial resolution of the probe may still be sorted by other features, but can not be fully separated by spike localization alone. **D**, Distribution of all pairwise waveform shape distances for all unimodal units in this set of recordings; see Methods for a description of the waveform distance metric. Similar distributions across site density patterns, indicates that the spatial variation of waveforms for most units is smooth enough to be adequately detected at lower site density. See **Supp. Fig. S9** for quantification of units passing a threshold of 0.25 (vertical line; corresponds to 25% of detected signal differs between two templates). Note that this assessment takes no account of noise or waveform variability, and thus underestimates the difficulty of real sorting.

**Supplemental Figure S9.**
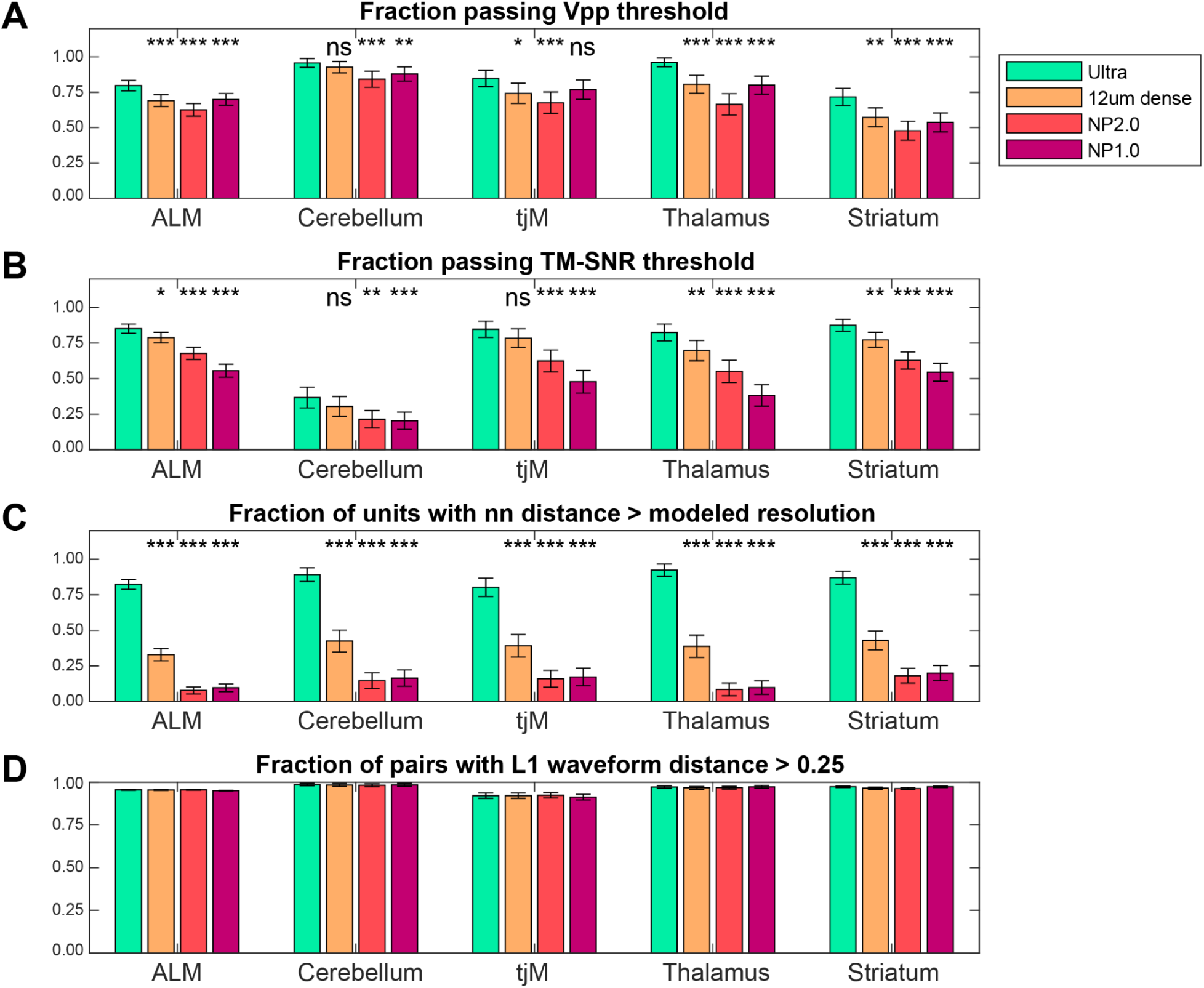
The impact of varying site patterns on key features for sorting, across a variety of brain regions. Regions with a larger fraction of small footprint units will have more units that fall below reasonable detection thresholds at lower site density. Regions with closely spaced units will have more pairs that fall below the spatial resolution of low site density probes, which can lead to sorting errors. **A,** Fraction of units with peak-to-peak voltage (Vpp) > 50 μV. **B,** Fraction of units passing TM-SNR > 4.5. **C,** Fraction of units with separation greater than the ideal spatial resolution of the site pattern (see thresholds in figure **Supp. Fig. S8**. **D,** Fraction of pairwise distances greater than 0.25. For all, errors bar shows 2*standard error of the proportion above threshold assuming an underlying binomial distribution. For each region, asterisks show significance of difference from Ultra pattern, calculated using 2-proportion z-test. Differences in fraction of pairs with L1 > 0.25 (panel D) are not significant according to this test. ALM: Anterolateral motor cortex; tjM: tongue-jaw motor cortex.

**Supplemental Figure S10.**
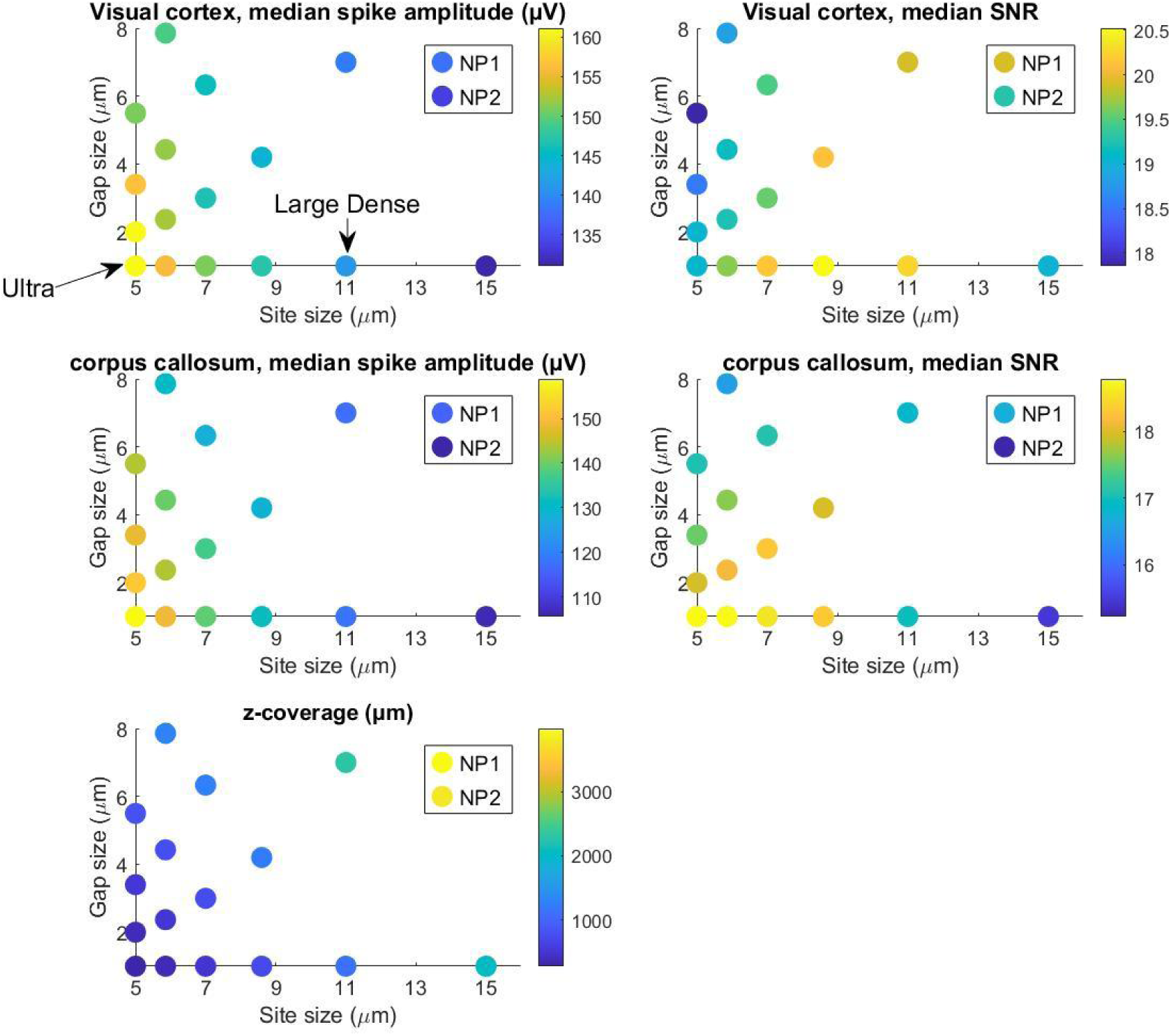
Predicting optimal site layout. Using interpolations of the measured average spike waveforms from the recordings discussed in Figure 5, we can predict the observed signal amplitude (peak to peak voltage) and signal to noise ratio (calculated here as amplitude/noise, with noise taken to be the predicted value from the model shown in S2) for a range of site patterns. In these plots, each point is colored by the median amplitude (right column) or median SNR (left column) for all units in a region, detected with a square site pattern with a given site size (x-axis) and edge-to-edge gap size (y-axis). All patterns cover the same 48 × 288 *μ*m area, matching the area of the Ultra probe. With small site sizes, the amplitude is higher, because the electrode averages over a small area and can sample the peak of the electric field. However, thermal noise increases with reduced site size, so SNR is reduced. The optimal site layout is a function of the average spatial footprint of units. In regions with many large footprint units, e.g. the visual cortex (top row) the peak SNR occurs with larger sites with small gaps (see points along the site size axis). In contrast, in corpus callosum (second row), units with small footprints dominate, and the best SNR is achieved with smaller site sizes. The predicted amplitude and SNR for NP 1.0 and NP 2.0 patterns are plotted as separate points. See the colorbar on each plot for the magnitude of these differences. The best detection is always achieved with the smallest gap size, but at the cost of reduced tissue coverage at a given site count. The map in the third row shows the z-coverage for the patterns tested, illustrating the trade off between density and coverage with a fixed channel count of 384.

**Supplemental Figure S11.**
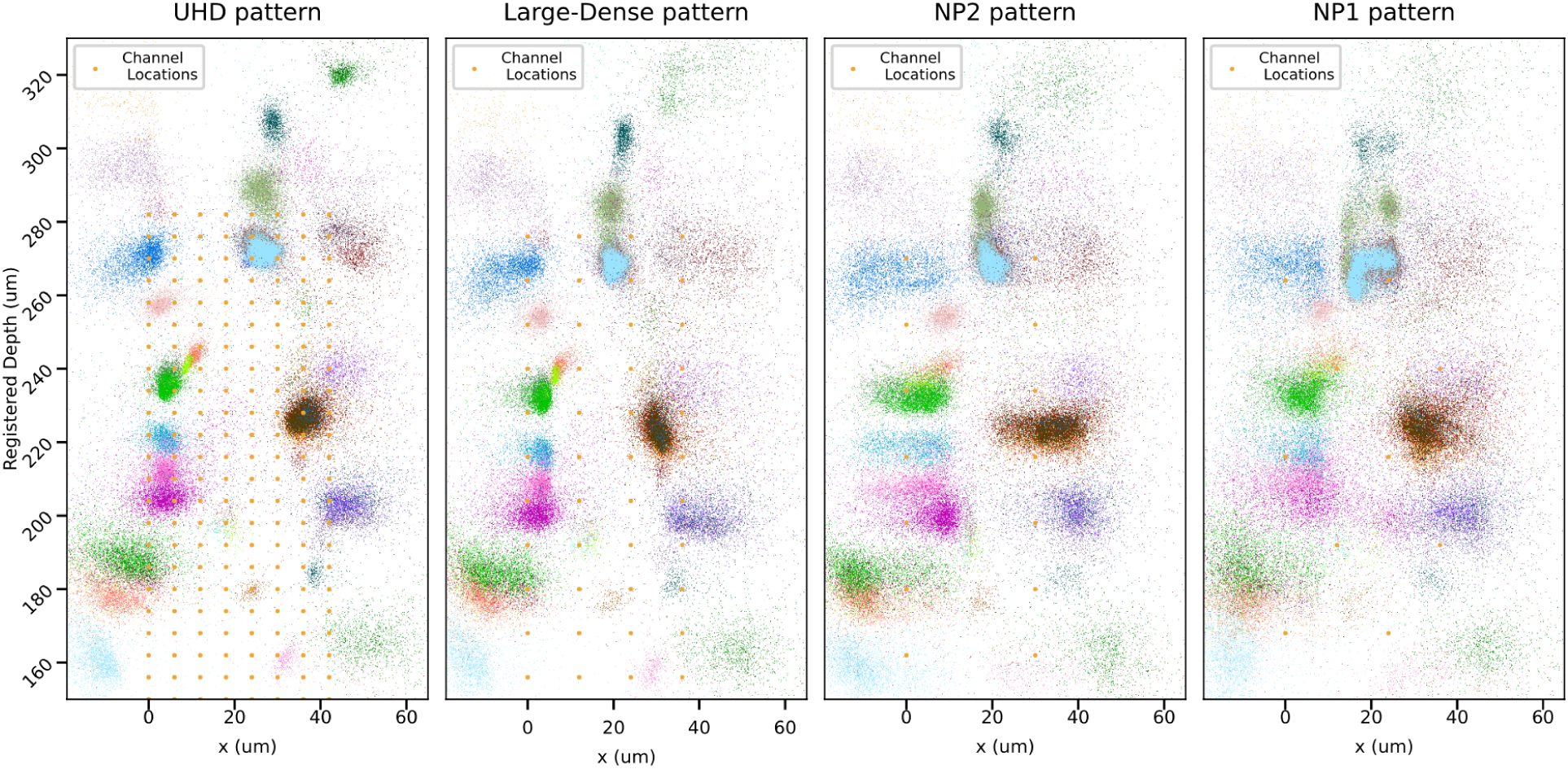
The impact of varying site patterns on spike localization. Spikes were detected and clustered according to their location and amplitude obtained on the original UHD recordings. They were then “relocalized” for each re-sampling pattern. The four panels show the spike locations above 150 μm, on the plane of the probe. We see that clusters are more spread out as channel density decreases. This effect is amplified when the source neuron is outside of the probe boundaries.

**Supplemental Figure S12.**
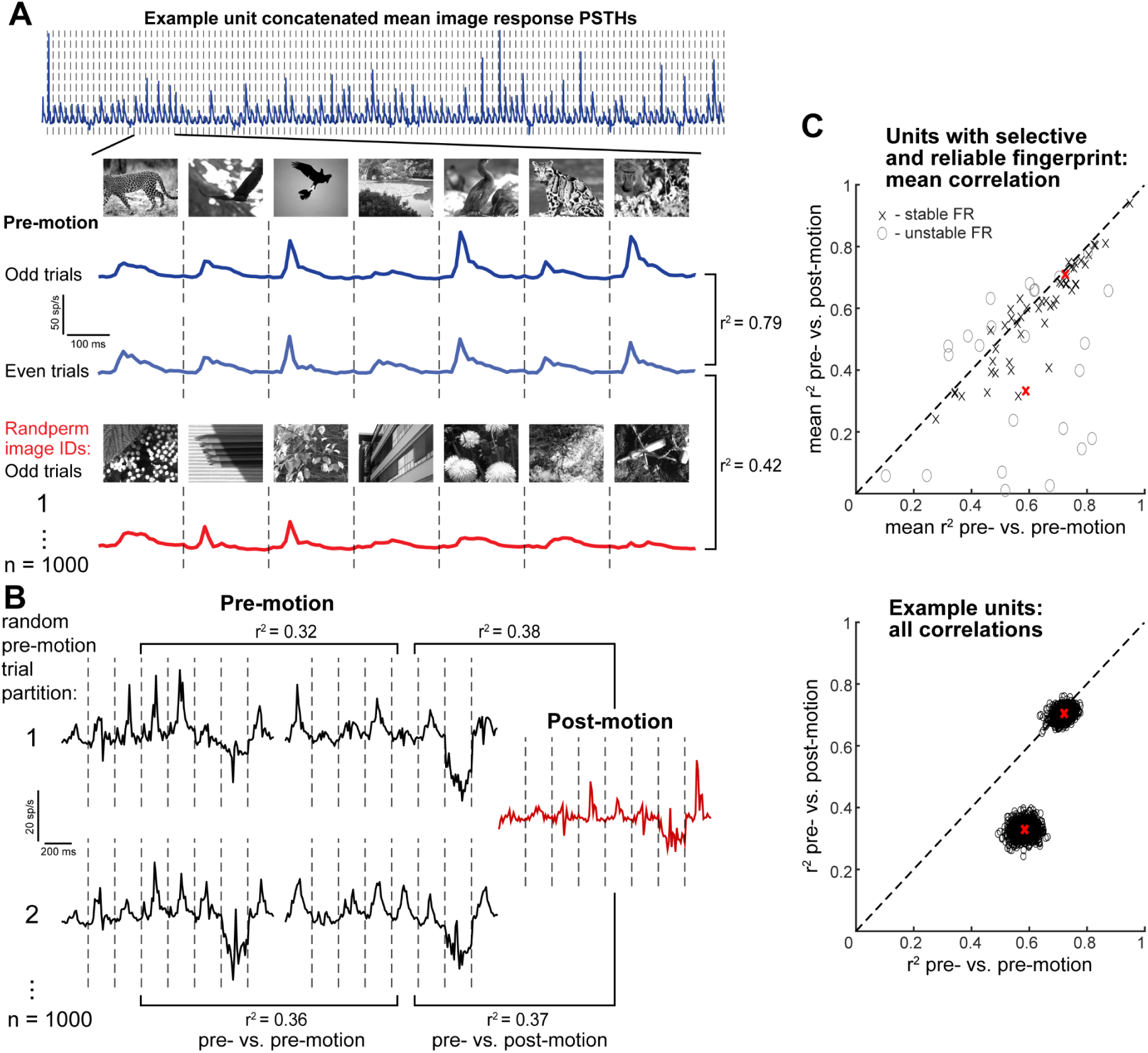
Methods for stability and yield analyses. **A**, Fingerprint calculation and inclusion criteria. An example neuron’s visual response “fingerprint” is shown in the top panel; it is calculated by concatenating the PSTHs (0 to 200 ms) aligned to the onset of 118 different natural images. Inset: Fingerprint of the same neuron for 6 images calculated from the odd-numbered trials prior to imposed motion and the even-numbered pre-motion trials. A null distribution is created by randomly permuting image identities and re-concatenating into the fingerprint vector 1000 times. This preserves the noise and nonspecific visual responses of a neuron while destroying the structure of any selective visual response pattern it may have. A neuron passes the inclusion criteria if the pre-odd vs. pre-even correlation is greater than all 1000 pre-even vs. shuffled correlations. **B**, Assessment of fingerprint tracking stability. Of the neurons passing the inclusion criteria for fingerprint reliability and selectivity, 1000 unique random partitions of the 30 pre-motion trials are created (2 sets of non-overlapping 15 trials for each partition) and fingerprints are constructed from each set. Fingerprints from each subset of pre-motion trials are correlated with each other, and one pre-motion fingerprint subset is correlated with the post-motion fingerprint. **C**, Top plot: mean pre- vs. pre- and pre- vs. post-motion correlations for all neurons that passed the inclusion criteria for fingerprint reliability and selectivity in an example NP Ultra session. Gray circles represent neurons with an unstable firing rate (>4x difference in mean PSTH firing rate pre- vs post-motion). Bottom plot, all 2000 correlations from each random partition are shown for 2 example neurons. Each dot represents the 2 correlation scores from the 1000 random partitions. Mean correlation scores for these two example neurons are marked by red X’s in bottom and top plots.

**Supplemental Figure S13.**
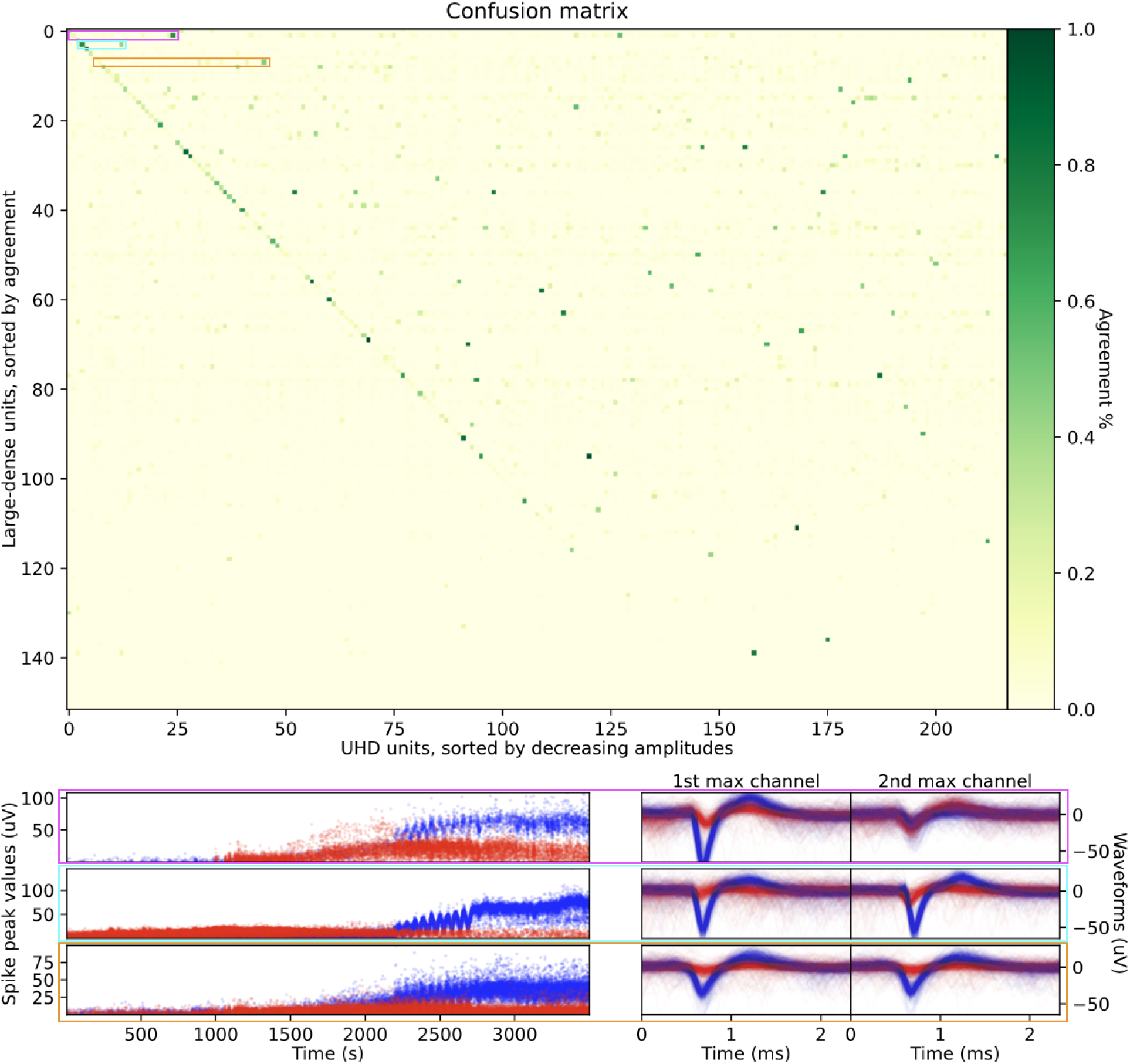
Improved yield for NP Ultra is not due to over-splitting units. To check that larger yield from NP Ultra probes is not due to oversplit units, thus artificially leading to a larger number of good units compared to resampled data sorts, we examined the confusion matrix between the Ultra and Large-dense sort (Top). The Ultra units (x-axis) are sorted by decreasing amplitudes, and Large-dense units (y-axis) by agreement percentage. Although most units have low agreement, partly because many more spikes are discovered in the Ultra data, we found some high amplitude units being potentially split i.e. multiple Ultra units having high agreement with a single Large-dense unit. We isolated, for the three highest-amplitude potential split units, the combination of units being merged in the Large-dense sort (pink, cyan and orange rectangles). For these three pairs, we plot their peak absolute value (Bottom left), and 500 randomly sampled waveforms on each unit’s max channel (Bottom right) in red and blue respectively. Waveforms are sampled after the imposed motion period ending around 3000s to avoid motion-induced variability. The lack of overlap between peak amplitudes and waveforms supports the notion that each of these pairs of units are indeed distinct.

**Supplemental Figure S14.**
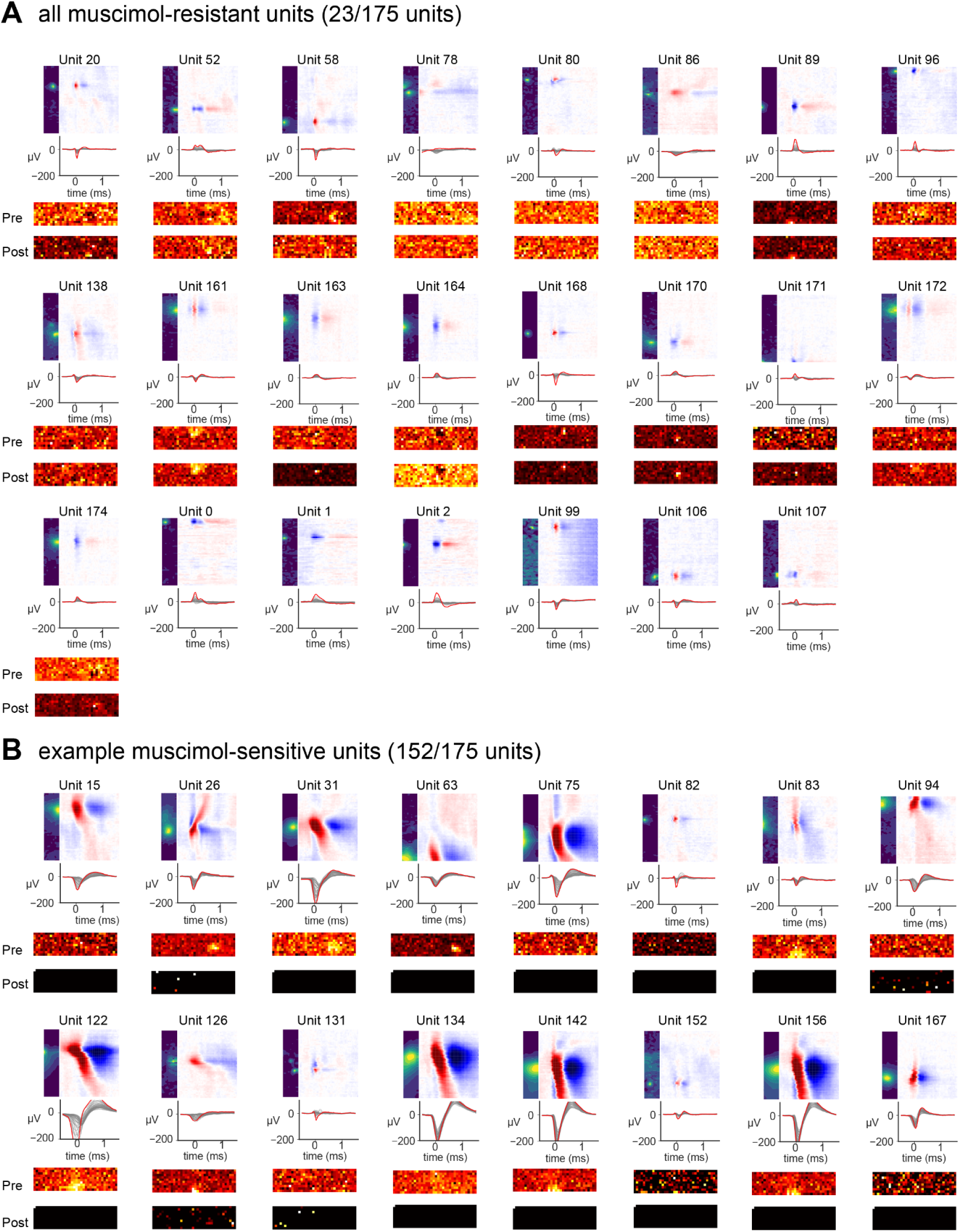
Muscimol-resistant and muscimol-sensitive waveforms and receptive field maps before and after muscimol application. **A**, waveform shapes and receptive field maps of all muscimol-resistant units (23/175 units). 17/23 units in VISp had receptive field maps, while 5/23 units were recorded in MOp without receptive field maps. Note that receptive field maps changed for some units but not others, and all muscimol-resistant units had small spatial footprint (spatial extent of channels with detectable signal >30 µV). Top: waveform shape plots with three views: probe width × height, height × time, and peak channel over time. Middle: receptive field maps before muscimol. Bottom: receptive field maps after muscimol. **B**, Example muscimol-sensitive units with receptive field maps in VISp. Note that all receptive field maps disappeared due to the silence of neural activity in muscimol, and muscimol-sensitive units had a mix of large and small spatial footprints.

**Supplemental Figure S15.**
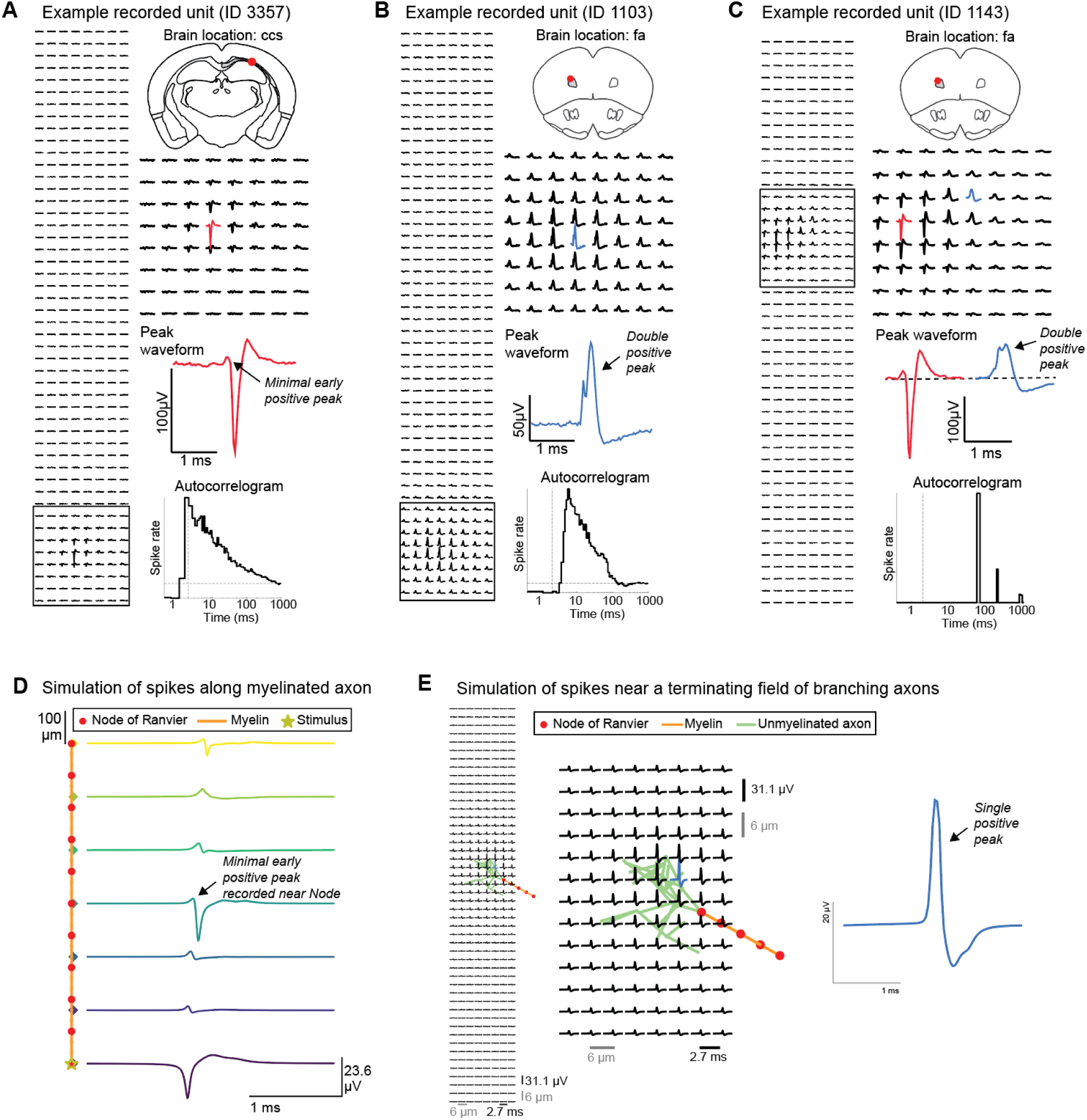
Examples of white matter putative axonal recordings, and biophysical simulations of axonal extracellular action potentials. **A**, An example unit recorded in the white matter which has a small-footprint, negative-going action potential. Left, Waveform on all 384 channels, showing a small footprint. Right, unit location (red dot on coronal slice; ccs: corpus callosum splenium), peak channel waveform exhibiting minimal early positive peak, and autocorrelogram (ACG) on log time axis (vertical dashed line at 2 ms). (https://npultra.steinmetzlab.net/?selected=3357 to browse this neuron and view waveform movie) **B**, An example unit recorded in the white matter (fa: corpus callosum anterior forceps) which has only positive-going action potentials on every channel. Waveforms exhibit a double positive peak. Format as in **A**. (https://npultra.steinmetzlab.net/?selected=1103) **C**, An example unit recorded in the white matter which has a region with negative-going action potential (similar to **A**) and a spatially-adjacent region with double-peaked positive-going action potential (similar to **B**). Format as in **A**. (https://npultra.steinmetzlab.net/?selected=1143) **D**, Simulation of extracellular action potential recorded at various positions along a myelinated axon. The physical arrangement of the simulation is shown at left, with simulated traces recorded at each of the colored diamonds shown to the right . Near a Node of Ranvier (red dots), the waveform is negative-going and narrow with minimal positive-going component (similar to the unit from data shown in panel **A**). See also simulations in Chapter 7 of (Halnes et al., 2024). The early positive peak, when it occurs, reflects return currents of the distant spike as it approaches the recording site along the axon, but can have minimal amplitude depending on the parameters of the simulation. This simulation uses 1 µm diameter axons, 100 µm inter-Nodal spacing, and 1000 µm total axonal length. **E**, A branching axon termination field also produces positive-going spikes, as demonstrated in (McColgan et al., 2017), but typically without a double-peaked shape. The double-positive peak therefore remains mysterious. Here, recordings are simulated from a grid of sites matching the NP Ultra site layout, with the physical layout of the simulated axon shown behind the grid of waveforms.

**Supplemental Figure S16.**
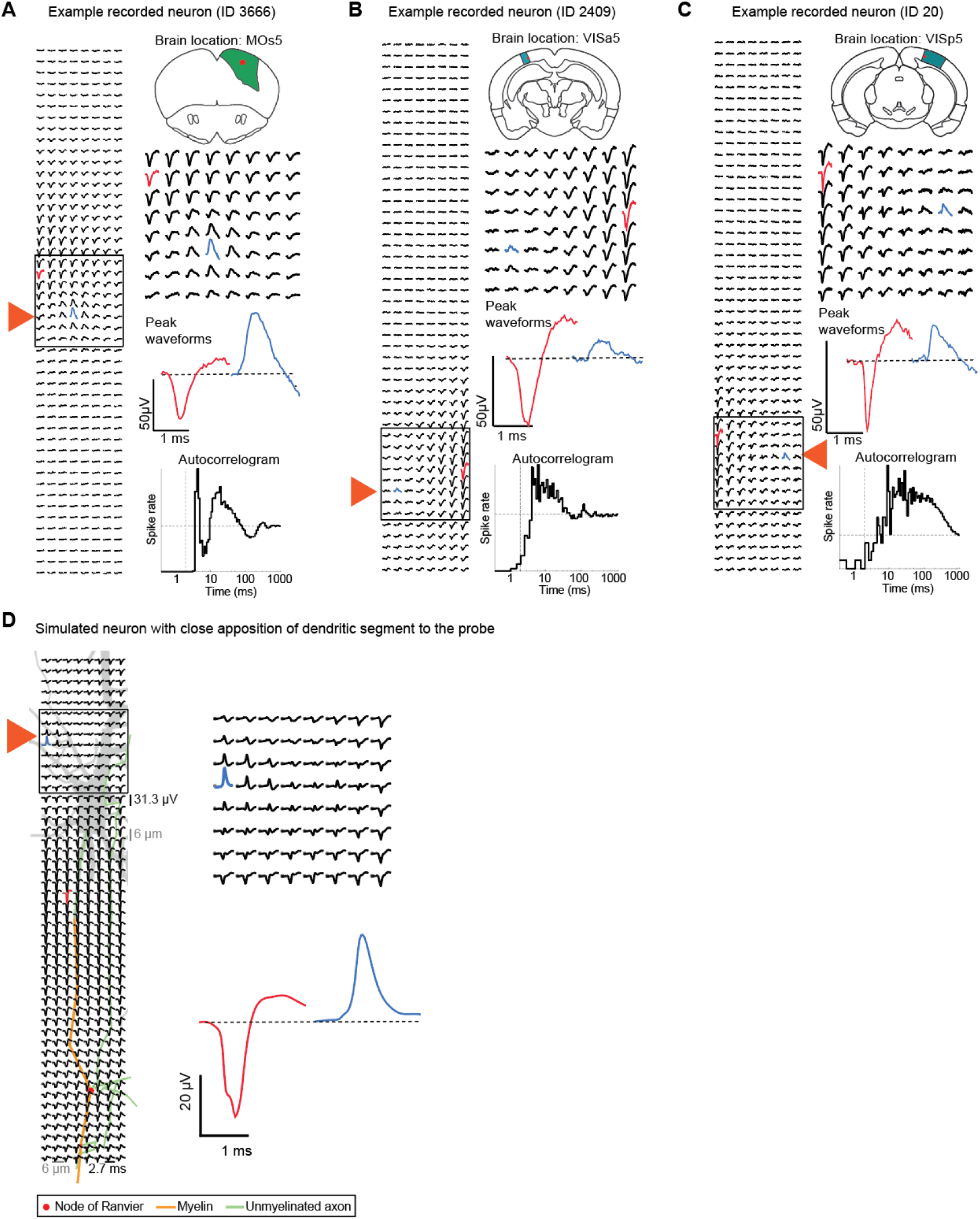
Putative observation of electrical fields from dendritic branches. **A,** Example neuron for which the common extracellular action potential pattern of cortical somatic recordings (negative-going spike, large spatial footprint) is accompanied by a very small region with positive-going potentials, indicated by an orange triangle. Figure format as in Supp. Fig. S15A. (https://npultra.steinmetzlab.net/?selected=3666 to browse this neuron). MOs5: secondary motor cortex, layer 5. **B**, Another example neuron as in A (https://npultra.steinmetzlab.net/?selected=3666). VISa5: anterior visual cortical area, layer 5. **C**, A third example neuron as in A (https://npultra.steinmetzlab.net/?selected=20). VISp5: primary visual cortex, layer 5. **D**, A simulated neuron with similar features to the example real neurons. The grey silhouette shows the spatial position of the simulated neuron relative to the probe, with a dendritic segment in close apposition to the probe near the orange triangle.

**Supplemental Figure S17.**
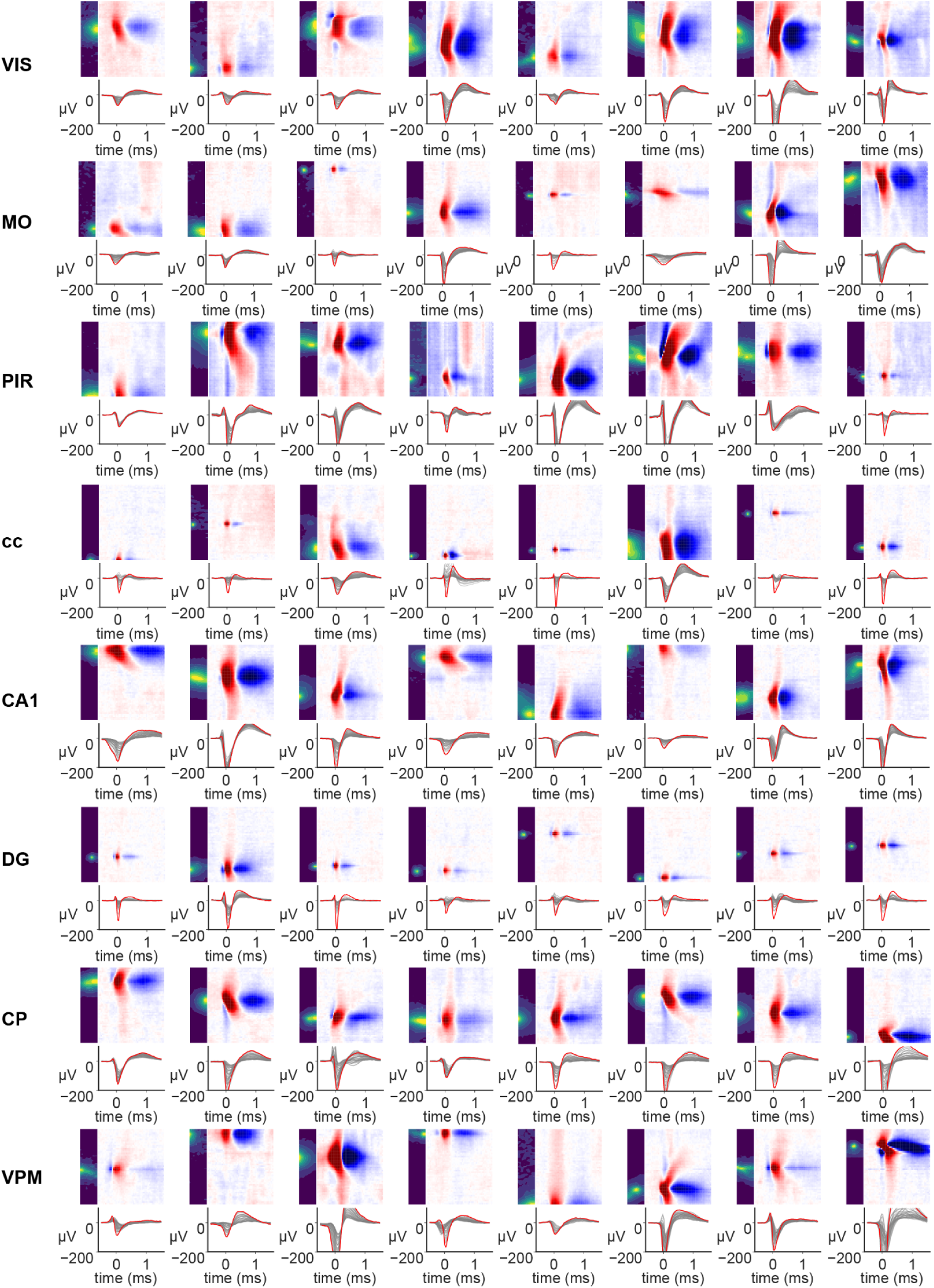
Eight example neurons recorded in each of eight example brain regions. Each panel depicts the spatial, spatiotemporal, and temporal waveform of a single neuron. VIS: visual areas; MO: Somatomotor areas; PIR: Piriform area; cc: corpus callosum; CA1: hippocampal subfield CA1; DG: Dentate gyrus; CP: Caudoputamen; VPM: ventral posteromedial nucleus of the thalamus.

**Supplemental Figure S18.**
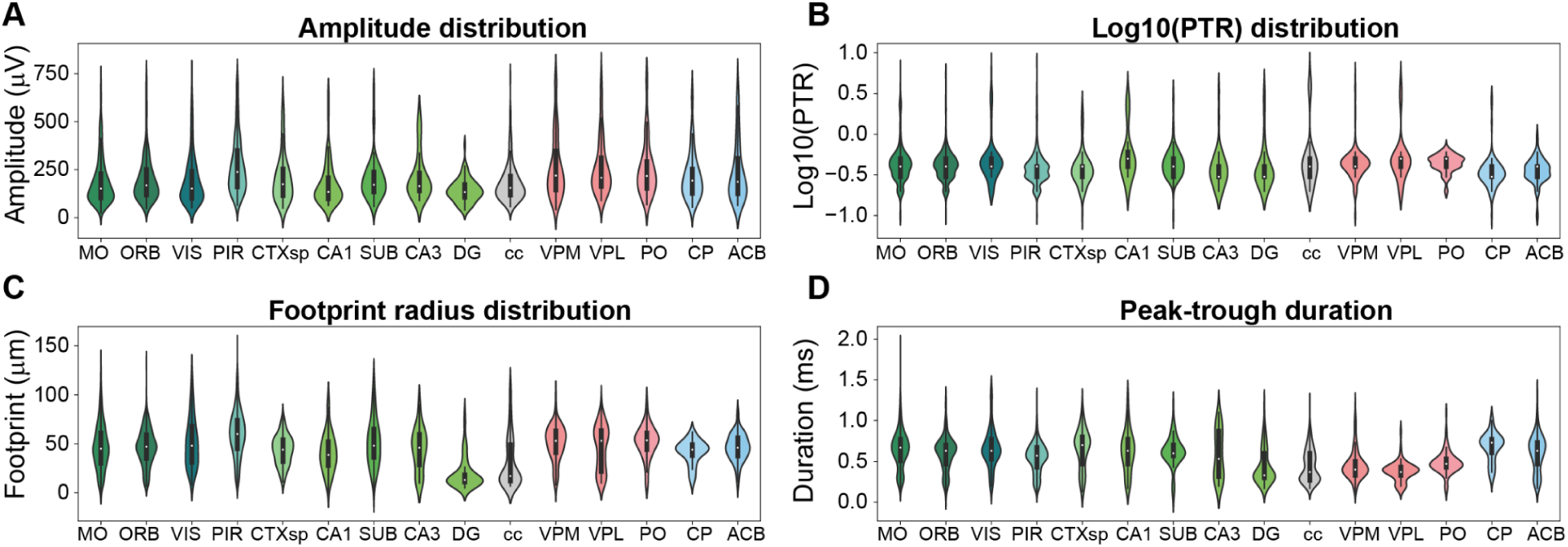
Features extracted from some top recorded brain regions. **A,** Absolute peak amplitude distribution. **B**, Spatial footprint radius distribution. Note that footprint radius distribution of DG and cc waveforms are smaller than the other brain regions. **C**, Log10 of peak-to-trough ratio (PTR) distribution. Note that CP neurons tend to have smaller PTR index. **D**, Peak-to-trough duration (ms) distribution. Note that thalamic regions VPM, VPL, and PO have smaller peak-to-trough duration than other brain regions. MO: Somatomotor areas; ORB: orbital area; VIS: visual areas; PIR: Piriform area; CTXsp: cortical subplate; SUB: Subiculum; DG: Dentate gyrus; cc: corpus callosum; VPM: ventral posteromedial nucleus of the thalamus; VPL: Ventral posterolateral nucleus of the thalamus; PO: Posterior complex of the thalamus; CP: Caudoputamen; ACB: Nucleus accumbens.

**Supplemental Figure S19.**
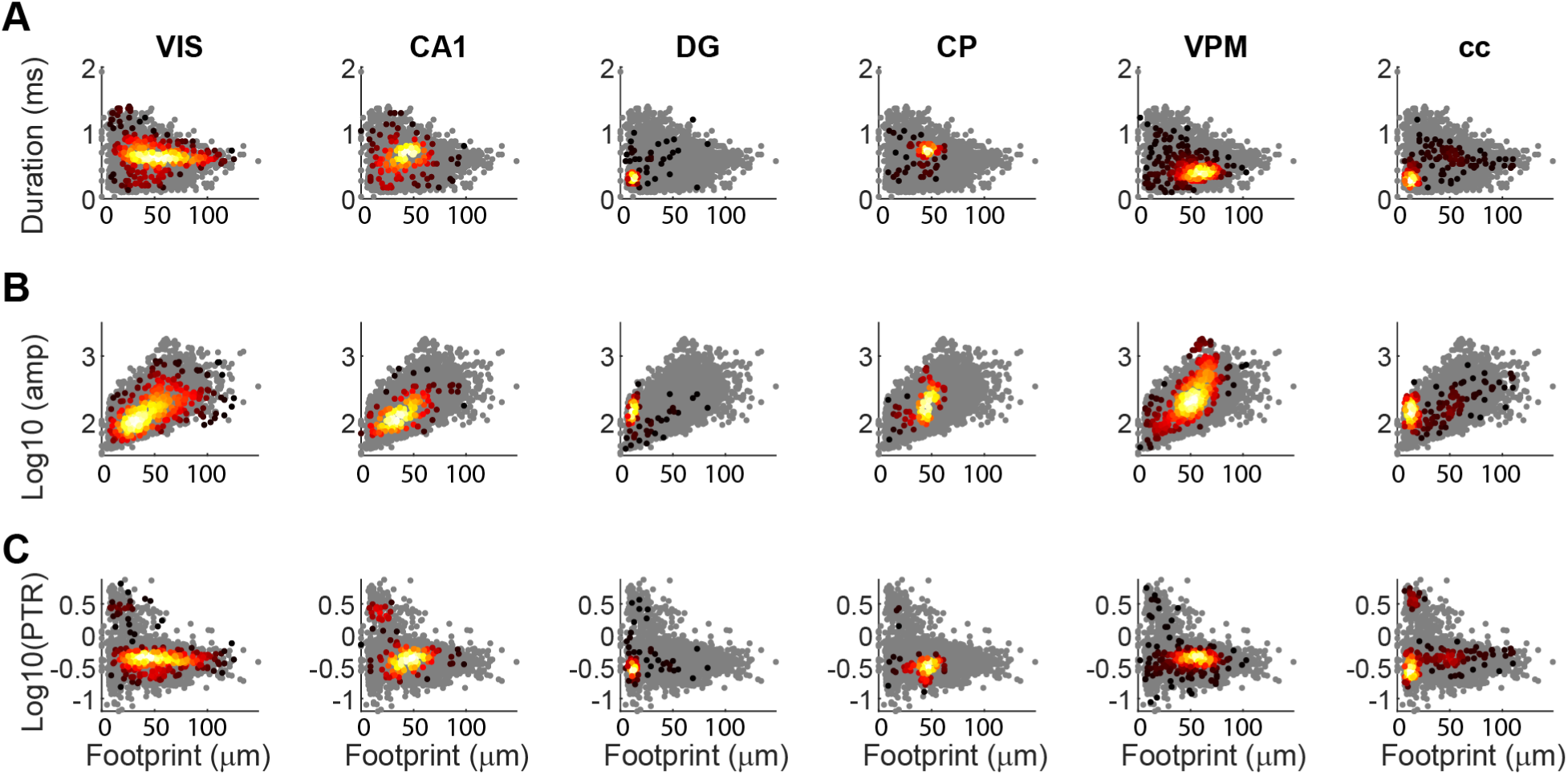
Relationships between features extracted from some top recorded brain regions. **A**, Waveform duration vs spatial footprint. Waveform duration is defined as duration between peak and trough. Dots colored black to yellow are units from this region; gray background dots are the distribution of units from all areas for reference. **B**, Log10(amplitude) vs spatial footprint. **C**, Waveform spatial footprint radius vs log10(peak-to-trough ratio). Values of log10(PTR) above 0 indicate absolute peak amplitude are higher than absolute trough amplitude, Values of log10(PTR) below 0 indicate absolute peak amplitude are lower than absolute trough amplitude.

**Supplemental Figure S20.**
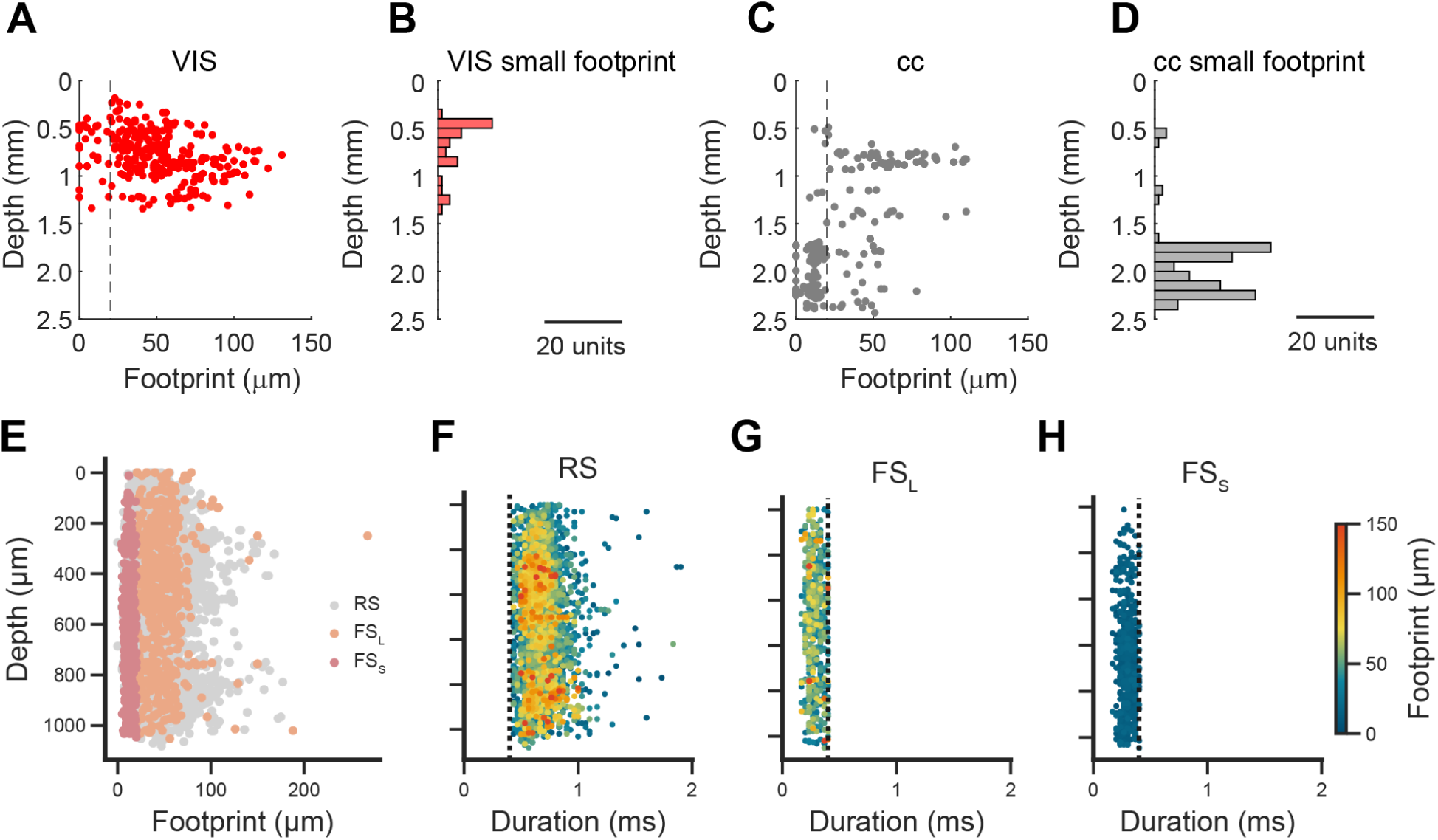
Depth separation of units with small footprint in the corpus callosum and visual cortex. **A**, Footprint and depth relationship of units in the visual cortex, from the brain-wide dataset (from Fig. 4). Depth 0 indicates the surface of the cortex. Note that a number of small footprint units are intermingled with larger footprint units across cortical depths. **B**, Depth histogram of units in the visual cortex with footprint smaller than 20 µm. **C**, Footprint and depth relationship of units in the corpus callosum. **D**, Depth histogram of units in the corpus callosum with footprint smaller than 20 µm. The cc can be reached at different depths depending on the location and angle of insertion, which varied across recordings. Note that the depth estimations are calculated from incremental manipulator advancement, which could result in a small overestimation of the true depth due to tissue compression. The border between VIS and cc was estimated with electrophysiological features (excluding footprint) in combination with histology, and was independent of the depth estimation taken from the micromanipulator readings. **E**, Distribution of RS, FS_L_, and FS_S_ units of the visual cortex by spatial footprint and depth from the cortical surface, from the visual cortex dataset (from Figs. 7,8). **F**-**H**, Depth by duration footprint distributions for each of the cell types in **E**, colored by footprint. Vertical dashed line indicates the threshold between RS and FS units.

**Supplemental Figure S21.**
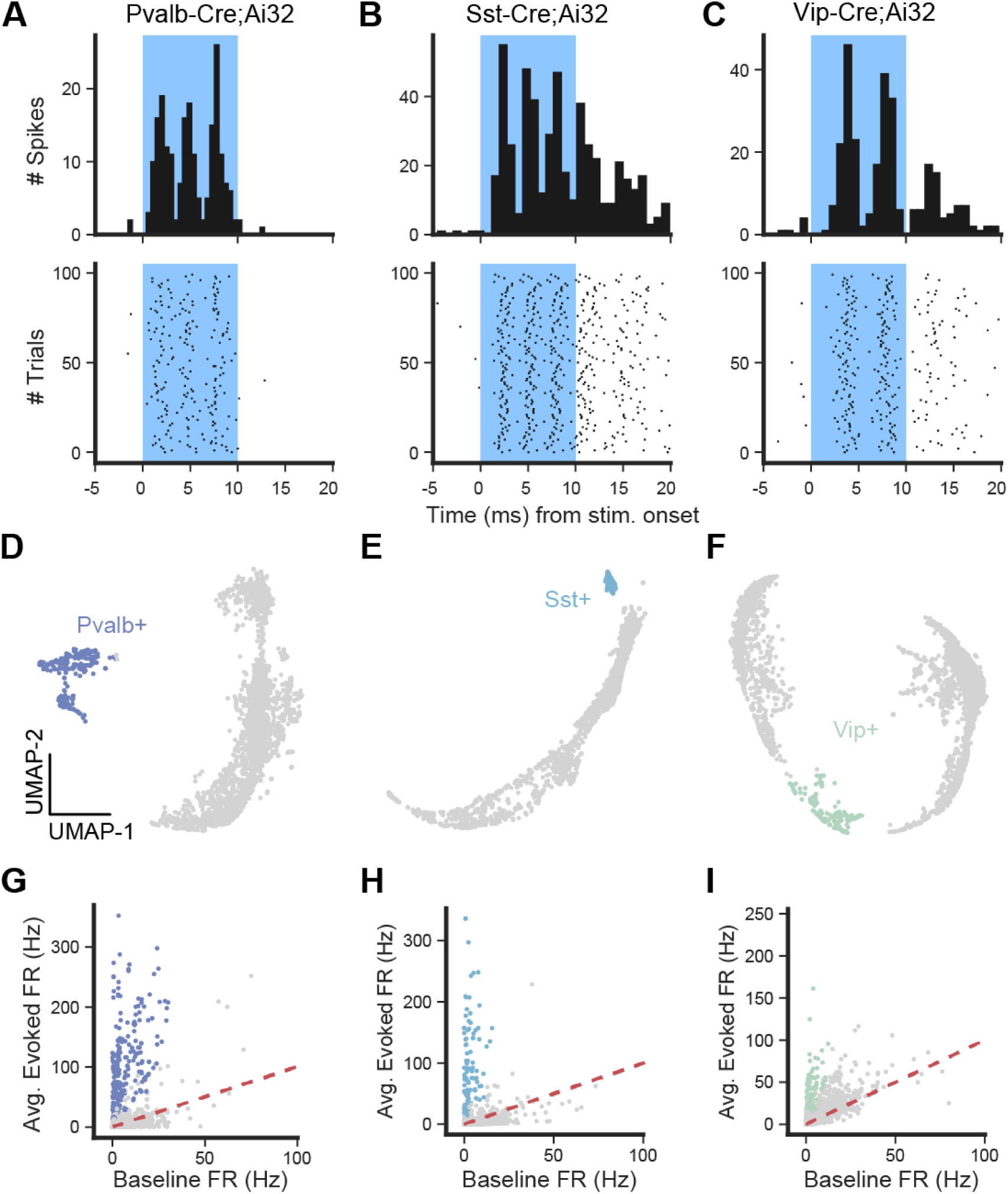
Identification of optotagged units. Optotagged units were identified by photo-evoked changes in firing rate in average PSTHs from both 10ms (square-wave) and 1s (cosine ramp) pulses. **A**-**C**, PSTHs (top) and spike rasters per photostimulation trial (bottom) in three example optotagged units from each transgenic mouse line. **D**-**F**, 2-dimensional UMAP representations of concatenated firing rate responses (see Methods) for all trials and stimulation-protocols for each transgenic mouse line. Optotagged units were found via unsupervised clustering (DBSCAN). **G**-**H**, Scatter plots showing baseline firing rate against the average evoked firing for each transgenic mouse line. Optotagged units found via unsupervised clustering (**D**-**F**) are shown in color, confirming that the clustering successfully identified units that were strongly modulated by light delivery.

**Supplemental Figure S22.**
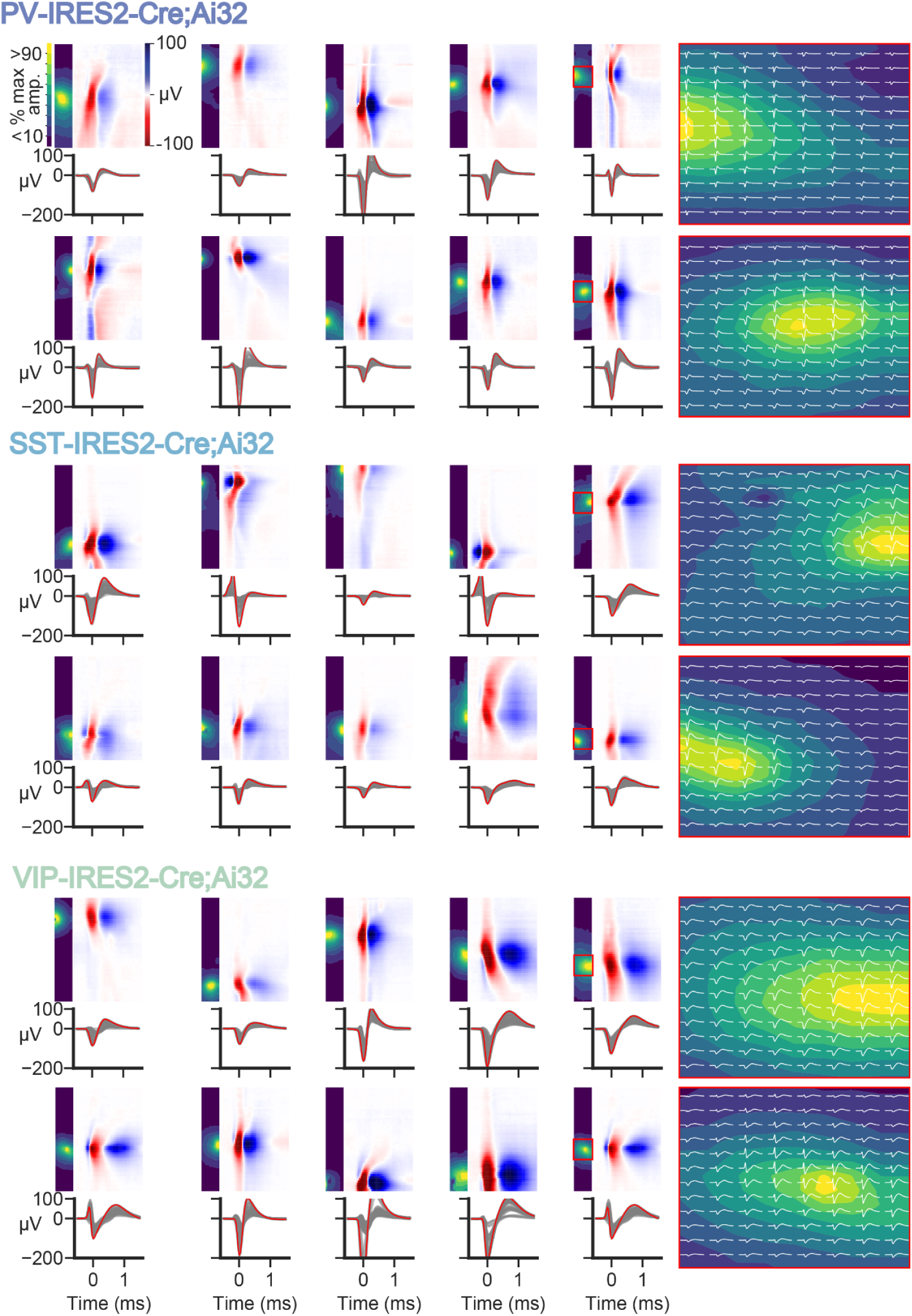
Diversity of extracellular waveforms across transgenically-defined inhibitory types. Ten example units demonstrating the variability in extracellular waveforms within the three examined inhibitory neuron subclasses. Plotting conventions as in **Fig. 1G**. Heatmaps at right are insets (red box) of the right-most unit displayed in each row, overlaid with waveforms recorded on the 12 × 8 channels surrounding the peak amplitude channel.

**Supplemental Figure S23.**
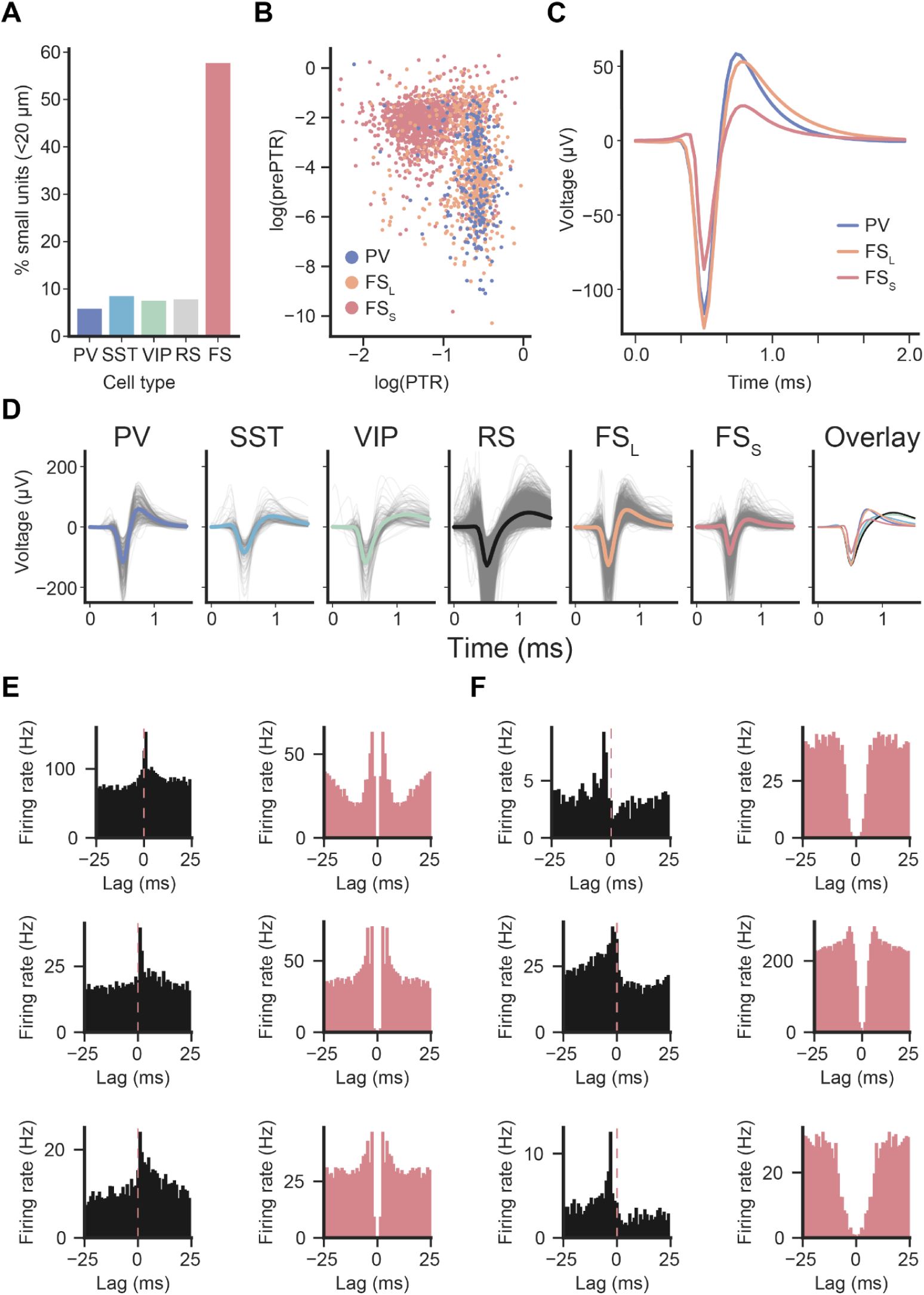
Waveform characteristics of each cell class, and inferred synaptic connections of FS_S_ units. **A**, Percent small footprint units within each cell class category. **B**, Scatter plot showing the log-scaled prePTR and PTR values for units in each of PV, FS_L_, and FS_S_ cell class, showing that FS_S_ units differ not only in spatial footprint but also in PTR. **C**, Comparison of grand-average waveforms from FS_L_, FS_S_, and PV units. **D**, Grand-average mean waveforms for each identified cell class, overlaid with each other in the right-most panel. Gray traces are mean waveforms from individual units. **E**, Putative excitatory CCGs (left column) and ACGs (right column) from identified FS_S_ units. **F**, Putative inhibitory CCGs (left column) and ACGs (right column) from identified FS_S_ units.

**Supplemental Figure S24.**
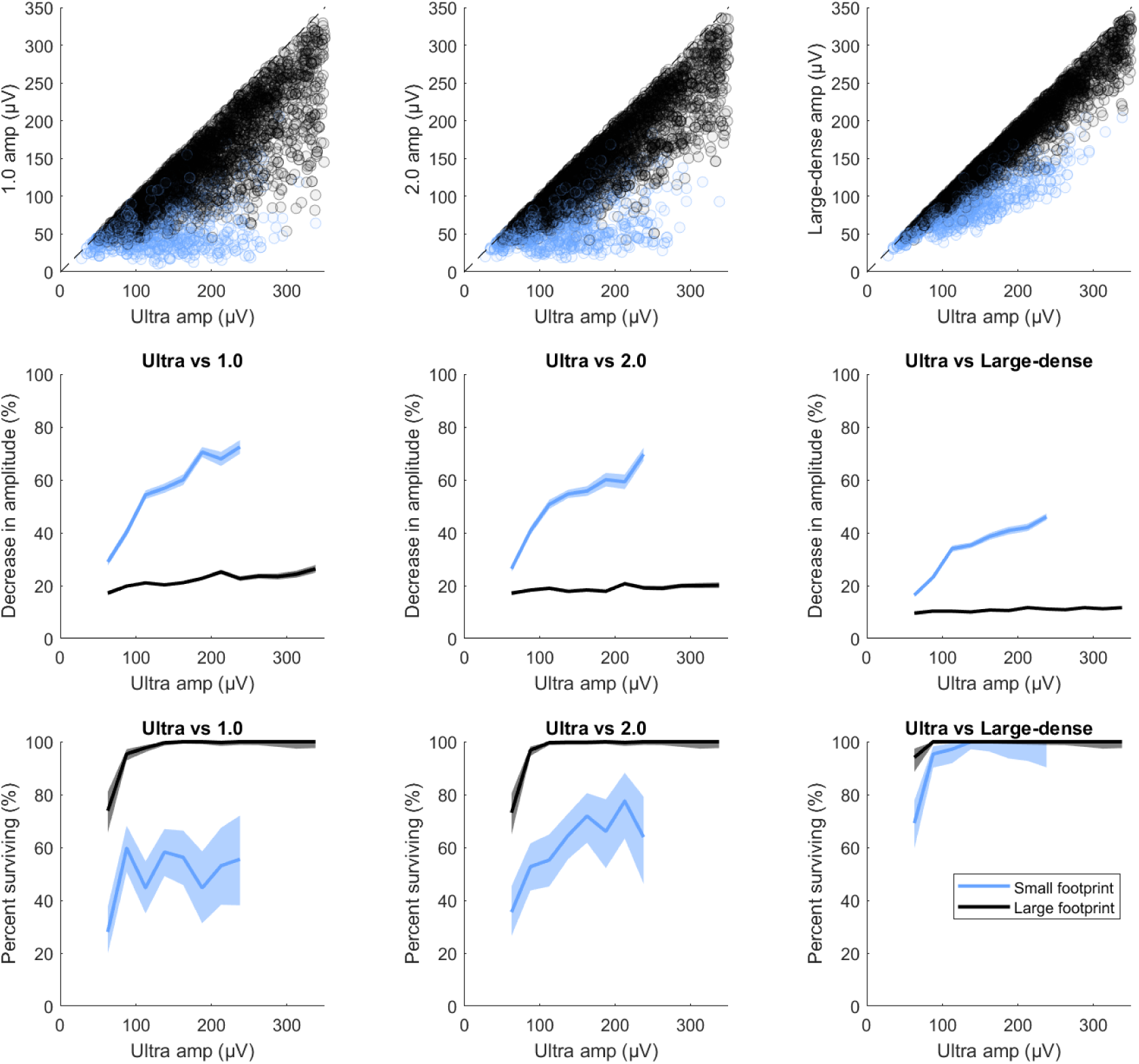
Units with small footprint are preferentially more difficult to detect on probes with lower recording site density. Top row, the amplitude of each neuron in the brain-wide acute mouse dataset (**Fig. 5**; n=4666) as measured in the original recording (x-axis) and on each spatially resampled dataset (y-axis; for 1.0-like resampling, 2.0-like resampling, and ‘large-dense’ resampling (**Fig. 2E**) from left to right). Small-footprint units are plotted in blue. Middle row, the percentage decrease in amplitude between NP Ultra and each spatially resampled dataset as a function of the original amplitude of the unit in the NP Ultra recording. Small-footprint units (blue) decrease more than large-footprint units (black), especially for resampling at lower density (1.0-like, left panel) and for higher amplitude units. For example, small-footprint units decrease their amplitude by 56.4 ± 0.8% (mean ± s.e.m.) when spatially resampled in the NP 1.0-like pattern, but large-footprint units only decrease by 23.6 ± 0.3%. Bottom row, Applying a cutoff of 40 µV minimum amplitude to approximately estimate the percentage of units that would ‘survive’ the spatial resampling (i.e. still be spike sortable), we find that nearly all large-footprint units of sufficiently large amplitude would remain above the amplitude threshold and thus survive to be recorded with the lower density probe, but approximately half of small-footprint units would instead fail to survive in a lower density recording.

**Supplemental Figure S25.**
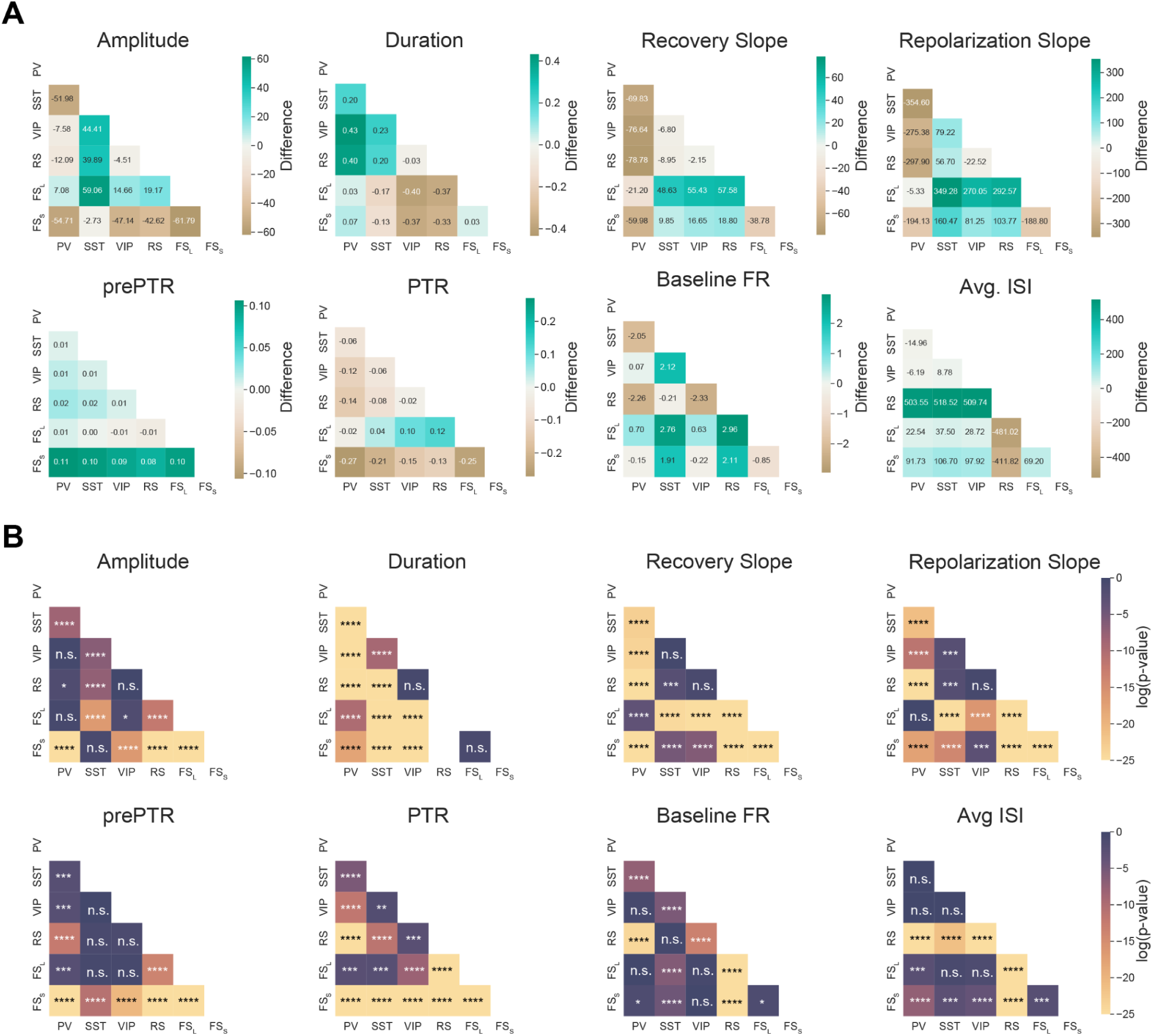
Statistics of 1D feature comparisons between cell classes. **A**, Pairwise median differences of extracted 1D single-channel waveform features. Values are represented as the difference in distribution medians (row-column) between cell classes for each feature. **B**, Pairwise statistics (Mann-Whitney U test, Benjamini-Hochberg multiple comparisons corrected) between each cell class for each 1D single-channel feature, log scaled. Blank spaces in Duration heatmap result from no overlap between RS and FS distributions.

**Supplemental Figure S26.**
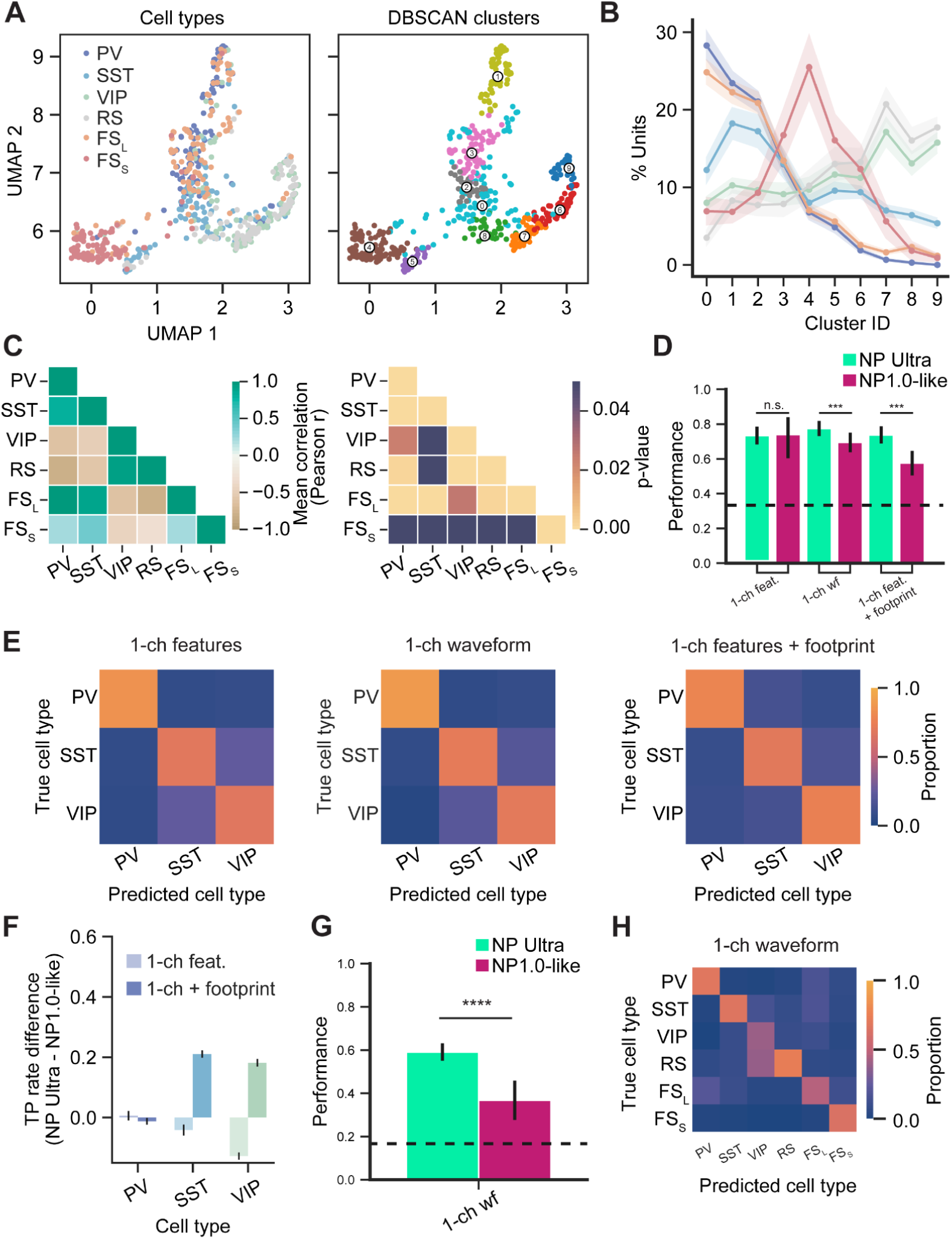
Further analyses on cell class discrimination. **A**, Example UMAP projections of an equally subsampled set of units from each cell class, overlayed with cell class (left) and cluster ID (right). **B**, Percentage of units found within each cluster through iterative application of UMAP on a subsampled dataset. Clustering was performed via DBSCAN and constrained to a maximum of 10 clusters. Error represents the 95% confidence interval about the mean. **C**, Average correlation of units found across all clusters in **B** (left) and the p-value of those correlation (right). **D**, Accuracy scores for optotagged-only classification. Bars represent mean scores ± SEM. **E**, Confusion matrices for optotagged only classification for each feature set. **F**, True Positive (TP) differences between NP Ultra and NP 1.0-like classification for 1-ch and 1-ch+footprint feature sets. **G**, Accuracy of 1-ch waveform classification when all units are used as samples. **H**, Confusion matrix of the classification performed in **G**.

## References

Allen WE, Chen MZ, Pichamoorthy N, Tien RH, Pachitariu M, Luo L, Deisseroth K (2019) Thirst regulates motivated behavior through modulation of brainwide neural population dynamics. Science 364.

Barthó P (2021) Extracellular recording of axonal spikes in the visual cortex. J Physiol 599:2131–2131.

Barthó P, Hirase H, Monconduit L, Zugaro M, Harris KD, Buzsaki G (2004) Characterization of neocortical principal cells and interneurons by network interactions and extracellular features. J Neurophysiol 92:600–608.

Barthó P, Slézia A, Mátyás F, Faradzs-Zade L, Ulbert I, Harris KD, Acsády L (2014) Ongoing Network State Controls the Length of Sleep Spindles via Inhibitory Activity. Neuron 82:1367–1379.

Beau M et al. (2024) A deep-learning strategy to identify cell types across species from high-density extracellular recordings. Available at: http://biorxiv.org/lookup/doi/10.1101/2024.01.30.577845 [Accessed March 26, 2024].

Beaulieu-Laroche L, Brown NJ, Hansen M, Toloza EHS, Sharma J, Williams ZM, Frosch MP, Cosgrove GR, Cash SS, Harnett MT (2021) Allometric rules for mammalian cortical layer 5 neuron biophysics. Nature 600:274–278.

Boussard J, Varol E, Lee HD, Dethe N, Paninski L (2021) Three-dimensional spike localization and improved motion correction for Neuropixels recordings. Neuroscience. Available at: http://biorxiv.org/lookup/doi/10.1101/2021.11.05.467503 [Accessed May 18, 2023].

Boussard J, Windolf C, Hurwitz C, Lee HD, Yu H, Winter O, Paninski L (2023) DARTsort: A modular drift tracking spike sorter for high-density multi-electrode probes. :2023.08.11.553023 Available at: https://www.biorxiv.org/content/10.1101/2023.08.11.553023v1 [Accessed August 15, 2023].

Buccino AP, Kordovan M, Ness TV, Merkt B, Häfliger PD, Fyhn M, Cauwenberghs G, Rotter S, Einevoll GT (2018) Combining biophysical modeling and deep learning for multielectrode array neuron localization and classification. J Neurophysiol 120:1212–1232.

Buzsaki G (2006) Rhythms of the Brain. Oxford university press.

Buzsáki G, Anastassiou CA, Koch C (2012) The origin of extracellular fields and currents—EEG, ECoG, LFP and spikes. Nat Rev Neurosci 13:407–420.

Buzsáki G, Kandel A (1998) Somadendritic Backpropagation of Action Potentials in Cortical Pyramidal Cells of the Awake Rat. J Neurophysiol 79:1587–1591.

Carnevale NT, Hines ML (2006) The NEURON Book. Cambridge: Cambridge University Press. Available at: https://www.cambridge.org/core/books/neuron-book/7C8D9BD861D288E658BEB652F593F273 [Accessed April 2, 2024].

Chen S, Liu Y, Wang Z, Colonell J, Liu LD, Hou H, Tien N-W, Wang T, Harris T, Druckmann S (2023) Brain-wide neural activity underlying memory-guided movement. BioRxiv:2023–03.

Chen S, Svoboda K (2020) “Uniclear” water-based brain clearing for light sheet imaging of electrode tracks. Available at: https://www.protocols.io/view/39-uniclear-39-water-based-brain-clearing-for-lig-zndf5a6 [Accessed August 22, 2023].

Chung JE, Sellers KK, Leonard MK, Gwilliams L, Xu D, Dougherty ME, Kharazia V, Metzger SL, Welkenhuysen M, Dutta B, Chang EF (2022) High-density single-unit human cortical recordings using the Neuropixels probe. Neuron 110:2409–2421.e3.

Cooper GF, Robson JG, Waldron I (1969) The action potentials recorded from undamaged nerve fibres with micro-electrodes. J Physiol 200:9P–11P.

Crawford E, Pineau J (2019) Spatially Invariant Unsupervised Object Detection with Convolutional Neural Networks. Proc AAAI Conf Artif Intell 33:3412–3420.

Deligkaris K, Bullmann T, Frey U (2016) Extracellularly Recorded Somatic and Neuritic Signal Shapes and Classification Algorithms for High-Density Microelectrode Array Electrophysiology. Front Neurosci 10 Available at: https://www.frontiersin.org/journals/neuroscience/articles/10.3389/fnins.2016.00421/full [Accessed March 27, 2024].

Fiáth R, Raducanu BC, Musa S, Andrei A, Lopez CM, Welkenhuysen M, Ruther P, Aarts A, Ulbert I (2018) Fine-scale mapping of cortical laminar activity during sleep slow oscillations using high-density linear silicon probes. J Neurosci Methods:0–1.

Gardner RJ, Hermansen E, Pachitariu M, Burak Y, Baas NA, Dunn BA, Moser M-B, Moser EI (2022) Toroidal topology of population activity in grid cells. Nature 602:123–128.

Gilman JP, Medalla M, Luebke JI (2017) Area-Specific Features of Pyramidal Neurons—a Comparative Study in Mouse and Rhesus Monkey. Cereb Cortex 27:2078–2094.

Gold C, Henze DA, Koch C, Buzsáki G (2006) On the origin of the extracellular action potential waveform: A modeling study. J Neurophysiol 95:3113–3128.

Goldberg JH, Fee MS (2012) A cortical motor nucleus drives the basal ganglia-recipient thalamus in singing birds. Nat Neurosci 15:620–627.

Gouwens NW et al. (2020) Integrated Morphoelectric and Transcriptomic Classification of Cortical GABAergic Cells. Cell 183:935–953.e19.

Hagen E, Næss S, Ness TV, Einevoll GT (2018) Multimodal Modeling of Neural Network Activity: Computing LFP, ECoG, EEG, and MEG Signals With LFPy 2.0. Front Neuroinformatics 12 Available at: https://www.frontiersin.org/articles/10.3389/fninf.2018.00092 [Accessed April 2, 2024].

Hallermann S, de Kock CPJ, Stuart GJ, Kole MHP (2012) State and location dependence of action potential metabolic cost in cortical pyramidal neurons. Nat Neurosci 15:1007–1014.

Halnes G, Ness T, Naess S, Hagen E, Pettersen K, Einevoll G (2024) Electric brain signals. Cambridge University Press.

Hay E, Hill S, Schürmann F, Markram H, Segev I (2011) Models of Neocortical Layer 5b Pyramidal Cells Capturing a Wide Range of Dendritic and Perisomatic Active Properties. PLOS Comput Biol 7:e1002107.

Herculano-Houzel S, Manger PR, Kaas JH (2014) Brain scaling in mammalian evolution as a consequence of concerted and mosaic changes in numbers of neurons and average neuronal cell size. Front Neuroanat 8 Available at: https://www.frontiersin.org/articles/10.3389/fnana.2014.00077 [Accessed May 17, 2023].

Hilgen G, Sorbaro M, Pirmoradian S, Muthmann J-O, Kepiro IE, Ullo S, Ramirez CJ, Encinas AP, Maccione A, Berdondini L (2017) Unsupervised spike sorting for large-scale, high-density multielectrode arrays. Cell Rep 18:2521–2532.

International Brain Laboratory et al. (2023a) Reproducibility of in-vivo electrophysiological measurements in mice. :2022.05.09.491042 Available at: https://www.biorxiv.org/content/10.1101/2022.05.09.491042v4 [Accessed August 18, 2023].

International Brain Laboratory, Benson B, Benson J, Birman D, Bonacchi N, Carandini M, Catarino JA, Chapuis GA, Churchland AK, Dan Y (2023b) A Brain-Wide Map of Neural Activity during Complex Behaviour. bioRxiv:2023–07.

Jia X, Siegle J, Bennett C, Gale S, Denman D, Koch C, Olsen S (2019) High-density extracellular probes reveal dendritic backpropagation and facilitate neuron classification. J Neurophysiol.

Jia X, Siegle JH, Durand S, Heller G, Ramirez TK, Koch C, Olsen SR (2022) Multi-regional module-based signal transmission in mouse visual cortex. Neuron 110:1585–1598.e9.

Jun JJ et al. (2017a) Fully integrated silicon probes for high-density recording of neural activity. Nature 551:232–236.

Jun JJ et al. (2017b) Fully integrated silicon probes for high-density recording of neural activity. Nature 551:232–236.

Katzner S, Nauhaus I, Benucci A, Bonin V, Ringach DL, Carandini M (2009) Local Origin of Field Potentials in Visual Cortex. Neuron 61:35–41.

Kiani R, Esteky H, Mirpour K, Tanaka K (2007) Object category structure in response patterns of neuronal population in monkey inferior temporal cortex. J Neurophysiol 97:4296–4309.

Kingma DP, Welling M (2022) Auto-Encoding Variational Bayes. Available at: http://arxiv.org/abs/1312.6114 [Accessed August 18, 2023].

Kvitsiani D, Ranade S, Hangya B, Taniguchi H, Huang JZ, Kepecs A (2013) Distinct behavioural and network correlates of two interneuron types in prefrontal cortex. Nature 498:363–366.

Lee EK, Balasubramanian H, Tsolias A, Anakwe SU, Medalla M, Shenoy KV, Chandrasekaran C (2021) Non-linear dimensionality reduction on extracellular waveforms reveals cell type diversity in premotor cortex. Elife 10:e67490.

Lee EK, Gül AE, Heller G, Lakunina A, Jaramillo S, Przytycki PF, Chandrasekaran C (2024) PhysMAP - interpretable in vivo neuronal cell type identification using multi-modal analysis of electrophysiological data. Available at: http://biorxiv.org/lookup/doi/10.1101/2024.02.28.582461 [Accessed March 26, 2024].

Lee J, Mitelut C, Shokri H, Kinsella I, Dethe N, Wu S, Li K, Reyes EB, Turcu D, Batty E, Kim YJ, Brackbill N, Kling A, Goetz G, Chichilnisky EJ, Carlson D, Paninski L (2020) YASS: Yet Another Spike Sorter applied to large-scale multi-electrode array recordings in primate retina. Neuroscience. Available at: http://biorxiv.org/lookup/doi/10.1101/2020.03.18.997924 [Accessed May 17, 2023].

Lima SQ, Hromádka T, Znamenskiy P, Zador AM (2009) PINP: A New Method of Tagging Neuronal Populations for Identification during In Vivo Electrophysiological Recording Nitabach MN, ed. PLoS ONE 4:e6099.

López CM, Welkenhuysen M, Musa S, Eberle W, Bartic C, Puers R, Gielen G (2012) Towards a noise prediction model for in vivo neural recording. In: 2012 Annual International Conference of the IEEE Engineering in Medicine and Biology Society, pp 759–762. IEEE.

Lopez CM, Welkenhuysen M, Musa S, Eberle W, Bartic C, Puers R, Gielen G (2012) Towards a noise prediction model for in vivo neural recording. Proc Annu Int Conf IEEE Eng Med Biol Soc EMBS:759–762.

Malzer C, Baum M (2019) A Hybrid Approach To Hierarchical Density-based Cluster Selection. Available at: https://arxiv.org/abs/1911.02282 [Accessed May 17, 2023].

Mazer, Jamie (2013) Pype2. Available at: https://github.com/mazerj/pype3.

McColgan T, Liu J, Kuokkanen PT, Carr CE, Wagner H, Kempter R (2017) Dipolar extracellular potentials generated by axonal projections van Rossum MC, ed. eLife 6:e26106.

McCormick DA, Connors BW, Lighthall JW, Prince DA (1985) Comparative electrophysiology of pyramidal and sparsely spiny stellate neurons of the neocortex. J Neurophysiol 54:782–806.

McHugh SB, Lopes-dos-Santos V, Gava GP, Hartwich K, Tam SKE, Bannerman DM, Dupret D (2022) Adult-born dentate granule cells promote hippocampal population sparsity. Nat Neurosci 25:1481–1491.

McLelland D, Baker PM, Ahmed B, Kohn A, Bair W (2015) Mechanisms for Rapid Adaptive Control of Motion Processing in Macaque Visual Cortex. J Neurosci 35:10268–10280.

Merrill EG, Wall PD, Yaksh TL (1978) Properties of two unmyelinated fibre tracts of the central nervous system: lateral Lissauer tract, and parallel fibres of the cerebellum. J Physiol 284:127–145.

Metzen MG, Chacron MJ (2021) Population Coding of Natural Electrosensory Stimuli by Midbrain Neurons. J Neurosci 41:3822–3841.

Mitchell JF, Sundberg KA, Reynolds JH (2007) Differential Attention-Dependent Response Modulation across Cell Classes in Macaque Visual Area V4. Neuron 55:131–141.

Musall S, Kaufman MT, Juavinett AL, Gluf S, Churchland AK (2019) Single-trial neural dynamics are dominated by richly varied movements. Nat Neurosci 22:1677–1686.

Neubrandt M, Oláh VJ, Brunner J, Marosi EL, Soltesz I, Szabadics J (2018) Single Bursts of Individual Granule Cells Functionally Rearrange Feedforward Inhibition. J Neurosci 38:1711–1724.

Niell CM, Stryker MP (2008) Highly Selective Receptive Fields in Mouse Visual Cortex. J Neurosci 28:7520–7536.

Norimoto H, Fenk LA, Li H-H, Tosches MA, Gallego-Flores T, Hain D, Reiter S, Kobayashi R, Macias A, Arends A, Klinkmann M, Laurent G (2020) A claustrum in reptiles and its role in slow-wave sleep. Nature 578:413–418.

Nunez PL, Srinivasan R (2006) Electric fields of the brain: the neurophysics of EEG. Oxford University Press, USA.

Paulk AC, Kfir Y, Khanna AR, Mustroph ML, Trautmann EM, Soper DJ, Stavisky SD, Welkenhuysen M, Dutta B, Shenoy KV, Hochberg LR, Richardson RM, Williams ZM, Cash SS (2022) Large-scale neural recordings with single neuron resolution using Neuropixels probes in human cortex. Nat Neurosci 25:252–263.

Pedroni A, Minh DD, Mallamaci A, Cherubini E (2014) Electrophysiological characterization of granule cells in the dentate gyrus immediately after birth. Front Cell Neurosci 8 Available at: http://journal.frontiersin.org/article/10.3389/fncel.2014.00044/abstract [Accessed July 17, 2023].

Reifenstein ET, Ebbesen CL, Tang Q, Brecht M, Schreiber S, Kempter R (2016) Cell-Type Specific Phase Precession in Layer II of the Medial Entorhinal Cortex. J Neurosci 36:2283–2288.

Rey HG, Pedreira C, Quian Quiroga R (2015) Past, present and future of spike sorting techniques. Brain Res Bull 119:106–117.

Robbins A, Fox S, Holmes G, Scott R, Barry J (2013) Short duration waveforms recorded extracellularly from freely moving rats are representative of axonal activity. Front Neural Circuits 7 Available at: https://www.frontiersin.org/articles/10.3389/fncir.2013.00181 [Accessed August 17, 2023].

Roux L, Stark E, Sjulson L, Buzsáki G (2014) In vivo optogenetic identification and manipulation of GABAergic interneuron subtypes. Curr Opin Neurobiol 26:88–95.

Schröder S, Steinmetz NA, Krumin M, Pachitariu M, Rizzi M, Lagnado L, Harris KD, Carandini M (2020) Arousal Modulates Retinal Output. Neuron 107:487–495.e9.

Senzai Y, Buzsáki G (2017) Physiological Properties and Behavioral Correlates of Hippocampal Granule Cells and Mossy Cells. Neuron 93:691–704.e5.

Shoham S, O’Connor DH, Segev R (2006) How silent is the brain: is there a “dark matter” problem in neuroscience? J Comp Physiol A 192:777–784.

Sibille J, Gehr C, Benichov JI, Balasubramanian H, Teh KL, Lupashina T, Vallentin D, Kremkow J (2022) High-density electrode recordings reveal strong and specific connections between retinal ganglion cells and midbrain neurons. Nat Commun 13:5218.

Siegle JH, Jia X, Durand S, Gale S, Bennett C, Graddis N, Heller G, Ramirez TK, Choi H, Luviano JA (2021) Survey of spiking in the mouse visual system reveals functional hierarchy. Nature 592:86–92.

Someck S, Levi A, Sloin HE, Spivak L, Gattegno R, Stark E (2023) Positive and biphasic extracellular waveforms correspond to return currents and axonal spikes. Commun Biol 6:1–15.

Steinmetz NA et al. (2021) Neuropixels 2.0: A miniaturized high-density probe for stable, long-term brain recordings. Science 372 Available at: https://science.sciencemag.org/content/372/6539/eabf4588 [Accessed July 20, 2021].

Steinmetz NA, Koch C, Harris KD, Carandini M (2018) Challenges and opportunities for large-scale electrophysiology with Neuropixels probes. Curr Opin Neurobiol 50:92–100.

Steinmetz NA, Zatka-Haas P, Carandini M, Harris KD (2019) Distributed coding of choice, action and engagement across the mouse brain. Nature 576:266–273.

Stringer C, Pachitariu M, Steinmetz N, Reddy CB, Carandini M, Harris KD (2019) Spontaneous behaviors drive multidimensional, brainwide activity. Science 364.

Stuart G, Schiller J, Sakmann B (1997) Action potential initiation and propagation in rat neocortical pyramidal neurons. J Physiol 505:617–632.

Sukhum KV, Shen J, Carlson BA (2018) Extreme Enlargement of the Cerebellum in a Clade of Teleost Fishes that Evolved a Novel Active Sensory System. Curr Biol 28:3857–3863.e3.

Sun SH, Almasi A, Yunzab M, Zehra S, Hicks DG, Kameneva T, Ibbotson MR, Meffin H (2021) Analysis of extracellular spike waveforms and associated receptive fields of neurons in cat primary visual cortex. J Physiol 599:2211–2238.

Swindale NV, Spacek MA (2014) Spike sorting for polytrodes: a divide and conquer approach. Front Syst Neurosci 8 Available at: http://journal.frontiersin.org/article/10.3389/fnsys.2014.00006/abstract [Accessed August 8, 2023].

Tosches MA, Yamawaki TM, Naumann RK, Jacobi AA, Tushev G, Laurent G (2018) Evolution of pallium, hippocampus, and cortical cell types revealed by single-cell transcriptomics in reptiles. Science 360:881–888.

Trautmann EM et al. (2023) Large-scale brain-wide neural recording in nonhuman primates.: 2023.02.01.526664 Available at: https://www.biorxiv.org/content/10.1101/2023.02.01.526664v2 [Accessed February 8, 2023].

Trautmann EM, Stavisky SD, Lahiri S, Ames KC, Kaufman MT, O’Shea DJ, Vyas S, Sun X, Ryu SI, Ganguli S (2019) Accurate estimation of neural population dynamics without spike sorting. Neuron 103:292–308.

Tymochko S, Munch E, Dunion J, Corbosiero K, Torn R (2020) Using persistent homology to quantify a diurnal cycle in hurricanes. Pattern Recognit Lett 133:137–143.

Vesuna S, Kauvar IV, Richman E, Gore F, Oskotsky T, Sava-Segal C, Luo L, Malenka RC, Henderson JM, Nuyujukian P (2020) Deep posteromedial cortical rhythm in dissociation. Nature 586:87–94.

Viswam V, Obien M, Frey U, Franke F, Hierlemann A (2017) Acquisition of bioelectrical signals with small electrodes. In: 2017 IEEE Biomedical Circuits and Systems Conference (BioCAS), pp 1–4. IEEE.

Windolf C et al. (2023) DREDge: robust motion correction for high-density extracellular recordings across species. :2023.10.24.563768 Available at: https://www.biorxiv.org/content/10.1101/2023.10.24.563768v1 [Accessed April 4, 2024].

Yamin HG, Stern EA, Cohen D (2013) Parallel Processing of Environmental Recognition and Locomotion in the Mouse Striatum. J Neurosci 33:473–484.

Yang Z, Zhao Q, Keefer E, Liu W (2009) Noise characterization, modeling, and reduction for in vivo neural recording. Adv Neural Inf Process Syst 22.

Yao Z et al. (2021) A taxonomy of transcriptomic cell types across the isocortex and hippocampal formation. Cell 184:3222–3241.e26.

Yu J, Hu H, Agmon A, Svoboda K (2019) Recruitment of GABAergic Interneurons in the Barrel Cortex during Active Tactile Behavior. Neuron 104:412–427.e4.

Zhang Y, He T, Boussard J, Windolf C, Winter O, Trautmann E, Roth N, Barrell H, Churchland M, Steinmetz NA (2024) Bypassing spike sorting: Density-based decoding using spike localization from dense multielectrode probes. Adv Neural Inf Process Syst 36 Available at: https://proceedings.neurips.cc/paper_files/paper/2023/hash/f499387f191d6be56e68966181095878-Abstract-Conference.html [Accessed April 9, 2024].

